# Limited parallelism in genetic adaptation to brackish water bodies in European sprat and Atlantic herring

**DOI:** 10.1101/2024.02.16.580647

**Authors:** Mats E. Pettersson, María Quintela, François Besnier, Qiaoling Deng, Florian Berg, Cecilie Kvamme, Dorte Bekkevold, Mai-Britt Mosbech, Ignas Bunikis, Roger Lille-Langøy, Iole Leonori, Andreas Wallberg, Kevin A. Glover, Leif Andersson

## Abstract

The European sprat is a small plankton-feeding clupeid present in the northeastern Atlantic Ocean, the Mediterranean Sea as well as in the brackish Baltic Sea and Black Sea. This species is the target of a major fishery and therefore an accurate characterization of its genetic population structure is crucial to delineate proper stock assessments that aid ensuring the fishery’s sustainability. Here we present (*i*) a draft genome assembly, (*ii*) pooled whole genome sequencing of 19 population samples covering most of the species’ distribution range, and (*iii*) the design and test of a SNP-chip resource and use this to validate the population structure inferred from pooled sequencing. These approaches revealed, using the populations sampled here, three major groups of European sprat: Oceanic, Coastal and Brackish with limited differentiation within groups even over wide geographical stretches. Genetic structure is largely driven by six large putative inversions that differentiate Oceanic and Brackish sprats, while Coastal populations display intermediate frequencies. Interestingly, populations from the Baltic and the Black Seas share similar frequencies of haplotypes at these putative inversions despite their distant geographic location. The closely related clupeids European sprat and Atlantic herring both show genetic adaptation to the brackish Baltic Sea, providing an opportunity to explore the extent of genetic parallelism. This analysis revealed limited parallelism in the form of three sharp signals of selection that overlapped between the two species and contained single genes such as *PRLRA*, which encodes the receptor for prolactin, a freshwater-adapting hormone in euryhaline species, and *THRB*, a receptor for thyroid hormones, important both for metabolic regulation and the development of red cone photoreceptors.

## Introduction

The European sprat (*Sprattus sprattus*), hereafter denoted sprat, is an economically important clupeid that constitutes the basis for a substantial fishery in Northern European waters. It inhabits an extensive range and is found all along the European Atlantic coastline, from Norway to Portugal, as well as in the Mediterranean Sea. Additionally, it has colonized several brackish environments, including the Baltic Sea, the Black Sea and Landvikvannet in southern Norway. The latter is a former freshwater lake that became brackish after the opening of a small canal in the 19th century (Whitehead 1985; Eggers et al. 2014; Berg et al. 2018). This colonization of multiple brackish environments provides an ideal situation to study the genetics of salinity-related adaptation, as it increases the power to infer specific salinity-related genetic differentiation.

Recent advances (Martinez-Barrio et al. 2016; Pettersson et al. 2019; Han et al. 2020)in understanding adaptive genetic differentiation in the closely related Atlantic herring (*Clupea harengus*; estimated split ∼10 MYA) (Jamsandekar et al. 2023) provide context to the Sprattus system, allowing an analysis of to which extent adaptation to a brackish environment shows genetic parallelism in two closely related species. Although the species are phylogenetically close, their split far exceeds the formation of the shared brackish waters, the Baltic Sea (approx. 12 kY old (Lass and Matthäus 2008)) and Landvikvannet (150 Y old), ensuring that adaptation took place independently in each species.

Currently, genetic differentiation among sprat populations has been assessed using reduced-representation sequencing approaches, resulting in a set of relatively internally homogenous genetic groups: (*a*) Coastal Norwegian; (*b*) Northeast Atlantic including the North Sea, Celtic Sea, and Bay of Biscay; (*c*) Baltic Sea (McKeown et al. 2020; Quintela et al. 2020). Likewise, distinct groups were found in the Adriatic and Black Seas (Quintela et al. 2020) as well as in Landvikvannet, Norway (Quintela et al. 2021), respectively. However, the sparseness of markers assessed in these studies, coupled with the lack of a reference genome, has precluded further characterization of the putative adaptive signals.

Here, we provide a draft assembly of the sprat genome, whole-genome pool-seq data from 19 populations spanning from the Baltic to the Black Sea and individual genotypes (*n* = 369 specimens), using the recently released MultiFishChip (Andersson et al. 2024). We identify strong signals of genetic adaptation to brackish waters, including six putative inversions. Additionally, we identify three sharp peaks of associations involving single genes that each overlap with a similarly narrow signal in Atlantic herring, providing evidence for some genetic parallelism between the two species as regards genetic adaptation to low salinity.

## Results

### Genome assembly and pool re-sequencing

The primary assembly, based on 14 M PacBio CSS reads, is composed of 2,115 contigs (contig *N50* = 1.0 Mb) with a total length of 971 Mb. This contig assembly was scaffolded based on 287 M HiC-read pairs, using pin-HiC (v3.0.0). It was thereafter manually curated using Juicebox (v1.11.08) and a custom de-duplication procedure, based on mapped read-depth and location of duplicated BUSCOs. This resulted in a total length of 785 Mb, with scaffold *N50* = 23.4 Mb and scaffold *L50* = 12 Mb. The total length is almost identical to the 786 Mb genome assembly of the Atlantic herring, including unplaced scaffolds (Pettersson et al. 2019).

The signal-to-noise ratio of the HiC-mappings was unusually poor by modern standards, possibly due to the comparatively high diversity and apparent high rate of structural variation between the chromosome copies, and the underlying PacBio assembly proved resistant to commonly used de-duplication methods. This leads to some ambiguity regarding scaffold breaks in the final assembly. In particular, the largest scaffolds are substantially longer than expected, given the size distribution of the herring chromosomes. Thus, in a subset of the following analyses, we relied on liftover to the current herring reference assembly to organize the results. Overall, this liftover revealed a good correspondence between the two assemblies, with 19 of 26 herring chromosomes mapping near-exclusively to a single sprat scaffold (Supplementary Figures 1 & 2). However, it also revealed that the two very large sprat scaffolds (s1111 and s1118) correspond to two (s1111: Chrs 21 and 23), and three (s1118: Chrs 8, 24, and 26) herring chromosomes, respectively, with the physical order along the sprat scaffolds strongly correlating with the order on herring chromosomes, i.e., herring Chr 21 maps to the beginning of s1111, while Chr 23 maps to the end. Whether these large scaffolds reflect true chromosome fusion events or assembly errors is currently unclear, but it is noteworthy that for both s1111 and s1118 there are cross-mappings into sections of the sprat scaffold that is otherwise homologous to another herring chromosome. This pattern of out-of-order matches to the same scaffold would be expected if there were post-fusion intra-chromosomal rearrangements, a common occurrence in teleosts, but otherwise would be unlikely given the sparsity of cross-mapping to other scaffolds, also the ends of these non-cognant blocks do not co-occur with contig breaks (Supplementary Figure 3). Unfortunately, there is no published karyotype data available for the sprat to the best of our knowledge that could be used to verify the presence of two particularly large chromosomes suggested by this assembly.

This scaffolded assembly was used to call SNPs in the 19 sample pools. The number of individuals per pool was in the range 16-24 (Supplementary Table 1). The pools were sequenced to an average depth of 36x coverage of paired Illumina NovaSeq reads, resulting in 2.8 M SNPs retained after stringent filtration. Samples were collected from a broad geographic area (Figure 1), from Nordfjord on the Norwegian coast in the North to the Celtic Sea in the West and the Black Sea in the South-East.

**Figure 1.**
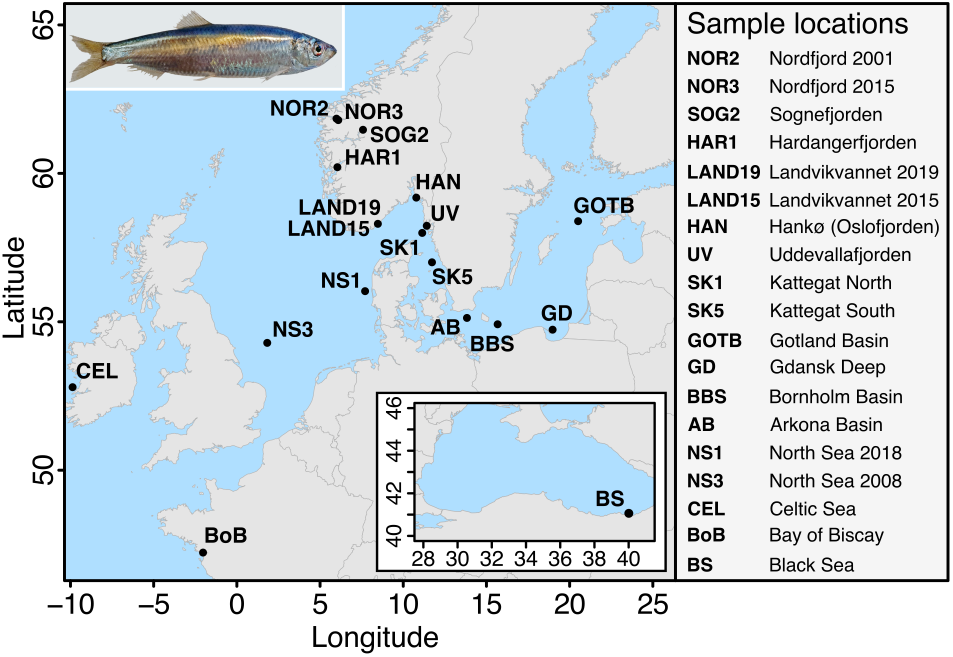
Sample locations. Geographic location of the 19 population samples of European sprat (Sprattus sprattus) used for pool-seq analysis. Sample codes are provided in parentheses. Photo: Merete Kvalsund, Institute of Marine Research, used with permission.

### Nucleotide diversity

Since the PacBio assembly provides complementary primary and alternative contigs, corresponding to the two chromosome copies in the reference individual, for most of the genome, we can use these to estimate a genomic average for nucleotide diversity (*π*). Based on the median of distribution of observed diversities in 362 alignment blocks (Supplementary Figure 4), we estimated average π to be 1.2% – four times the value in Atlantic herring (Martinez-Barrio et al. 2016) – but also note that the variance among blocks is substantial (*s.d.* = 1.1%), indicating that the average number is not representative of all genomic regions.

### Overall population structure

Based on the approximately 2.8 million SNPs that passed quality filtering, we obtained the neighbor-joining tree shown in Figure 2a, which shows that, based on the entire genome, samples group by habitat rather than geographic location. The three groups indicated in the tree correspond to distinct ecotypes: Oceanic, Coastal and Brackish. These will form the basis for the genetic contrast reported below. The “Brackish” group comprises three separate localities, the Baltic Sea (samples Gotland Basin, Gdańsk Deep, Bornholm Basin, and Arkona Basin), the Black Sea and Landvikvannet in Norway, and, in spite of their geographic dispersion, they form a well-defined clade. The “Oceanic” group consists of oceanic samples from Kattegat, North Sea, Celtic Sea, and Bay of Biscay whereas the “Coastal” group represents Norwegian coastal waters and fjords. The samples from Oslofjorden and Uddevallafjorden were placed between the Oceanic and Coastal groups suggesting they may represent admixed populations and, to reduce complexity, these two samples were therefore not included in the contrasts described below.

**Figure 2.**
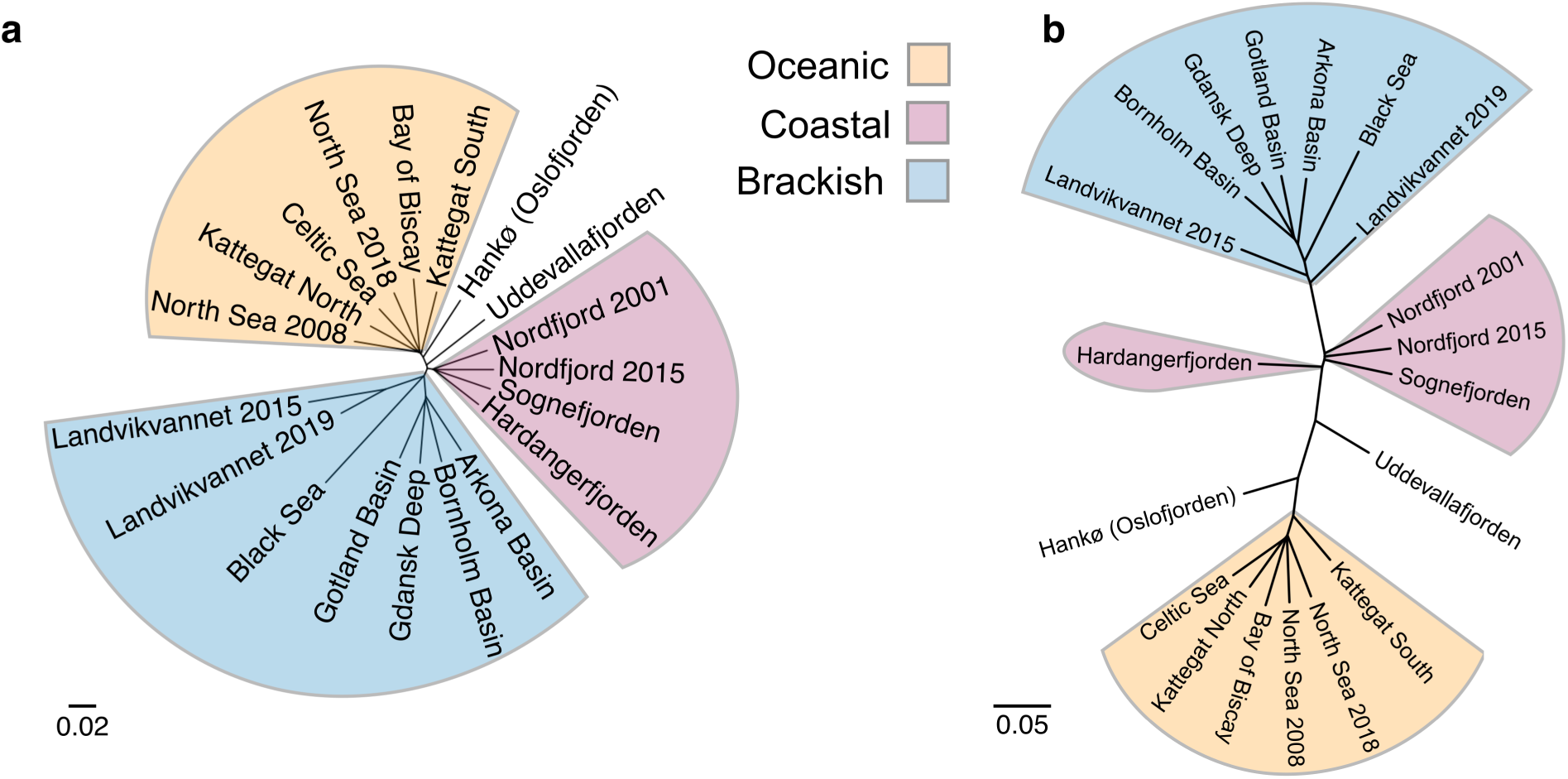
Neighbor-joining tree based on allele frequency distances among pools. *a*) Based on the entire genome. *b*) Using only highly differentiated markers (delta allele frequency > 0.5). The major groupings that have been used throughout the study are indicated by the colored slices and corresponding labels.

Contrasting the average allele frequencies in the three groups defined above reveals that the great majority of SNPs occur at very similar allele frequencies across populations, within the expected fluctuation due to sample size. However, the distribution contains a long tail of SNPs (Supplementary Figure 5). This mimics the situation in the Atlantic herring, in which the *F_ST_* distribution among populations deviates significantly from the one expected for selectively neutral alleles under a genetic drift model (Lamichhaney et al. 2017).

To validate the patterns of genetic differentiation detected using pooled whole genome sequencing (WGS), we employed the recently released MultiFishChip (Andersson et al. 2024), a resource developed to support cost-efficient typing of informative markers in several teleost species, to generate genotype information for 369 sprat individuals from 19 sampling sites largely overlapping the WGS sample set (Supplementary Table 1). The sprat component of the MultiFishChip (Andersson et al. 2024) was designed based on the WGS data presented herein (see methods) and the SNP-designs are included as Supplementary Data 1.

A PCA biplot built using 2,063 LD-pruned SNPs differentiated the Oceanic, Coastal, Landvikvannet, Baltic, Adriatic-Black Sea populations (Figure 3a). Most of the individuals from the Uddevalla fjord (UV) clustered with samples from the Coastal population while a few joined the Oceanic samples thus demonstrating that this is an admixed sample as suggested from pooled sequencing (see above). In contrast with the homogeneity of the Oceanic sprat, some individuals from Landvikvannet (LAND19) clustered with Coastal samples whereas other from the same location clustered with the brackish group, indicating that this is also a mixed sample. Discriminant Analysis of Principal Components (DAPC) revealed that the distribution of the individuals along the first axis (59.7% of the variation) seemed to follow a longitudinal gradient with the sample from the Black Sea occupying the farthest extreme (Supplementary Fig. 6a), whereas the second axis (16.5%) discriminated the Baltic sprat and placed UV in an intermediate position between Oceanic and Baltic fish. The third axis (11.9%) further separated Landvikvannet sprat samples leaving the recent one (LAND19) closer to the Coastal sprat (Supplementary Fig. 6b). However, the dendrogram coupled with the pairwise *F_ST_* matrix revealed a first dichotomic division between brackish (plus the Adriatic Sea) and non-brackish environments (Figure 3b). Non-brackish sprat was divided into Coastal and Oceanic with UV stemming from the Coastal branch. The genetic differentiation was essentially null among population samples within Coastal, Oceanic and Baltic sprat, respectively (*F_ST_* in the range 0-0.006). In Landvikvannet, the average differentiation towards the Coastal sprat was 0.078 in 2015 and circa half this amount (0.037) four years later. If we, instead of pruning by LD, use all high-quality markers selected from the high differentiation regions found in the WGS analysis (*n* = 2,354), the pattern somewhat changes; the dichotomy between Oceanic sprat and the remaining ones is emphasized, while the Black Sea comes closer to the Baltic Sea samples (Figure 3c,d), mirroring the WGS results. The reason for this is that SNPs from the six inversions, shared between Baltic Sea and Black Sea populations, have a prominent impact on the pattern.

**Figure 3.**
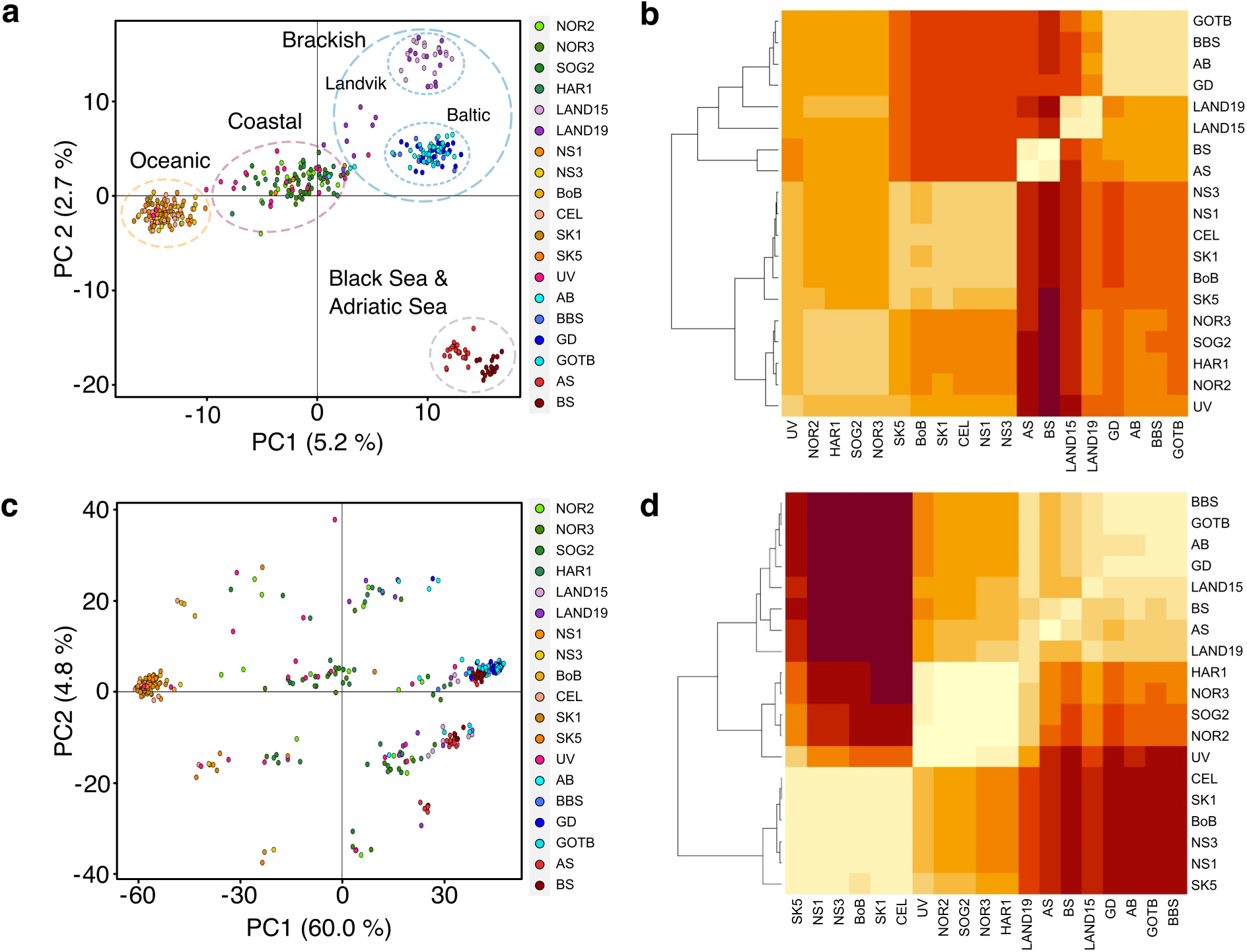
SNP-chip based PCA and FST heatmaps. *a*) PCA analysis based on 2,063 LD-pruned markers. *b*) Heatmap showing pair-wise *F_ST_* values, based on the markers used in (*a*). *c*) PCA analysis based on 2,354 highly-differentiated markers. *d*) Heatmap showing pair-wise *F_ST_*values, based on the markers used in (*c*), light yellow indicates low *F_ST_*, deep red indicates high *F_ST_*. Sample codes are given in Fig 1 and Supplementary Table 1.

The outcome of STRUCTURE (Pritchard et al. 2000) analyzed through Evanno’s test revealed *K*=2 as the most likely number of genetic groups (Supplementary Fig. 7a), which discriminated Landvikvannet, Baltic, Adriatic and Black Sea from the Oceanic samples and leaving admixed Coastal sprat and hybrid zone sprat. In contrast, Puechmaille’s statistics reported *K*=5 (Supplementary Fig. 7b), and revealed four rather homogenous clusters (Landvikvannet, Oceanic, Baltic and Adriatic/Black Sea, respectively).

### Genomic regions of differentiation among subpopulations

Using the scaffolded version of the sprat genome assembly, we performed a genome wide compilation of independent regions of differentiation, defined as having a gap of at least 500 kb to the next highly significant SNPs (Figure 4). The threshold was chosen to minimize the risk of artificially dividing a region that, represents a single signal at the expense of possibly combining nearby, but independent, signals. We estimated differentiation for two contrasts: Oceanic vs Brackish (*OvsB*) and Oceanic vs Coastal (*OvsC*). This yielded a total of 103 regions in the *OvsB* contrast. If we exclude regions consisting of a single SNP, this number drops to 68. The regions cover 14.0 Mb of the genome, with six putative inversions comprising the majority of that total. All identified regions are listed in Supplementary Table 2. These six genomic regions were identified as putative inversions because they showed the characteristic of equally strong genetic differentiation over a large genomic region, on the megabase scale, and sharp borders to the flanking region not showing differentiation (Figure 5).

**Figure 4.**
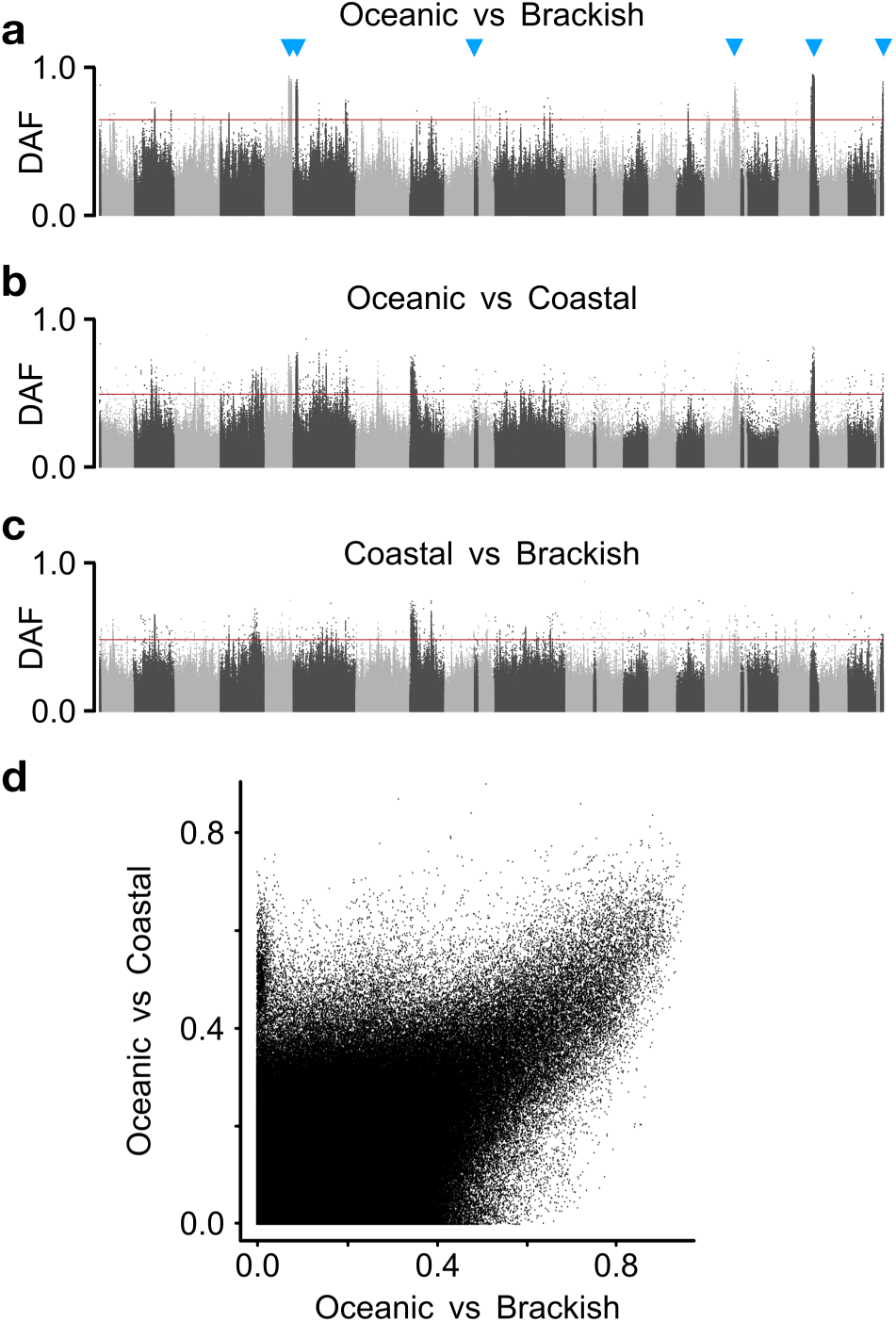
Delta Allele Frequencies (*DAF*) of SNPs in three genome-wide contrasts. *a*) Oceanic vs Brackish populations. Blue arrows mark the location of six putative inversions. *b*) Oceanic vs Coastal. *c*) Coastal vs Brackish. *d*) Scatter plot of the SNP data from (*a*) and (*b*).

**Figure 5.**
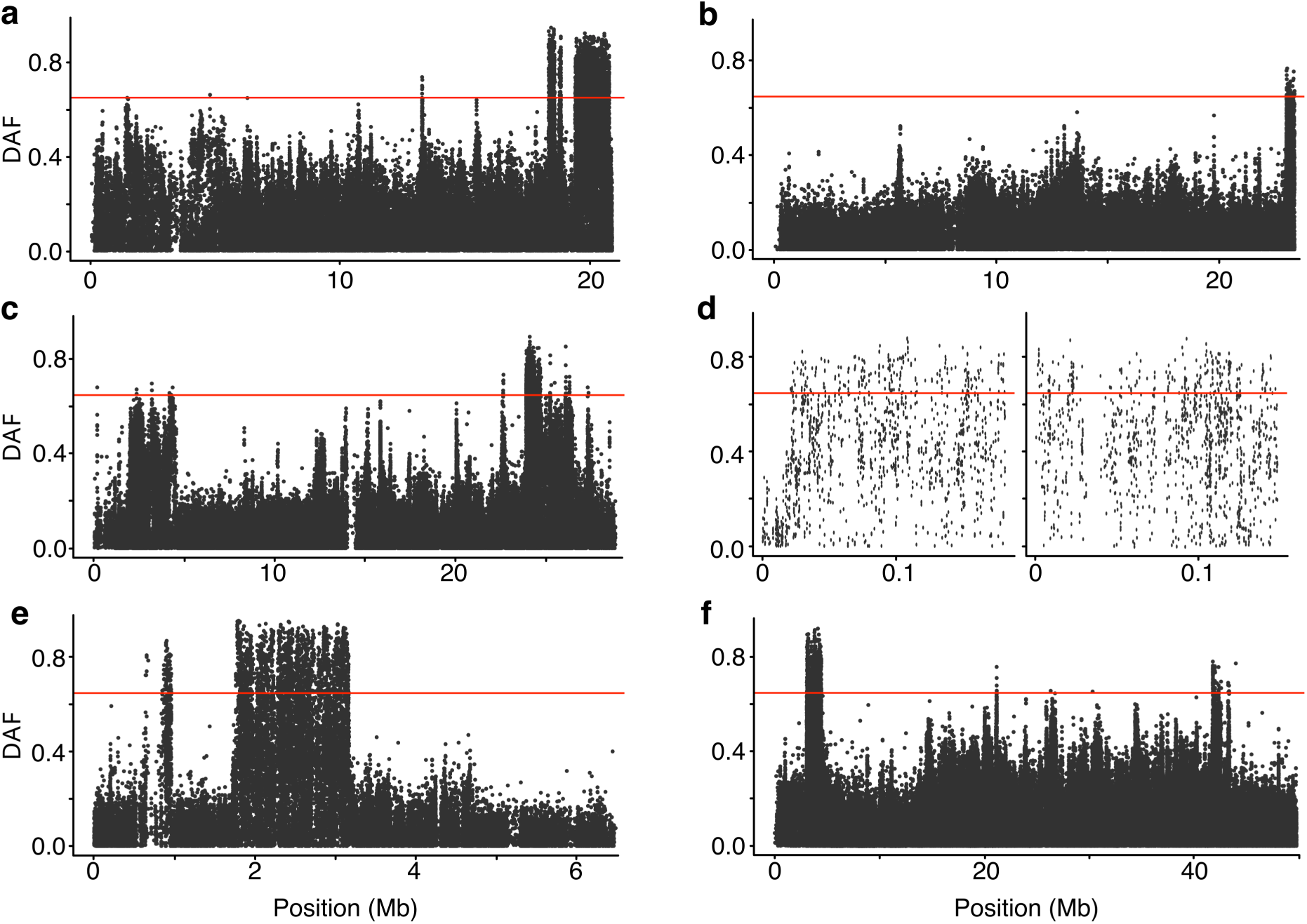
Six putative inversions detected in the “Oceanic” vs “Brackish” contrast. Per-SNP Delta Allele Frequencies (*DAF*) high-light putative inversions, the regions are found on the following scaffolds: s1110 (*a*), s1114 (*b*), s13(*c*), s173* and s374*(*d*), s40 (*e*) and s1111(*f*). *This signal is located on two minor scaffolds in the sprat assembly, that both map to nearby regions on *C. harengus* chromosome 13.

The *OvsC* contrast contains more regions than the *OvsB* contrast: 286 in total, 138 if excluding single SNP regions, and also covering a substantially larger part of the genome (48.9 Mb). However, this should not be taken as evidence for a stronger genetic differentiation in this contrast. The lower Delta Allele Frequencies (*DAF*s) of the inversion regions, in particular, lowers the standard deviation of the overall distribution in the *OvsC* contrast and thus the significance threshold (0.65 in *OvsB* vs 0.50 in *OvsC*). In essence, the peaks of divergence in *OvsC*, while more numerous, are noticeably less differentiated than in *OvsB* (Figure 4a,b). Furthermore, there is a strong overlap with the *OvsB* contrast, with a total of 11.2 Mb being called as significant in both contrasts. In general, the Coastal populations are more similar to the Brackish populations than to the Oceanic ones, in particular at strongly differentiated loci, causing less differentiation overall in the Brackish vs Coastal (*BvsC*) contrast (Figure 4c), as well as a relatively strong correlation (*r^2^* = 0.30) between *DAF* values in the *OvsC* and *OvsB* contrasts (Figure 4d). Nevertheless, there are signals specific to the Coastal populations, as can be seen by a cluster of SNPs with *DAF* in the range 0.4-0.8 in the *OvsC* contrast but *DAF* < 0.1 in the *OvsB* contrast (Figure 4d). The majority of those SNPs are located within a region on scaffold s1113, corresponding to 0.1 to 1.0 Mb on herring Chr 12. This region harbors another putative inversion, for which all Coastal populations have intermediate to high frequencies of a haplotype that is essentially missing from all other samples (Supplementary Fig. 8).

We used the data from the SNP-chip analysis for a Redundancy Analysis (RDA) as an alternative method to identify loci potentially under selection, resulting in 84 detected outlier SNPs. Most of the outliers flagged by the RDA were associated to salinity (*n* = 60), followed by dissolved oxygen (*n* = 11), and temperature of sea surface (*n* = 9) whereas only two outliers were associated to pH and current velocity, respectively. The SNPs used in the RDA are listed in Supplementary Table 3, while the outlier SNPs are found in Supplementary Table 4.

### Six putative inversions associated with ecological adaptation to brackish waters from the Black Sea to the Baltic Sea

We identified six putative megabase-scale inversions strongly associated with adaptation to brackish waters (Figure 5). Inspection of allele frequencies for SNPs in the putative inversions revealed a common pattern; i.e., that the Coastal population samples were intermediate between the Oceanic and Brackish ones, and that in many cases the Black Sea and the Baltic samples appeared to carry closely related haplotypes at high frequencies (Figure 6, Supplementary Fig. 9). This provides an explanation for the topology of the phylogenetic tree (Figure 2a) because many of the most differentiated SNPs are located within the putative inversion regions. A tree based on markers from outside the strongly differentiated regions shows a similar topology, albeit with a relatively longer branch to the Black Sea sample (not shown). This indicates some allele sharing among brackish samples also outside the most divergent regions. On the other hand, using only differentiated markers, defined as having *DAF* higher than 0.5 in any contrast, reinforces the “Brackish” and “Oceanic” groupings, while positioning the “Coastal” samples along the main branch (Figure 2b). In addition to the signals of divergence consistent with the main groupings, we also detected other genome regions with strong differentiation between samples, but with patterns that do not exactly conform to the three main groups. Instead, they are restricted to a subset of the brackish samples, and include a region around 5.8 Mb on HiC-scaffold 1144, where the signal is restricted to the Baltic Sea (Supplementary Fig. 10), and a 1 Mb region from 18.5 to 19.5 Mb on HiC-scaffold s4, where the Landvikvannet and Black Sea samples display similar allele frequencies that are distinct from that in other populations (Supplementary Fig. 11).

**Figure 6.**
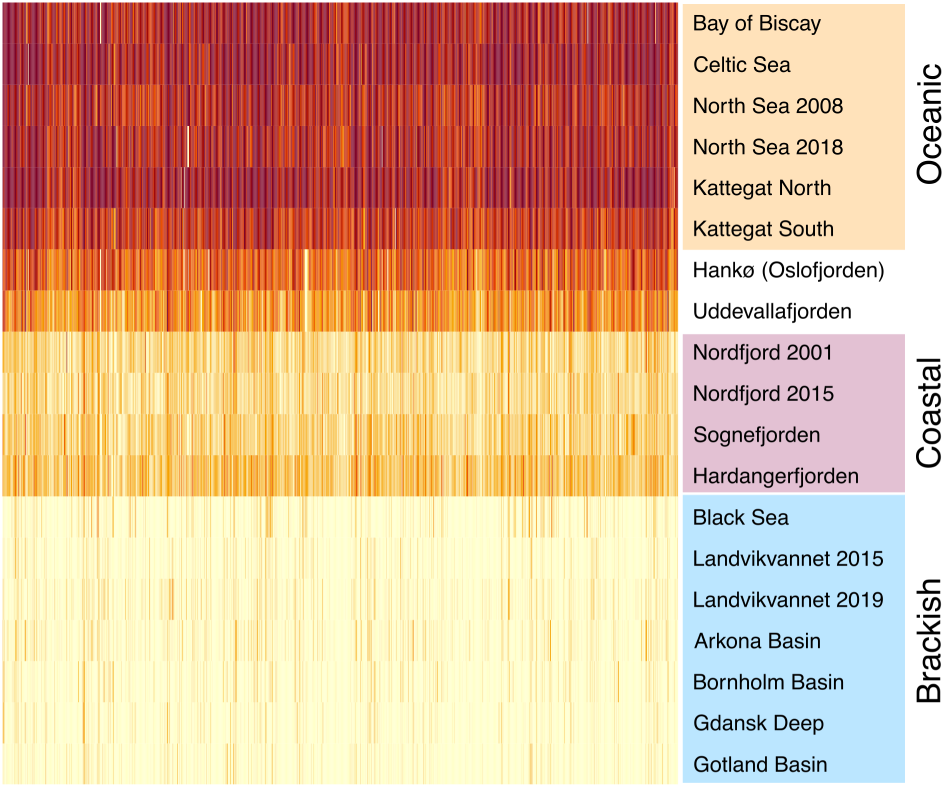
Heatmap showing allele frequencies at the putative inversion on sprat scaffold s40. Population groupings are indicated by labels and colored boxes, in accordance with Figure 2. This 1.5 Mb region in sprat (shown in Fig. 5e) corresponds to two regions on *C. harengus* Chr 18 (13.9-16.5 Mb and 17.9-19.3 Mb) suggesting a rearrangement between the two species. Heat maps for the other five putative inversions are shown in Supplementary Fig. 9.

### Limited genetic parallelism between European sprat and Atlantic herring

The sprat and herring are closely related clupeids that show genetic adaptation to brackish waters. The liftover procedure allowed us to explore to which extent this adaptation shows genetic parallelism, i.e., that genetic variation in the same genes have contributed to adaptation. The main outcome of these analyses is that the observed parallelism is limited to a few loci. Firstly, none of the sprat’s six putative inversions can be identified in the Atlantic herring. Secondly, there are about 125 independent loci that show strong genetic differentiation between Atlantic and Baltic herring (Pettersson et al. 2019) and the great majority of these loci do not show an obvious overlap with signals of selection detected in the marine-brackish contrast in sprat. However, for three genes the narrow signals of selection overlap. Firstly, one of these regions is located around 2.3 Mb on chromosome 12 in Atlantic herring and harbors the prolactin receptor (*PRLRA*) gene only (Figure 7). A similar narrow signal involves herring chromosome 19 containing the thyroid receptor beta (*THRB*) gene (Supplementary Figure 12a). Finally, there is a signal covering a cluster of troponin I2 (*TNNI2*) genes on herring chromosome 3 (Supplementary Fig. 12b). We did not find any missense mutations in any of these three genes, neither in sprat nor in herring.

**Figure 7.**
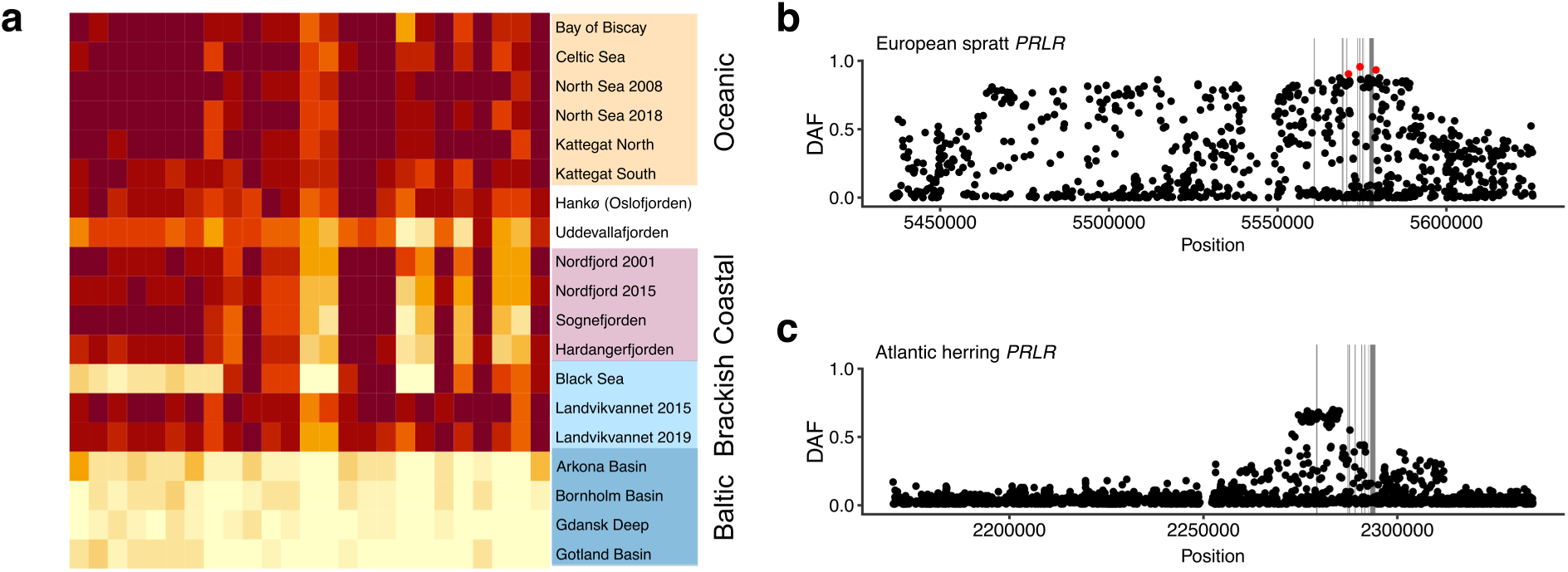
Genetic parallelism between European sprat and Atlantic herring – *PRLRA*. *a*) Heatmap of the *PRLRA* region in European sprat. Samples are colored as in Figure 2, with the exception of Baltic samples being highlighted within the Brackish group. *b*) Zoom-in showing *DAF* in the Oceanic and Brackish contrast, in European sprat, across the *PRLRA* locus (found on scaffold s1113 in the European sprat assembly). *c*) Zoom-in showing *DAF* between Atlantic and Baltic herring populations across the *PRLRA* locus on Chr 12 (5). In (*b*) and (*c*), grey boxes indicate location of exons.

## Discussion

Here, we have shown that the European sprat is a species with high, albeit not extreme, genetic variability, with overall nucleotide diversity estimated to be in excess of 1%. This is considerably higher than the 0.3% found in the Atlantic herring (Martinez-Barrio et al. 2016), which we tentatively attribute to the fact that the sprat has a more southerly habitat range, reducing the risk of severe bottlenecks during periods of extensive polar ice cap expansion. The colonization of brackish waters has led to marked genetic differentiation, approaching fixation of different alleles, at a discrete set of loci across the genome. This adaptive response is, on the other hand, qualitatively similar to that previously shown in Atlantic herring (Martinez-Barrio et al. 2016; Pettersson et al. 2019; Han et al. 2020). However, while this study has shed light on the differentiation between major groupings of sprat populations, the differentiation within these groups remains very limited with FST values in the range 0-0.006 between population samples. However, this result is based on a very limited number of sampled populations, not quite reaching a quarter of the number used by Han et al. (Han et al. 2020) for Atlantic herring. A cost-efficient way of remedying this situation is to use a SNP-chip, and, to this end, the signals identified here have been used to select markers for the sprat component of the recently released MultiFishChip (Andersson et al. 2024) containing thousands of SNPs from each of seven species including the European sprat and Atlantic herring.

Like the herring, sprat has adapted to spawning in brackish conditions. This is important, since heterogeneity in environmental conditions during spawning has recently been brought up as a critical aspect resulting in genetic differentiation between subpopulations due to natural selection (Fuentes-Pardo et al. 2023; Andersson et al. 2024). This puts species that spawn in diverse environmental conditions, but may intermingle as adults, in contrast with species that spawn communally and then later on disperse. The archetypical example of the latter is the European eel (*Anguilla anguilla*), in which all spawning takes place in the Sargasso Sea. Consequently, that species has no discernible signs of genetic adaptation in response to the environments encountered by the adults, in spite of the habitat range for the adult stage extending, in geographic terms, from the Baltic to the Mediterranean and, in terms of salinity, from fully marine to freshwater (Enbody et al. 2021). The sprat and according to a recent study the Atlantic horse mackerel (*Trachurus trachurus*) (Fuentes-Pardo et al. 2023) appear intermediate between two extremes, the European eel being one panmictic population and Atlantic herring showing extensive genetic differentiation (4). An important difference between sprat and Atlantic horse mackerel on one hand and herring on the other, is that the former are pelagic spawners (de Silva 1973; Abaunza et al. 2003), whereas herring deposits fertilized eggs at specific bottom types or on vegetation (Runnström 1941; Aneer et al. 1983). The latter are therefore exposed to a much more heterogenous environment than eggs from a pelagic spawner, possibly resulting in a difference in local selection pressure. Furthermore, it is expected that a homing behavior is less precise for a pelagic spawner than for a fish where the fertilized eggs are developing at spawning grounds.

We have strong indications that at least six putative inversions play important roles in the adaptation to brackish conditions in sprat. In Atlantic herring there are at least four major inversions with prominent frequency gradients along a North-South axis, but no inversion appears to be underlying adaptation to low salinity (Han et al. 2020). The observation that the same inversion haplotype groups are present at high frequencies in geographically distant brackish populations separated by marine habitats (Fig. 1) suggests that the brackish haplotypes may occur at low frequencies in marine populations, and in particular in coastal populations which display temporary brackish conditions. These haplotypes can then be subject to strong positive selection when sprat colonize brackish environment. This must have happened when Landvikvannet in Norway was colonized about 150 years ago after this former freshwater lake became a brackish environment when a canal to the ocean was established for transporting timber. This mimics the situation in sticklebacks where marine populations constitute a reservoir of alleles critical for adaptation to freshwater conditions (Jones et al. 2012).

The putative inversions here reported for sprat and the confirmed inversions in herring (Han et al. 2020; Jamsandekar et al. 2023) all associated with local adaptation, adds to a growing list of so-called supergenes contributing to ecological adaptation in marine organisms. For instance, in Atlantic cod, four large chromosome inversions (on chromosomes 1, 2, 7 and 12) are allegedly linked to a migratory lifestyle and environmental adaptations such as salinity tolerance (Berg et al. 2016; Berg et al. 2017; Barth et al. 2019; Matschiner et al. 2022). Similarly, three large putative chromosomal inversions have been found to be associated with sea surface temperature in the king scallop (*Pecten maximus*) (Hollenbeck et al. 2022).

An important question in evolutionary biology is how often the same gene contributes to a similar genetic adaptation in different species (Conte et al. 2012). One example is that selection at the human *EPAS1* gene has contributed to adaptation to high altitude in both Tibetan and Andean highlanders (Lawrence et al. 2024). Another striking example of convergence at the molecular level concerns genes encoding visual opsins that often respond to selection related to differences in light conditions among species habitats (Musilova et al. 2021). For instance, one third of all fish species adapted to freshwater or brackish waters, including Baltic herring, express a rhodopsin protein with tyrosine at residue 261, whereas essentially all marine fish have phenylalanine at this position (Hill et al. 2019). However, most biological traits have a highly polygenic background (Pritchard and Di Rienzo 2010), which implies that such striking genetic parallelism is expected to be uncommon. The genome-wide screens of the two closely related clupeids European sprat and Atlantic herring as regards their adaptation to the brackish Baltic Sea provide an excellent opportunity to explore how common genetic parallelism is. Our conclusion is that it is limited because the adaptation in sprat is dominated by six putative inversions not being present in Atlantic herring, and out of the 100+ loci identified in Atlantic herring (Han et al. 2020) only three show a striking overlap with signals in sprat. One of these three genes, *TNNI2* does not have an obvious link to adaptation to a brackish environment; *TNNI2* encodes troponin I2, an important component of fast twitch muscles. *THRB* encodes thyroid hormone receptor beta, a nuclear receptor that affects gene expression in many cell types. Of potential relevance to the selection signals in sprat and herring is that *thrb* in zebrafish has a critical role for development of retinal red cones and long-wave vision (Volkov et al. 2020). This is of interest because of the strong selection acting on visual opsins in fish in general (Musilova et al. 2021) and for improved vision in the red spectra in the Baltic Sea (Hill et al. 2019). The third case of genetic parallelism concerns the prolactin receptor (*PRLR*) gene (Figure 7), which has an obvious link to adaptation to a brackish environment due to its important role in osmoregulation (Manzon 2002). The hormone prolactin has many functions in vertebrates, including stimulation of milk secretions in mammals as indicated by its name. However, in fish it constitutes the freshwater-adapting hormone in euryhaline species (Manzon 2002). Prolactin is released from the pituitary but its effect on target tissues (gills, kidney, intestine, urinary bladder and skin) is mediated by prolactin receptor signaling. The fact that we did not find any *PRLR* missense mutations neither in sprat nor in herring implies that genetic adaptation to the brackish environment is mediated by an altered expression pattern. The results imply that *PRLR* may be a gene that often show genetic parallelism in fish species adapted to differences in salinity.

## Materials and methods

### Sample collection

All the individuals typed in the different steps of this study, with the exception of samples HAN and AS (the latter of which was not used for re-sequencing due to low DNA quality), were formerly used to outline sprat management boundaries (ICES 2018; Quintela et al. 2020) and, in the case of the samples from Landvikvannet (LAND15 & LAND19), to describe the genetic response to human-induced habitat changes (Quintela et al. 2021). Sample HAN was collected in the Oslofjorden area in 2018 (59.2° N, 10.9° E), whereas sample AS was collected in 2021 in the Adriatic Sea (43.1° N, 13.9° E) during the MEDIAS survey (Leonori et al. 2021), following a common protocol. DNA was extracted from fin clips stored in ethanol using the Qiagen DNeasy 96 Blood & Tissue Kit in 96-well plates; each of which contained two or more negative controls.

### Preparation of high-molecular weight DNA

Flash-frozen tissue from one individual sprat was collected from Langenuen in the outer part of Hardangerfjorden, Norway (59.975°N, 5.376°E). High-molecular weight DNA for PacBio long read sequencing was extracted using the Nanobind Tissue Big DNA Kit v1.0; Standard TissueRuptor Protocol. The genomic DNA was subsequently cleaned using the Pacific Bioscience “Guidelines for Using a Salt:Chloroform Wash to Clean Up gDNA”. DNA concentration was measured using the Qubit Broad Range kit (Thermo Fisher Scientific #Q32850), purity was checked by UV-absorbance, and fragment lengths were determined using the Genomic DNA 165 kb Kit (Agilent #FP-1002-0275) on the Femto Pulse System.

### Long read sequencing and draft assembly construction

1.47*106 M PacBio CCS-reads (≥Q20) covering a total of 24.6*10^9^ bases, was generated from a single flow cell (SMRT link v8.0.0.79519). This read-set was used to create the contig version of the draft assembly, using IPA v1.3.2 (https://github.com/PacificBiosciences/pbipa). The scaffolded version was done bases on the above-mentioned contain assembly and 287 M HiC-read pairs, using pin-HiC (v3.0.0) (https://github.com/dfguan/pin_hic). The pin-HiC output was manually curated using Juicebox (v1.11.08) (Dudchenko et al. 2018) and a custom de-duplication procedure based on read depth and the location of duplicated BUSCO hits.

### Short read sequencing

Illumina short read sequencing, using the standard configuration of paired 150 bp reads, of pooled population samples was performed at NGI Uppsala - SNP&SEQ Technology Platform. Data was generated using one lane of Illumina NovaSeq S4, comprising 3.2*10^9^ read-pairs (1.69 ± 0.38 *10^8^ per pool), resulting in 9.7*10^11^ bases sequenced. The reads are available at NCBI’s Short Read Archive (BioProject: PRJNA1023385).

### Read mapping and SNP calling

Reads were mapped to the draft sprat assembly (see above) using bwa mem (Li 2013) (v 0.7.17-r1188). The resulting bam files were then processed using GATK (McKenna et al. 2010) (v4.1.1.0), with the following workflow: First, we used “HaplotypeCaller” to generate per-sample gvcf-files. These were merged using “CombineGVCFs” and genotyped using “GenotypeGVCFs.” The raw SNP genotypes were then passed through the “VariantFiltration” module, with the following arguments:

--filter-name “stringent_combined_filter” --filter-expression “QD < 8.0 || FS > 50.0 || MQ < 30.0 || MQRankSum < -10.0 || ReadPosRankSum < -6.0” --filter-name “depth_filter” --filter-expression “DP < 200.0 || DP > 2500” --missing-values-evaluate-as-failing.

### Bioinformatic analysis

The bioinformatic analysis was performed using custom R (R Core Team 2019) scripts; these are deposited in the “Sprat_pool_reseq” repository at the LeifAnderssonLab GitHub page (https://github.com/LeifAnderssonLab/Sprat_pool_reseq).

### Genome-wide nucleotide diversity

We estimated genome-wide nucleotide diversity (*π*) by comparing the sequences in the primary and alternative assembly from the reference individual. First, we performed a “satsuma chromosemble” run to align the primary and alternative assemblies. Then, we randomly selected a set of positions (*n*

= 600) from the primary assembly to serve as starts for alignment blocks. Starting at these positions, we extracted 25 kb regions and queried the Satsuma output for a matching contig in the alternative assembly. These were then aligned using Clustal Omega (Sievers and Higgins 2018), and then diversity in the block was calculated using the “dist.dna” method from the R package “ape” (Paradis and Schliep 2019). Blocks yielding either no, or very short, alignments – typically indicative of the corresponding contig being missing for the alternative assembly – were eliminated, as were those with observed divergence > 5%. The latter outcome is not consistent with regular sequence divergence, but rather likely to be the result of alignment artifacts induced by structural differences between the two haplotypes. Similar issues are likely to contribute to the highest of the retained values as well, which is why we are using the median, rather than mean, observed divergence as our estimate of *π*. Retaining all alignments would lead to a small increase in estimated *π*, from 1.2% to 1.3%.

### MultiFishChip design

The sprat component of the MultiFishChip was based on the data presented herein, and comprises three subsets. First, neutral set containing SNPs with low variation between groups but comparatively high minor allele frequencies (*MAF* >= 0.3), estimated across the entire set of samples. Second, a set selected from the contrast based analysis presented here, with representative SNPs selected form identified signals of divergence. Lastly, a set of SNPs with high variance across populations, not tied to any particular pre-defined contrast. Together, the design resulted in 7742 candidate markers (Supplementary data 1), out of which 5916 were eventually included on the MultiFishChip.

### MultiFishChip analysis

A subset of 381 individuals from 19 samples was sent to IdentiGEN (Ireland) for genotyping with the MultiFishChip SNP array (Andersson et al. 2024); 18 of the samples were overlapping with the WGS set, the sample from Hankø (HAN) had to be discarded due to amplification issues whereas the sample from the Adriatic Sea was not available for WGS but could be acquired and genotyped with the SNPchip at a later stage of the project. Twelve of the individuals failed to amplify, thus leaving 369 fish ranging between 16 (BoB) and 24 (AS) per sample. Data were obtained for 4,602 out of the 5,916 SNPs of the chip, 75 of which were discarded due to >10% missing data. PLINK (Purcell et al. 2007) was used to LD-prune the dataset (*r^2^* = 0.25, *MAF* > 0.05) thus leaving 2,063 SNPs for statistical analyses. Principal Component Analysis (PCA) was conducted using the function “dudi.pca” in ade4 (Dray and Dufour 2007). The relationship among geographically-explicit samples was assessed using pairwise *F_ST_* (Weir and Cockerham 1984) and through the Discriminant Analysis of Principal Components (DAPC) (Jombart et al. 2010) implemented in adegenet (Jombart 2008) using the cross-validation function (Jombart and Collins 2015; Miller et al. 2020) to avoid overfitting. In addition, STRUCTURE (Pritchard et al. 2000) (v.2.3.4) analysis was conducted using the software ParallelStructure (Besnier and Glover 2013) to identify genetic groups under a model assuming admixture and correlated allele frequencies without using population information. Ten runs with 100,000 burn-in and 1,000,000 MCMC iterations were performed for *K* = 1 to *K* = 5 clusters. STRUCTURE output was analyzed using *a*) the *ad hoc* summary statistic *ΔK* of Evanno et al. (Evanno et al. 2005), and *b*) the four statistics of Puechmaille (Puechmaille 2016), both implemented in StructureSelector (Li and Liu 2018). Finally, runs for the selected *K*s were averaged with CLUMPP v.1.1.1 (Jakobsson and Rosenberg 2007) using the FullSearch algorithm and the G’ pairwise matrix similarity statistic, and graphically displayed using bar plots.

Redundancy Analysis (RDA) is a genotype-environment association (GEA) method to detect loci under selection (Forester et al. 2018). Environmental data (temperature, salinity, pH, dissolved oxygen, current velocity, chlorophyll) was obtained for the different sampling points using the Bio-ORACLE database https://www.bio-oracle.org (Tyberghein et al. 2012; Assis et al. 2018). Collinearity between variable pairs was investigated and only non-correlated ones were retained for analyses. Analysis was conducted with the R package vegan v.2.5–7 (Oksanen et al. 2019) using the 2,063 LD-pruned loci with samples classified into Brackish, Oceanic and Coastal. The samples from the hybrid zone and the Adriatic Sea were discarded for not falling in any of these categories.

## Supplementary material

Supplementary material is available online.

## Acknowledgements

The project was financially supported by the Research Council of Norway (CoastRisk project -299554), Vetenskapsrådet (2017-02907; to LA), Knut and Alice Wallenberg Foundation (KAW 2016.0361; to LA). The National Genomics Infrastructure (NGI)/Uppsala Genome Center provided service in massive parallel sequencing and the computational infrastructure was provided by the Swedish National Infrastructure for Computing (SNIC) at UPPMAX partially funded by the Swedish Research Council (2018-05973).

## Author contributions

L.A. and M.Q. conceived the study. M.E.P. was responsible for the genome assembly and bioinformatic analysis of WGS data. M.Q and Fr.B. were responsible for the analysis of SNP chip analysis. Q.D and A.W. contributed to the bioinformatic analysis of WGS data. I.B produced the PacBio assembly. M.B isolated HMW-DNA needed for long-read sequencing. Fl.B., C.K., D.B., R.L.-L., I.L. and K.A.G. contributed to the collection of sprat samples. M.E.P, L.A. and, M.Q. wrote the paper with input from all other authors. All authors approved the manuscript before submission.

## Data availability statement

The sequence data generated in this study and genome assemblies have been submitted to NCBI (https://www.ncbi.nlm.nih.gov/bioproject/PRJNA1023385). The analyses of data have been carried out with publicly available software and all are cited in the Methods section. Code associated with bioinformatic analyses are available at: https://github.com/LeifAnderssonLab/ Sprat_pool_reseq. Correspondence and requests for materials should be addressed to L.A.

