## Supplementary figures for "Limited parallelism in genetic adaptation to brackish water bodies in European sprat and Atlantic herring"

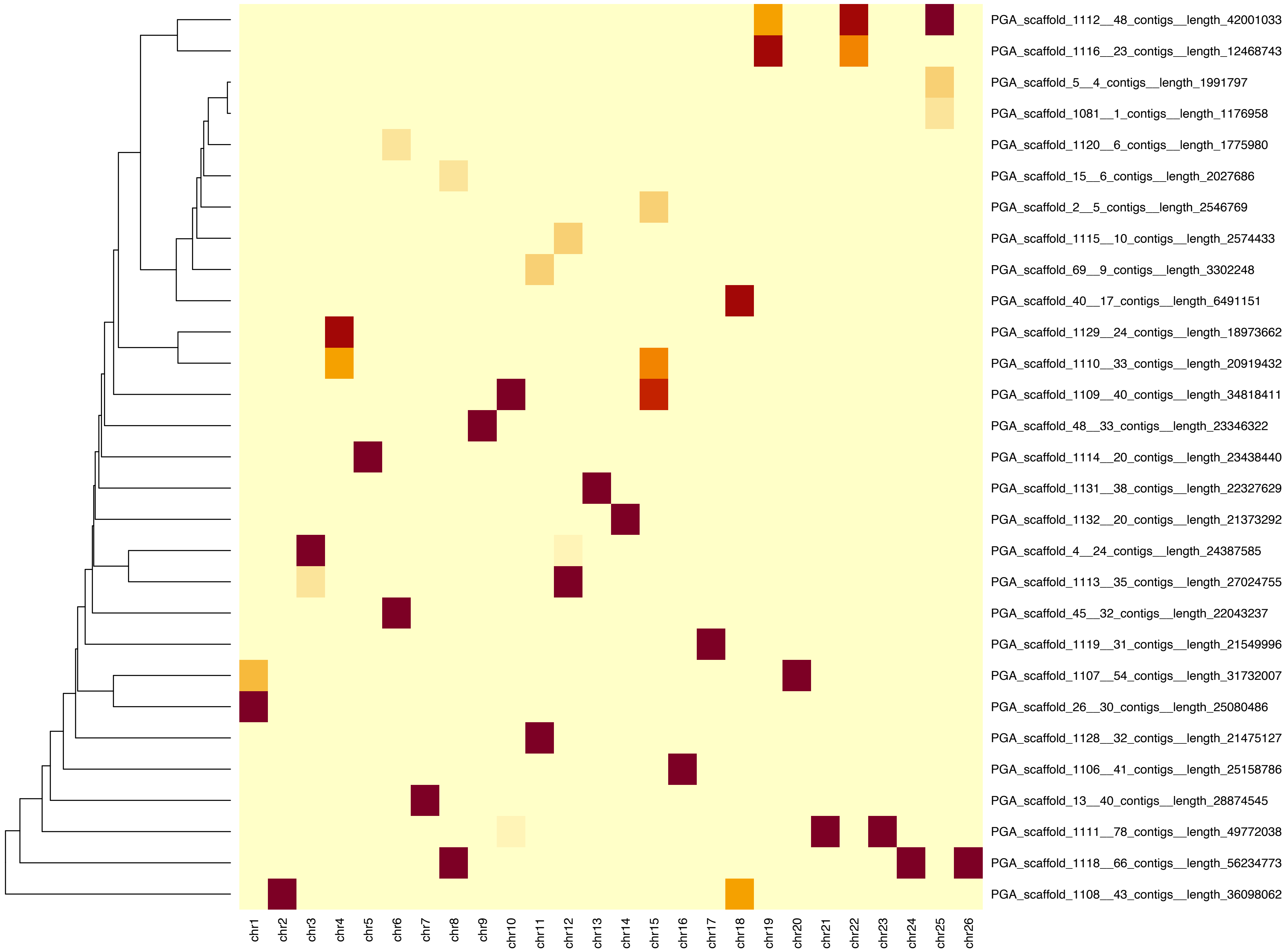


**Supplementary Fig. 1. Heatmap showing mapping intensities between European sprat scaffolds and Atlantic herring chromosomes.**

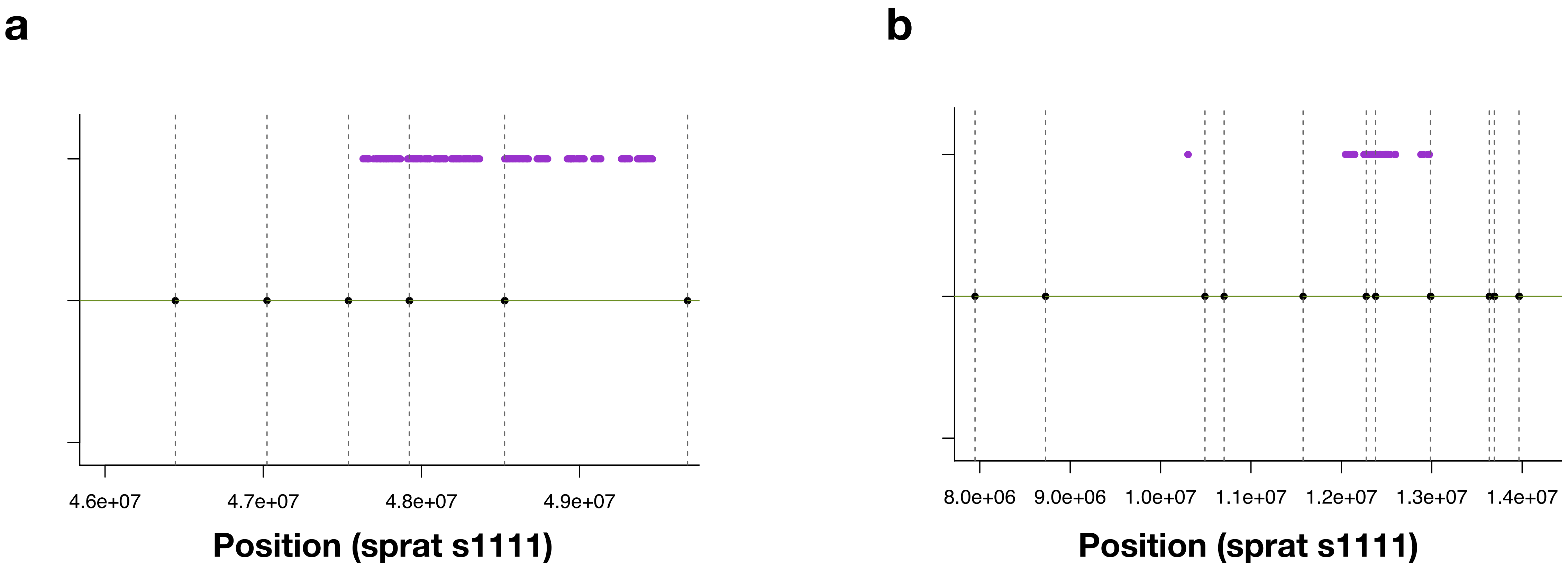


**Supplementary Fig. 3. Mappings of non-cognant regions.** The purple dots are lift-over based SNP locations from *C. harengus* Chr 21 (*a*) and Chr 23 (*b*), the horizontal green lines represent contigs from the underlying sprat PacBio assembly, while black dots and vertical dashed lines indicate scaffolding gaps.

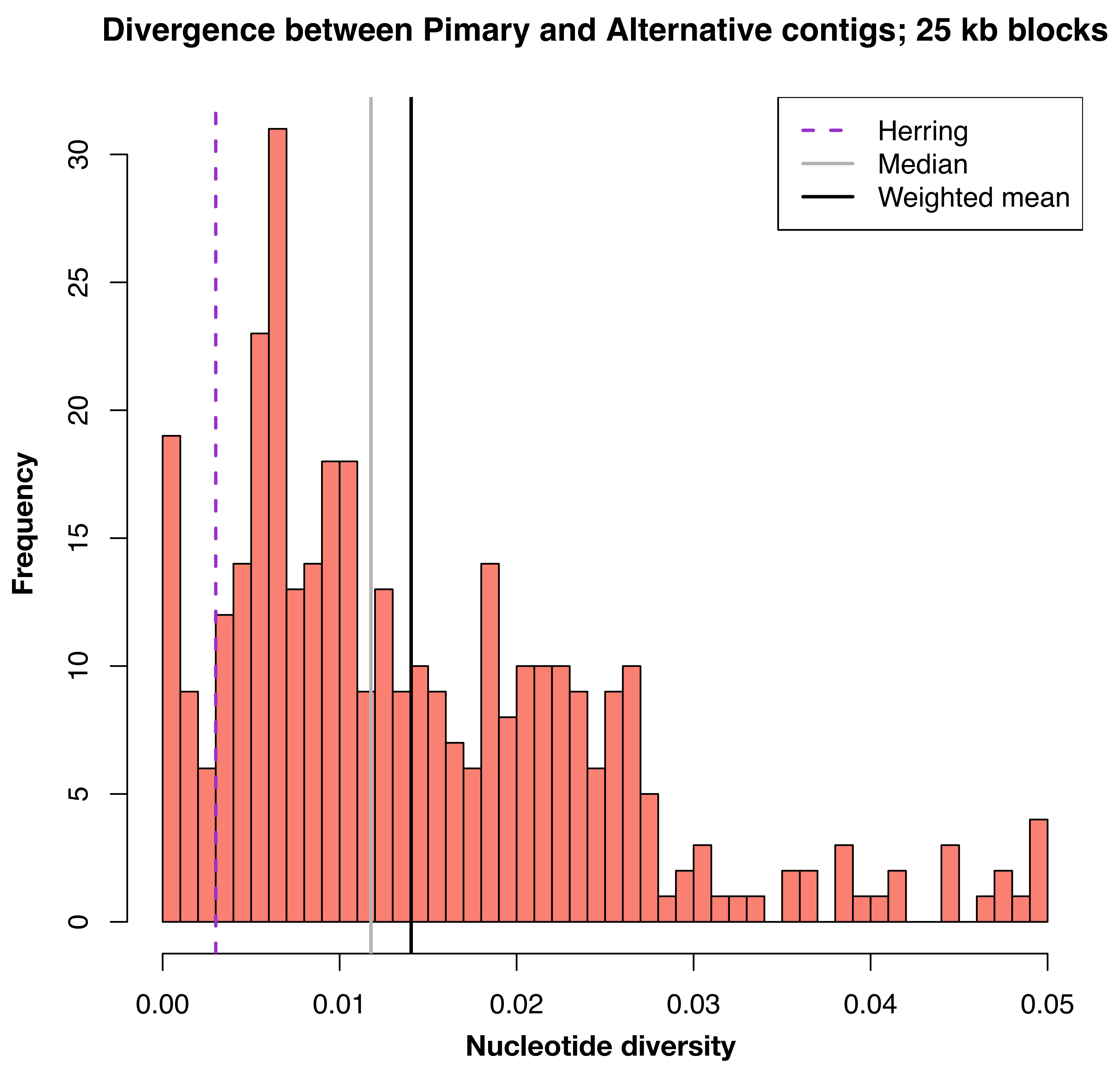


**Supplementary Fig. 4. Histogram over observed nucleotide diversity in randomly selected blocks across the genome.** The median, weighted mean and the corresponding value in Atlantic herring is highlighted by vertical lines.

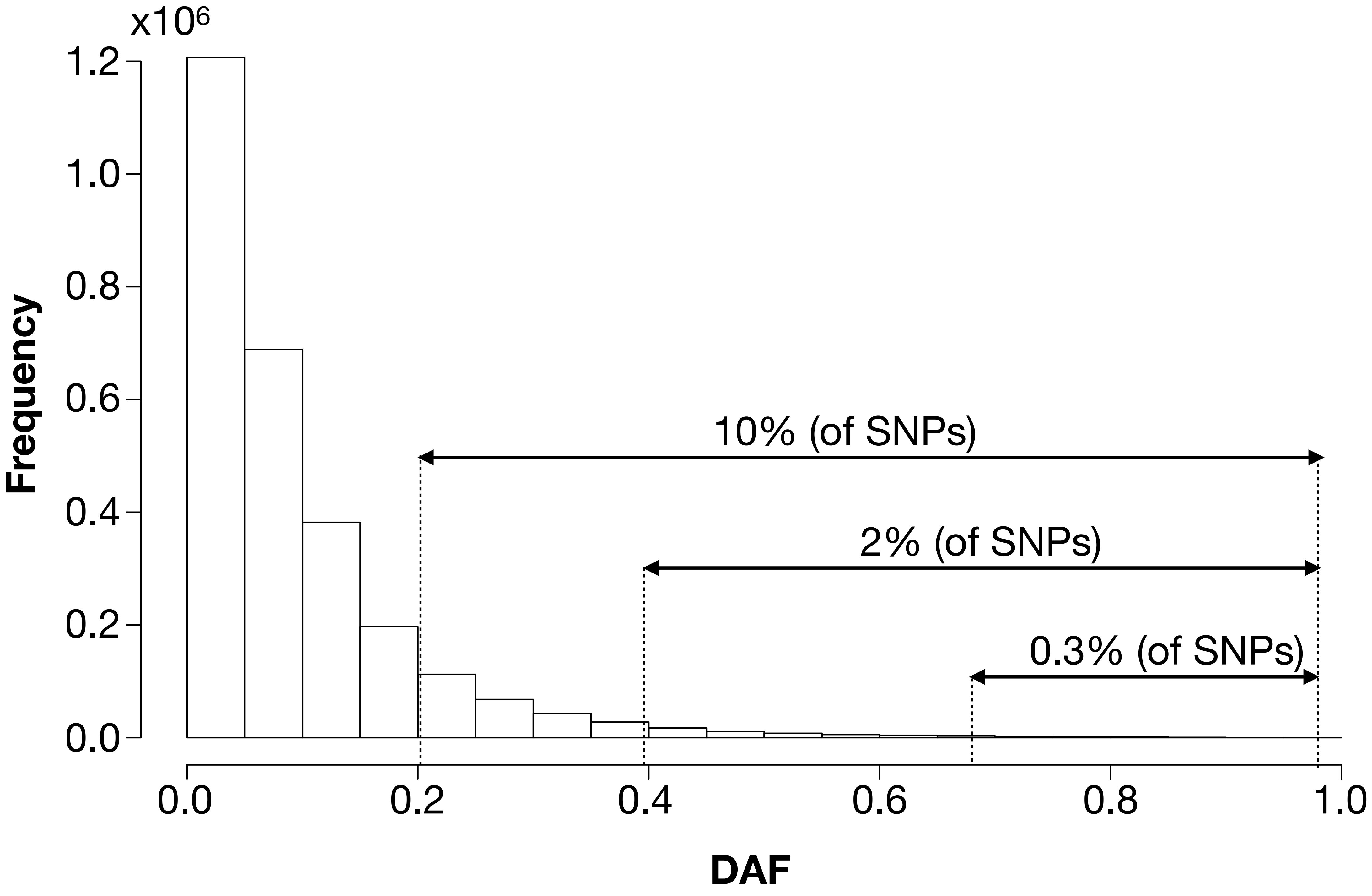


**Supplementary Fig. 5. Histogram of Delta Allele Frequencies (DAF) in the "Oceanic" vs "Brackish" contrast.** Arrows indicate proportions of SNPs within the respective DAF brackets.

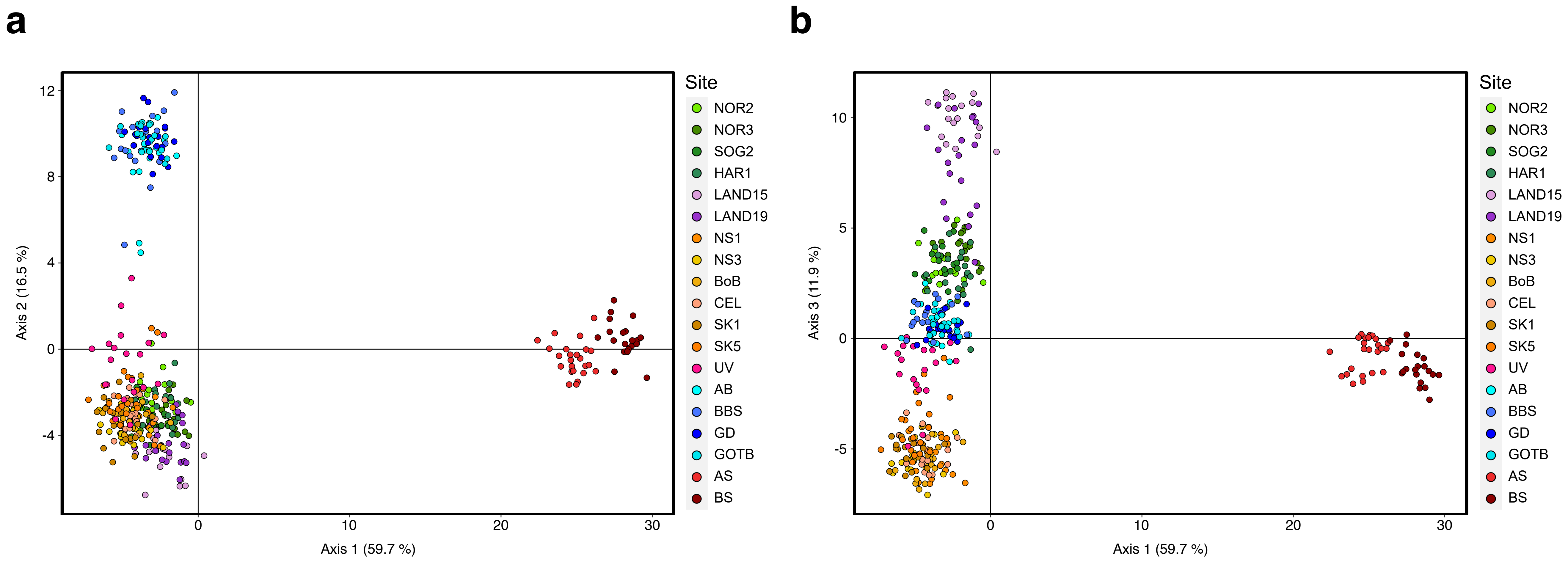


**Supplementary Fig. 6. Discriminant analysis of principle components (DAPC) of LD-pruned SNPs from the MultiFishChip.** Panels show axes 1 vs 2 (a) and 1 vs 3 (b), respectively. Sample codes are given in Supplementary Table 1.

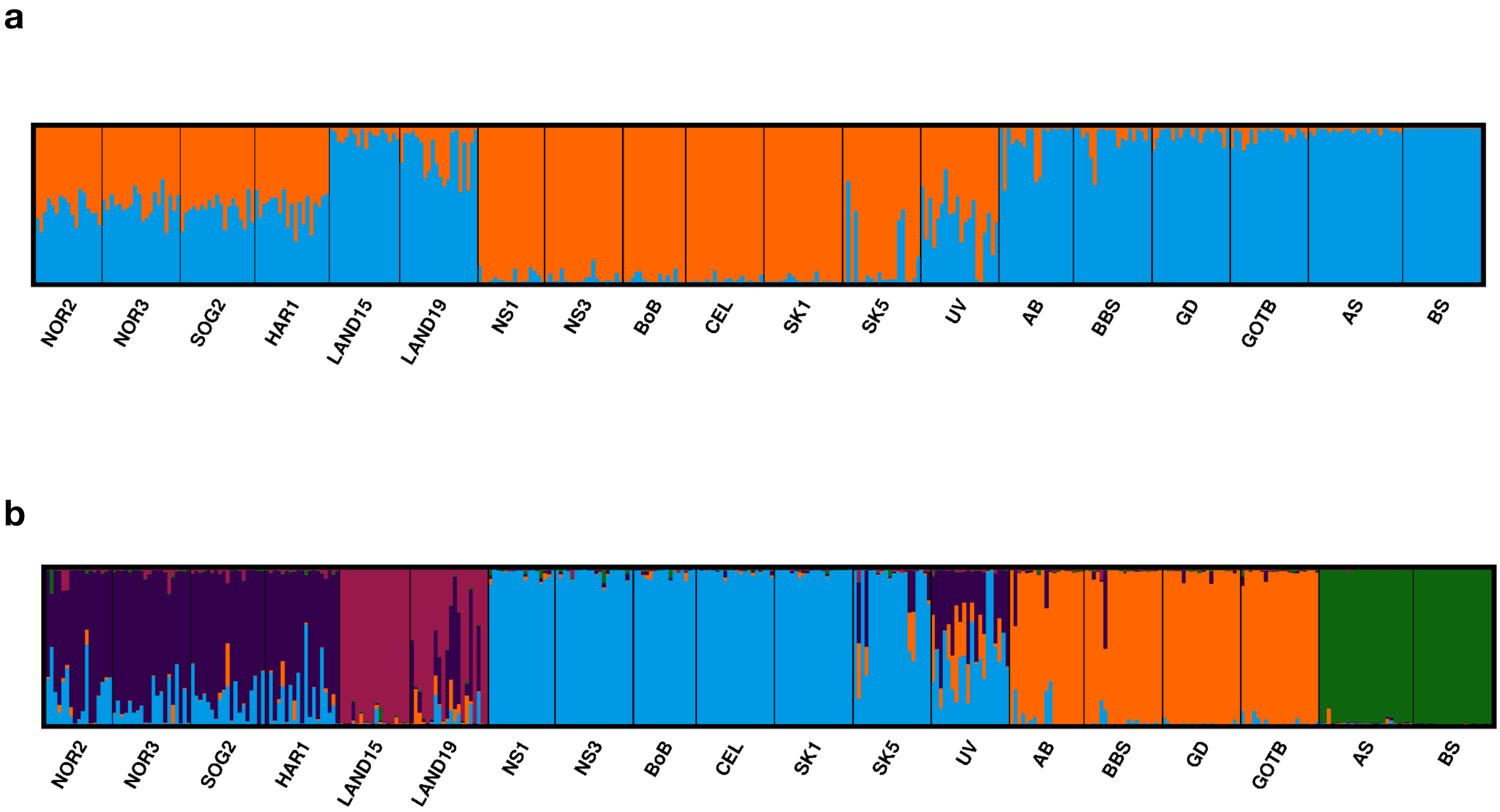


**Supplementary Fig. 7. Admixture analysis for 19 population samples of European sprat based on 2,063 SNPs.** STRUCTURE results for different values of *K*. a) *K* = 2; b) *K* = 5. Sample codes are given in Supplementary Table 1.

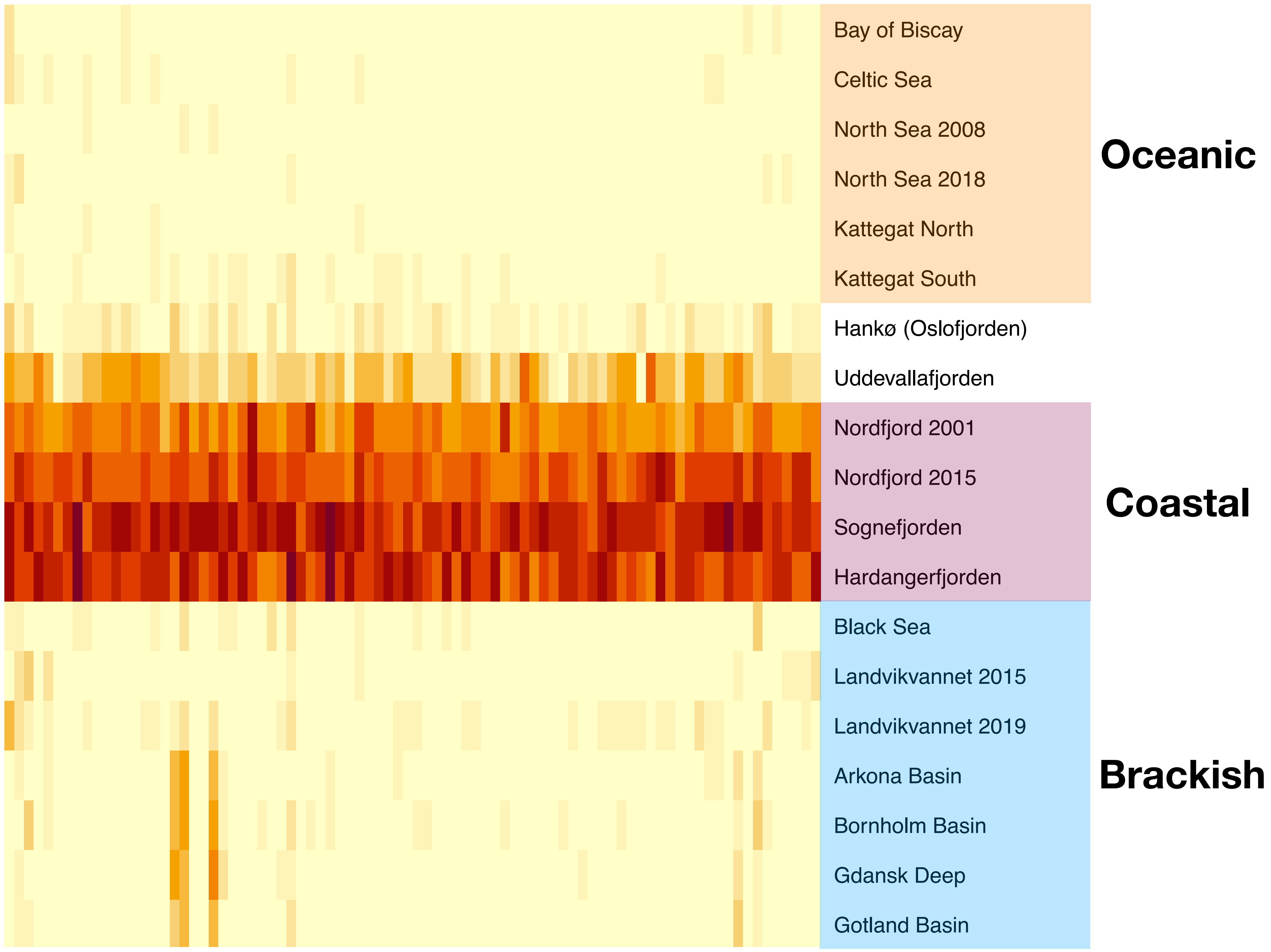


**Supplementary Fig. 8: Heatmap showing allele frequencies at the reginon corresponding to Chr 12 (0.1 -1.0 Mb) in the Atlantic herring.** Population groupings are indicated by labels and colored boxes, in accordance with Figure 2.

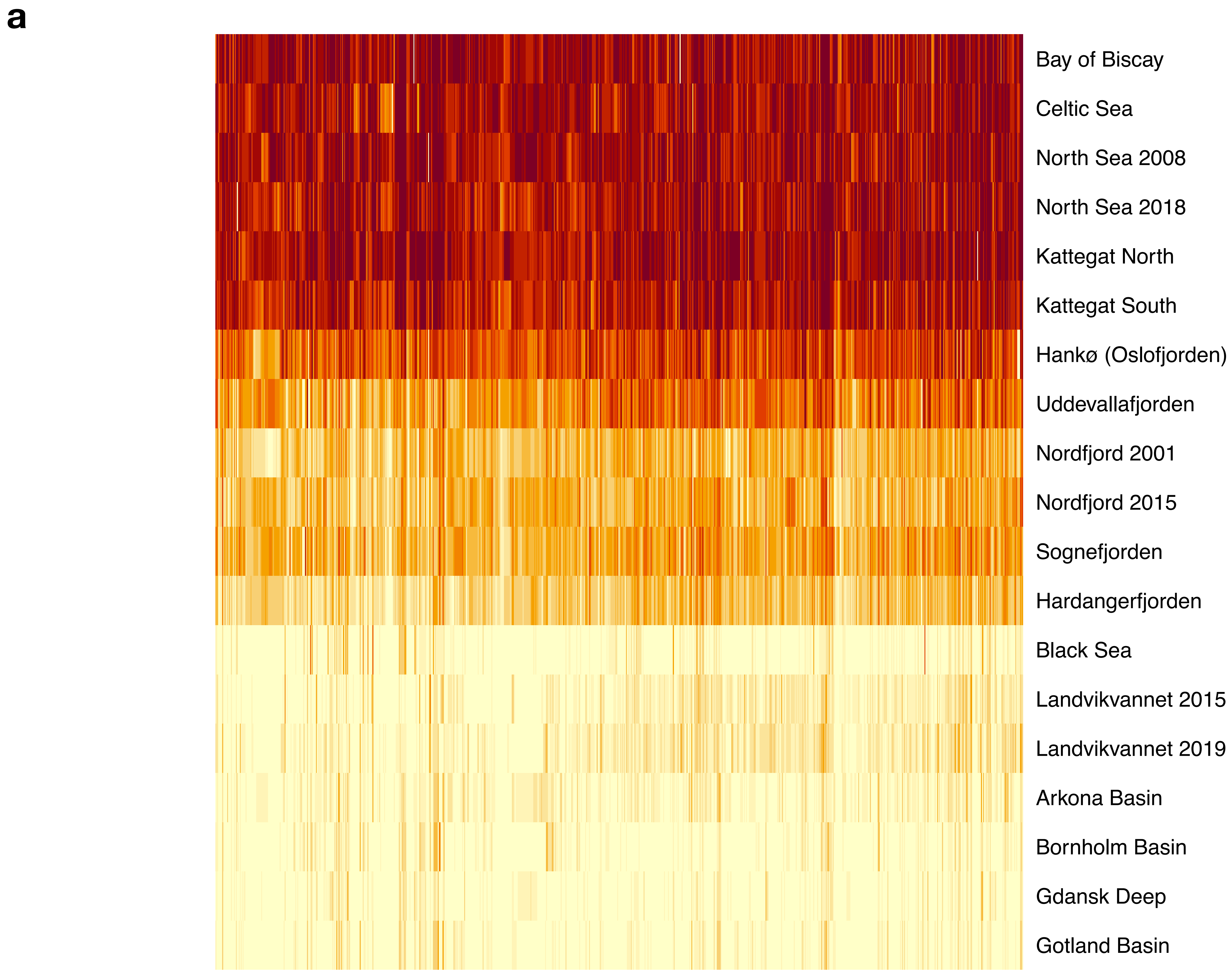


**Supplementary Fig. 9a. Heatmap showing allele frequencies at the inversion on scaffold s1110.**

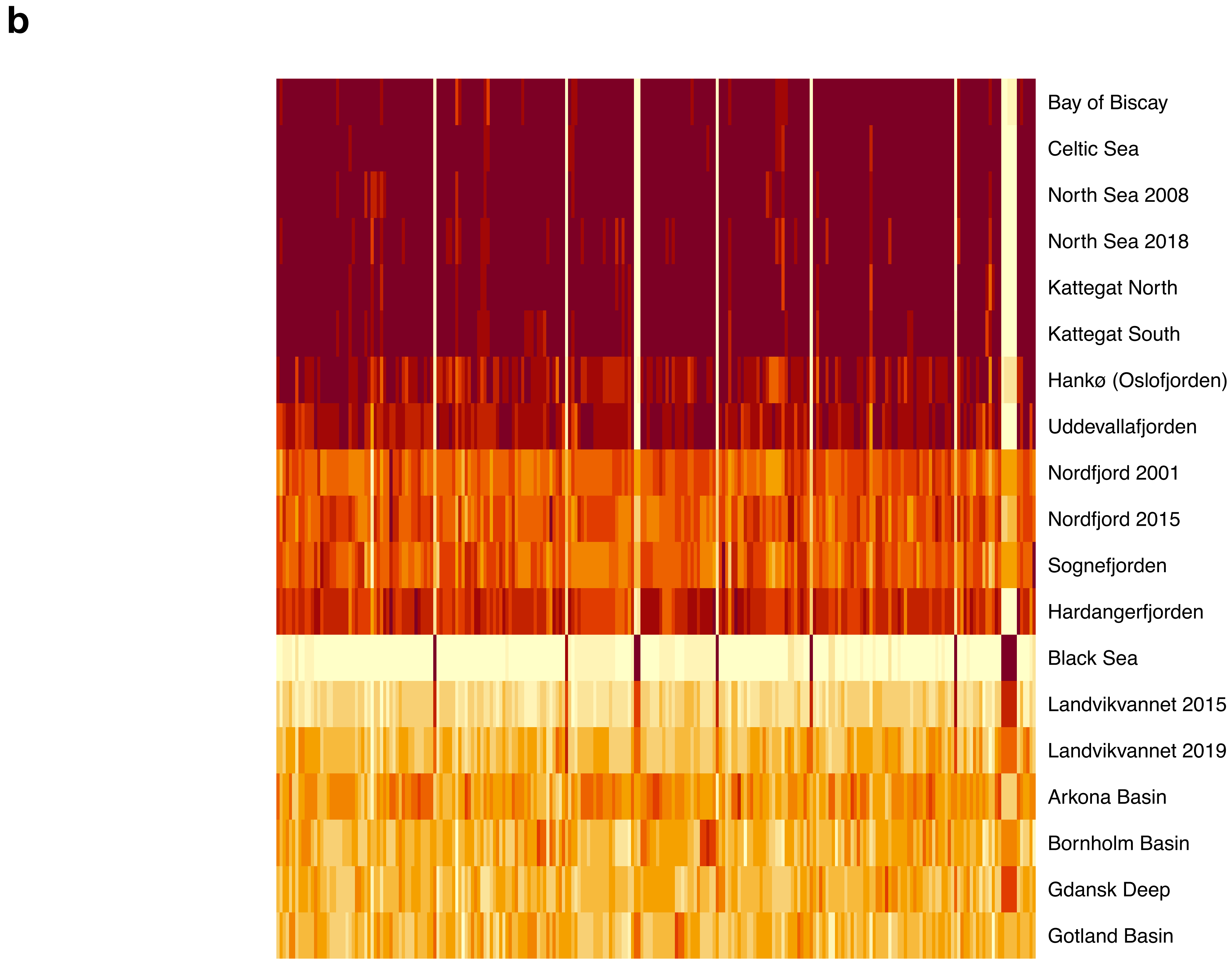


**Supplementary Fig. 9b. Heatmap showing allele frequencies at the inversion on scaffold s1114.**

**c**

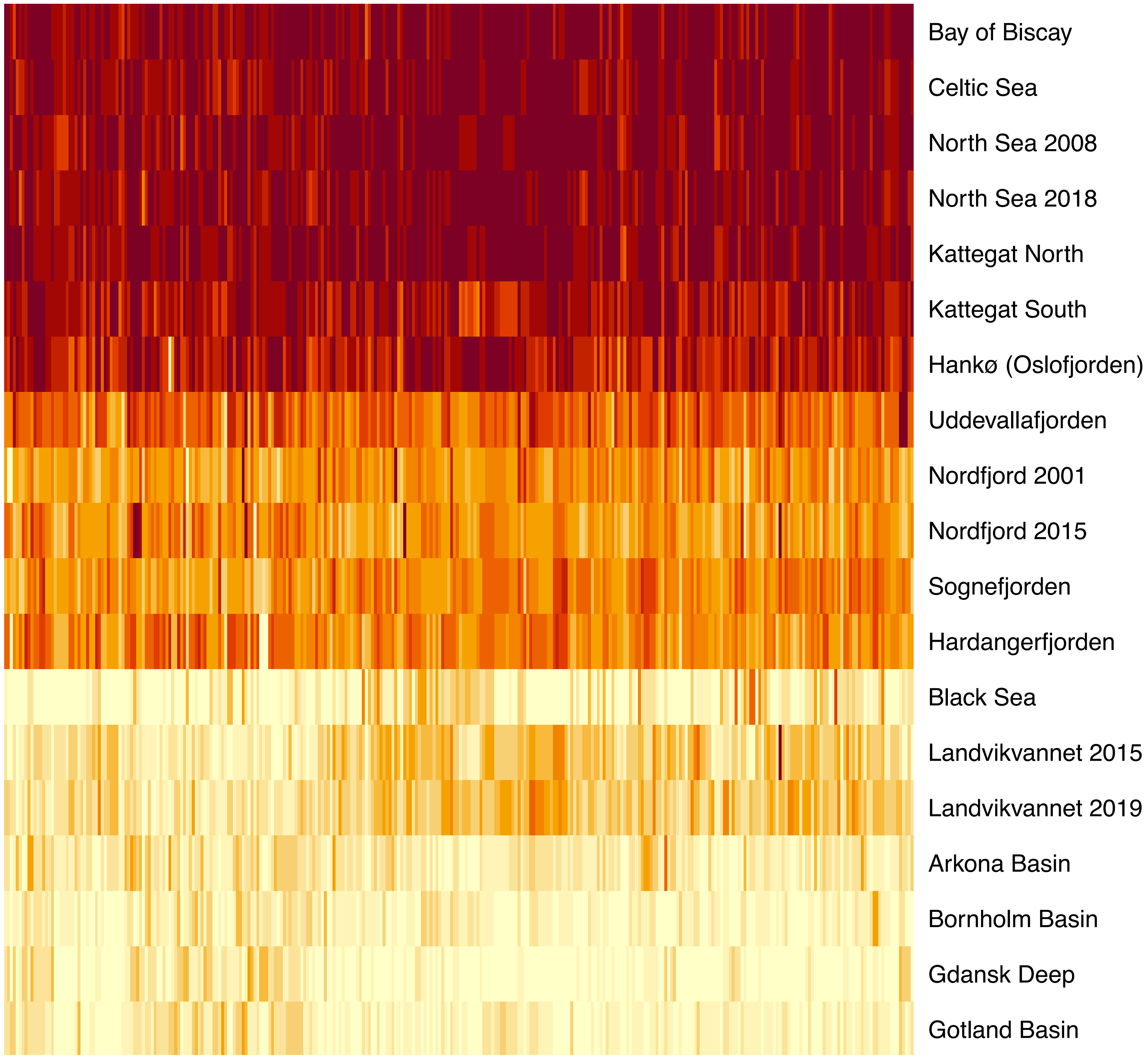


**Supplementary Fig. 9c. Heatmap showing allele frequencies at the inversion on scaffold s13.**

d

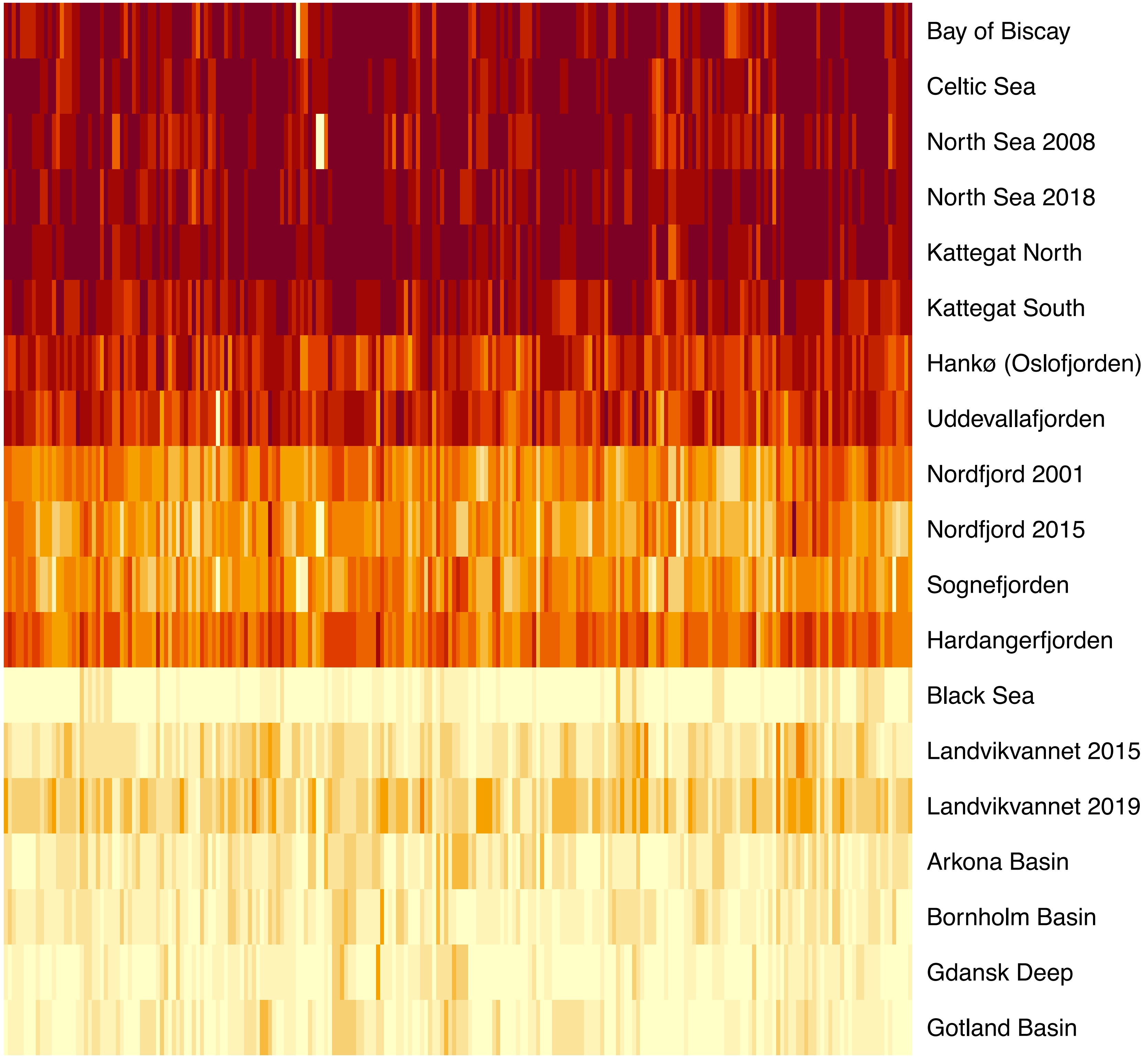


**Supplementary Fig. 9d. Heatmap showing allele frequencies at the inversion shared between scaffolds s173 and s374.**

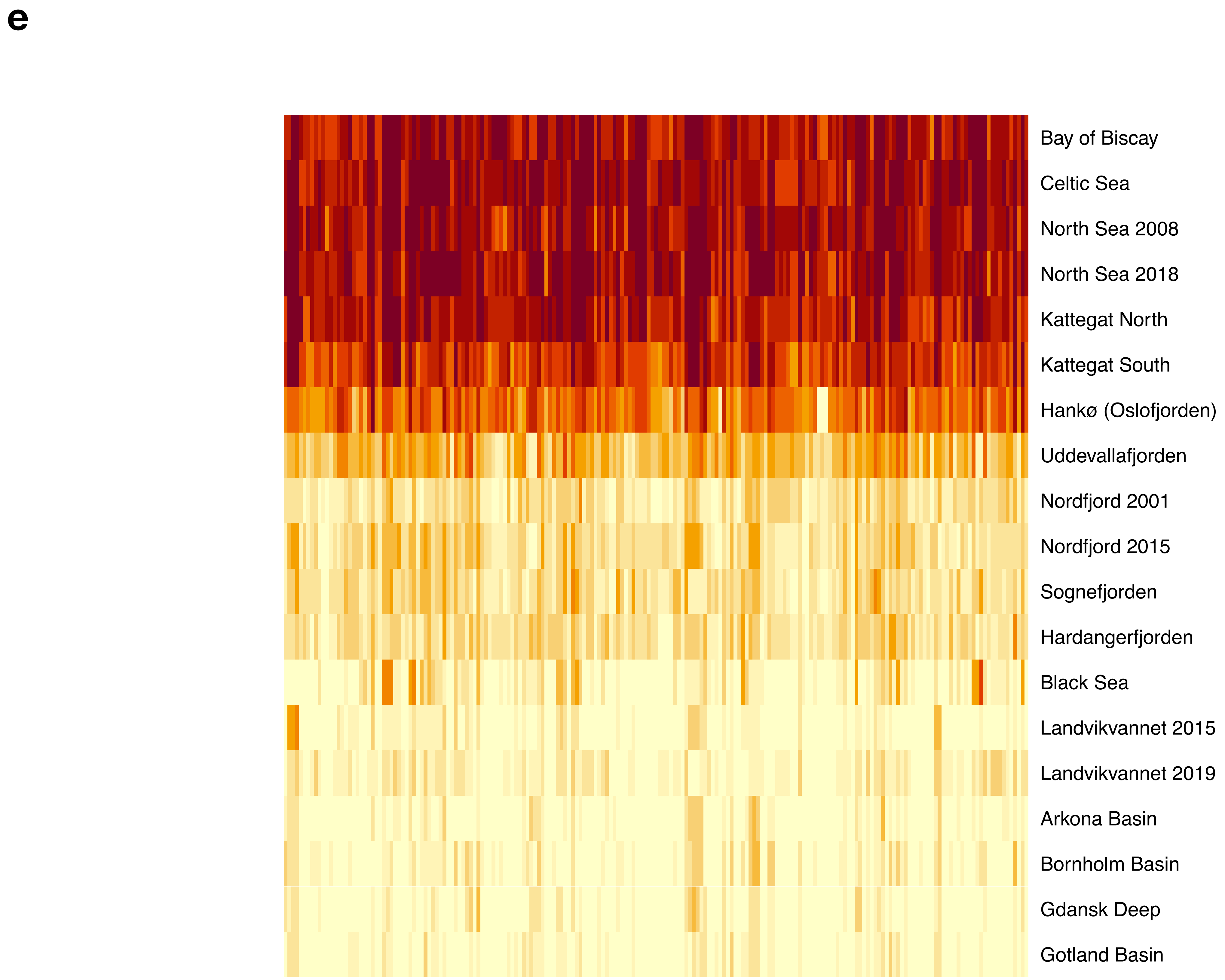


**Supplementary Fig. 9e. Heatmap showing allele frequencies at the inversion on scaffold s1111.**

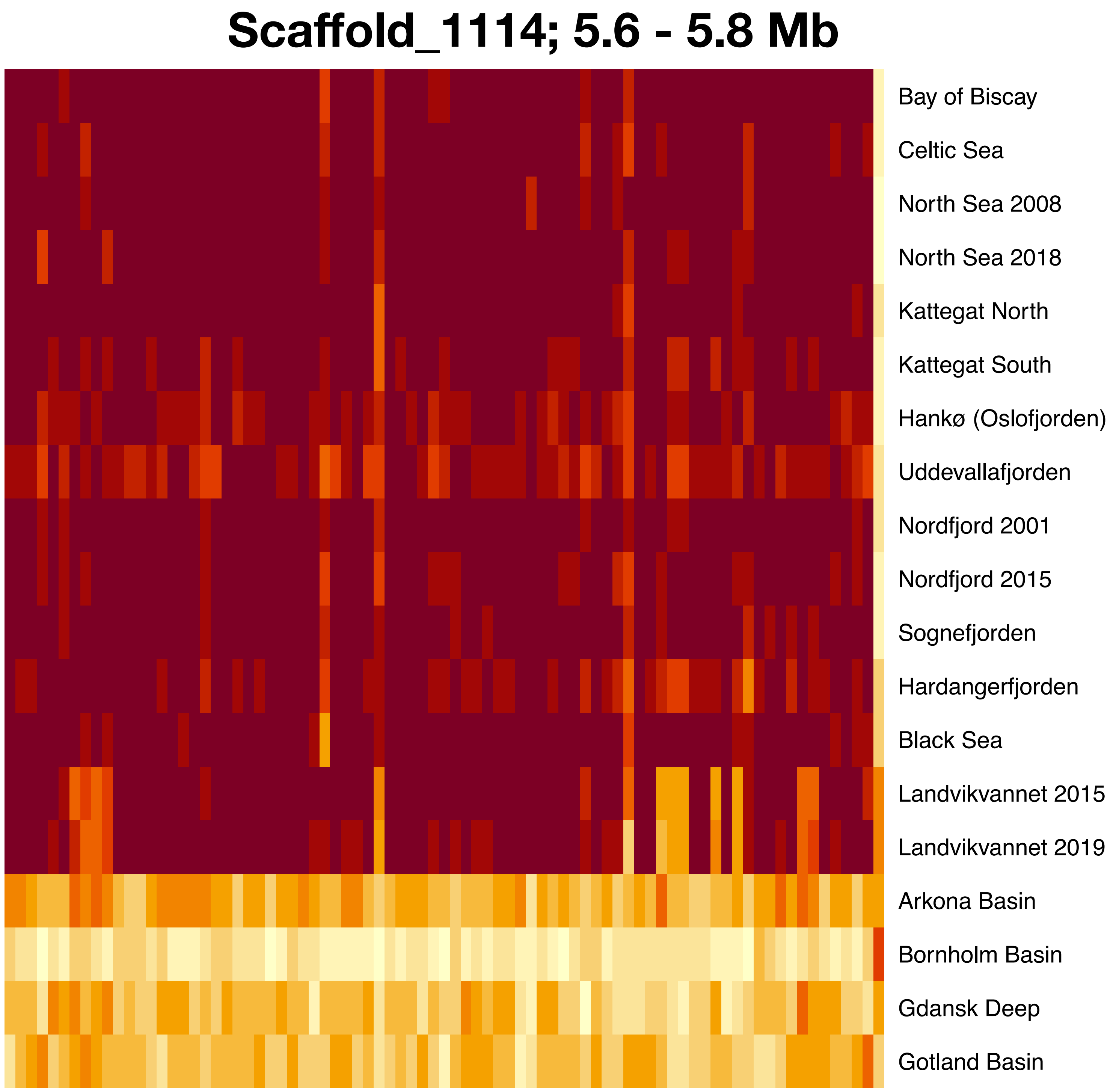


**Supplementary Fig. 10. Heatmap showing allele frequencies at a region of scaffold s1114.**

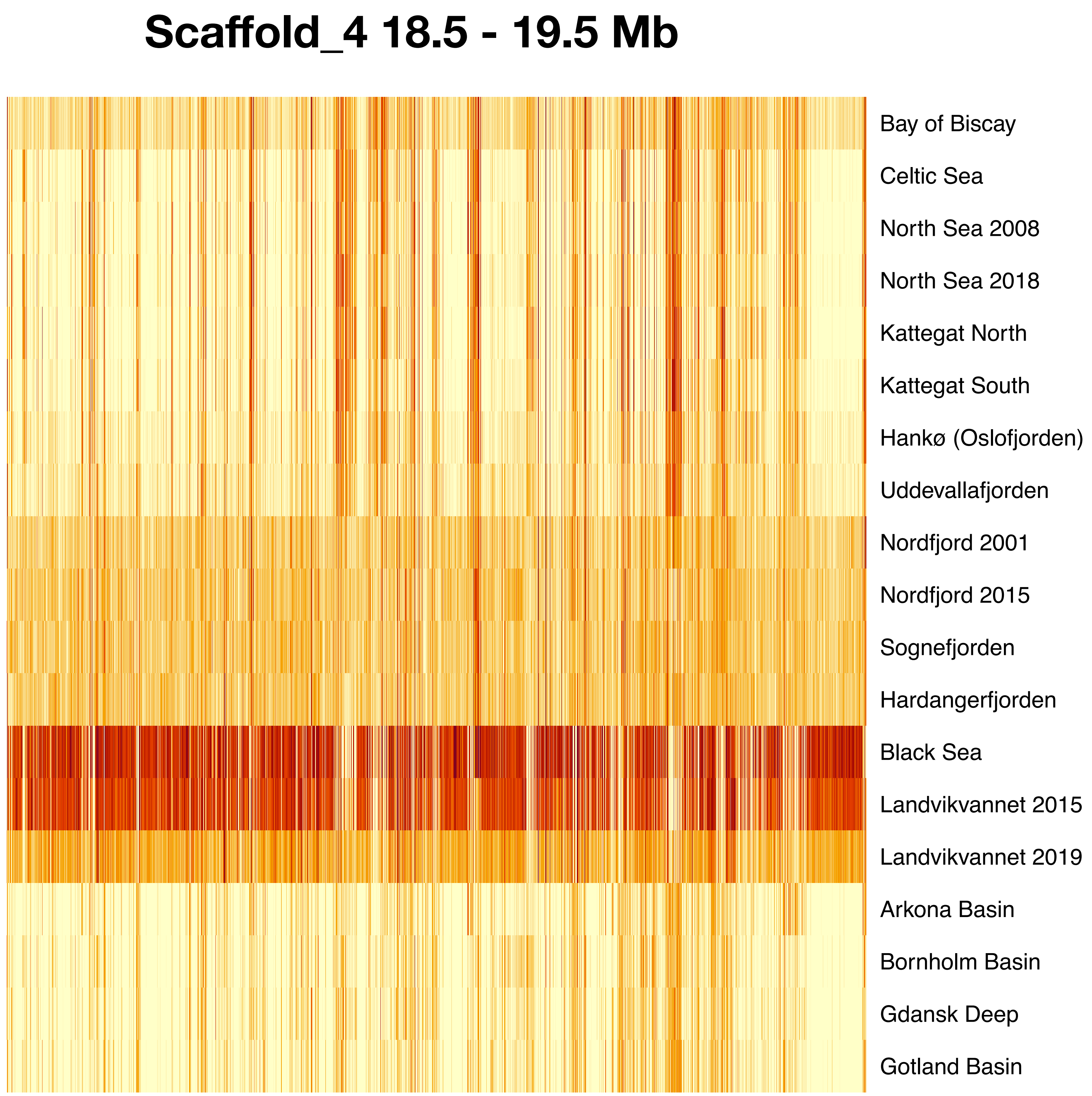


**Supplementary Fig. 11. Heatmap showing allele frequencies at a region of scaffold s4.**

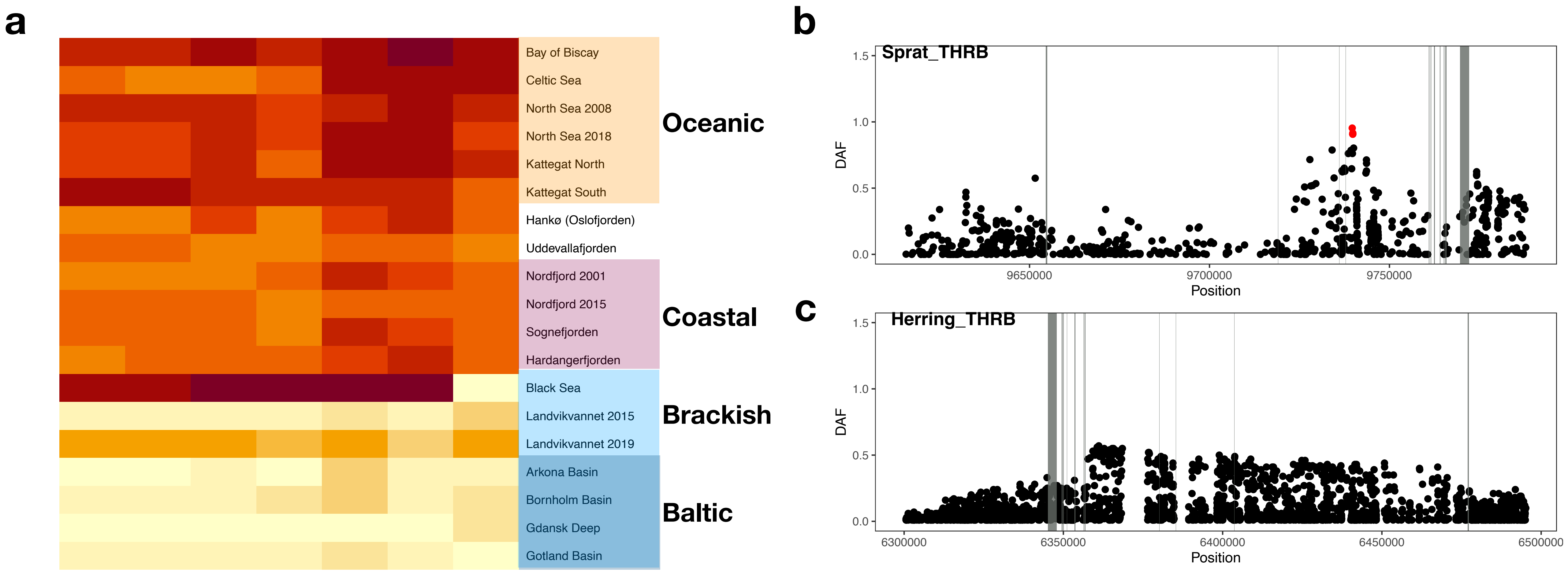


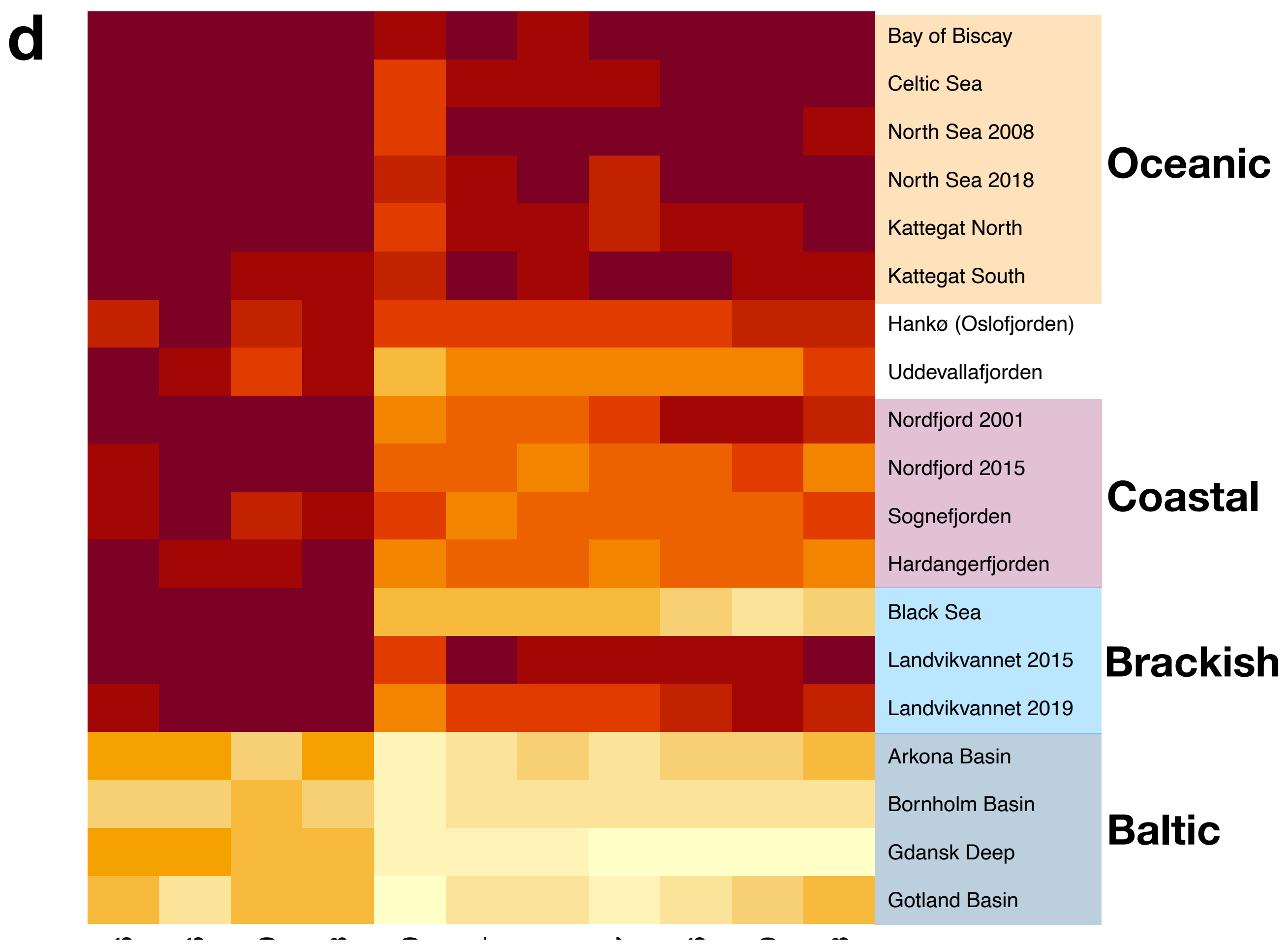


**Supplementary Figure 12. Genetic parallelism between European sprat and Atlantic herring – *THRB*.** *a*) Heatmap of the *THRB* region. Color coding of samples is as in Figure 2, with the exception of Baltic samples being highlighted within the Brackish group. *b*) Zoom-in showing DAF in the Oceanic and Brackish contrast, in European sprat, across the *THRB* locus (found on scaffold s1116 in the European sprat assembly). *c*) Zoom-in showing DAF between Atlantic and Baltic herring populations across the *THRB* locus on Chr 19 (4). In (*b*) and (*c*), grey boxes indicate exon locations. *d*) Heatmap of the *TNNI2* region. Color coding of samples is as in Figure 2, with the exception of Baltic samples being highlighted within the Brackish group.
