## Supplementary figure 2a for "Limited parallelism in genetic adaptation to brackish water bodies in European sprat and Atlantic herring"

Supplementary Fig. 2. Cross-mapping between major sprat scaffolds and C. harengus chromosomes. a) SNP liftover from sprat scaffolds to the two most represented herring chromosomes. b) as in (a), but in opposite direction.

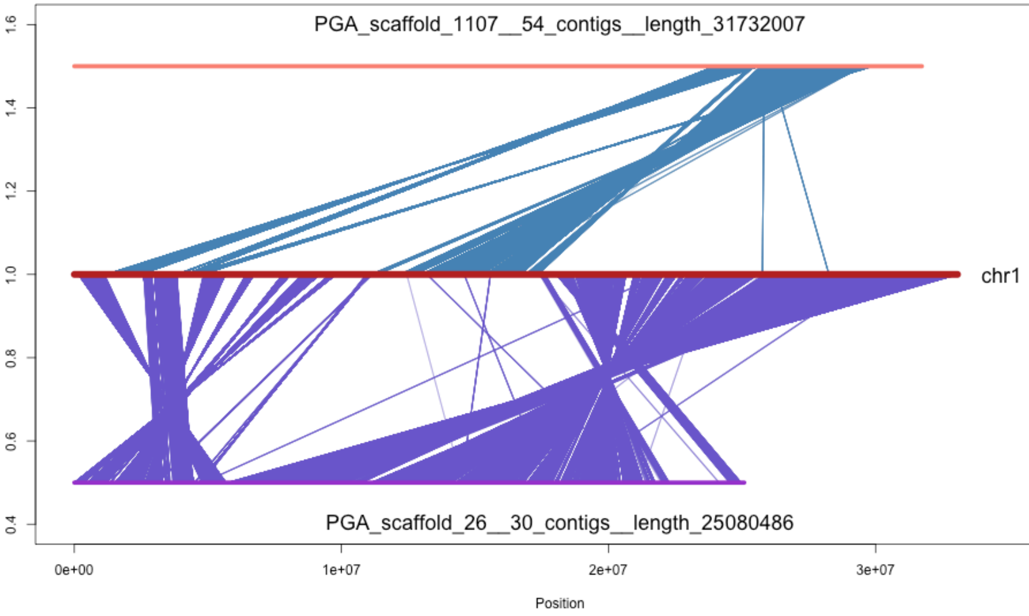
PGA_scaffold_1107__54_contigs__length_31732007
chr1
PGA_scaffold_26__30_contigs__length_25080486
1.6
1.4
1.2
1.0
0.8
0.6
0.4
0e+00
1e+07
2e+07
3e+07
Position

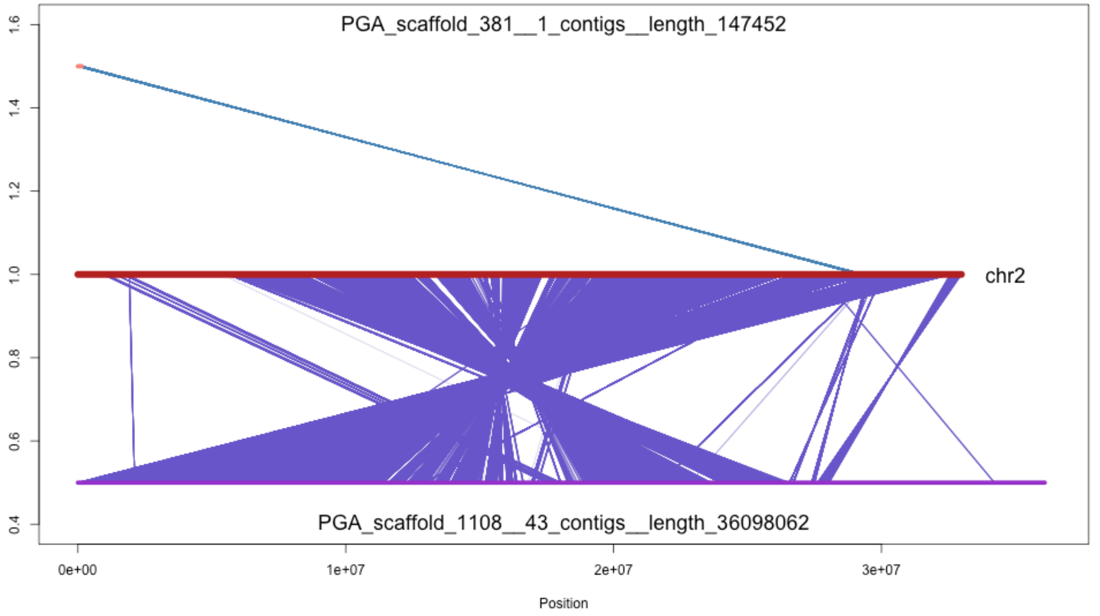
PGA_scaffold_381__1_contigs__length_147452
chr2
PGA_scaffold_1108__43_contigs__length_36098062
1.6
1.4
1.2
1.0
0.8
0.6
0.4
0e+00
1e+07
2e+07
3e+07
Position

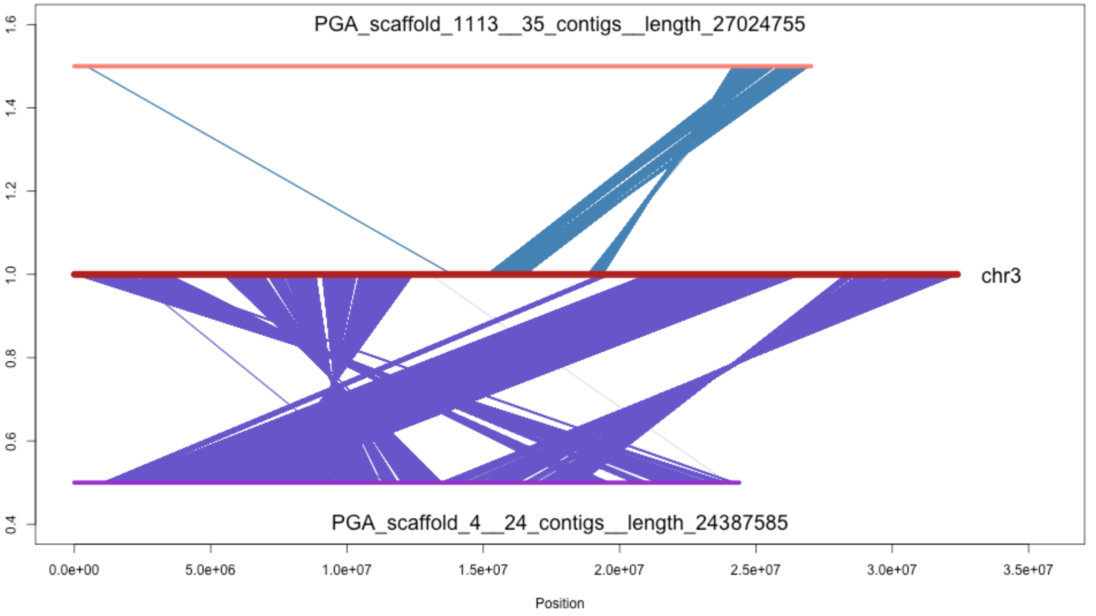

PGA_scaffold_1113__35_contigs__length_27024755
chr3
PGA_scaffold_4__24_contigs__length_24387585
1.6
1.4
1.2
1.0
0.8
0.6
0.4
0.0e+00
5.0e+06
1.0e+07
1.5e+07
2.0e+07
2.5e+07
3.0e+07
3.5e+07
Position

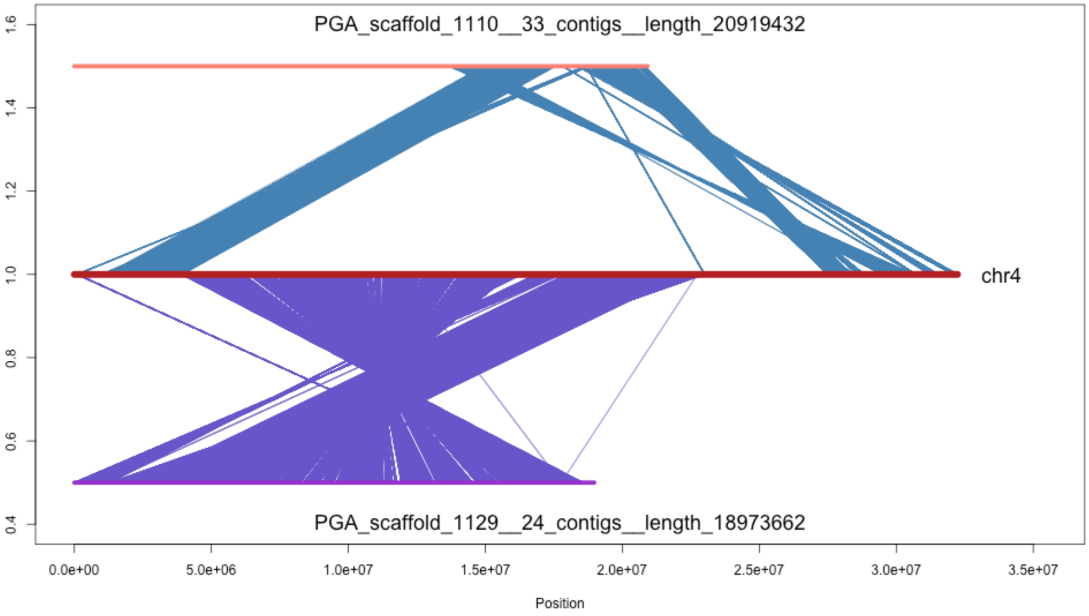
PGA_scaffold_1110__33_contigs__length_20919432
chr4
PGA_scaffold_1129__24_contigs__length_18973662
1.6
1.4
1.2
1.0
0.8
0.6
0.4
0.0e+00
5.0e+06
1.0e+07
1.5e+07
2.0e+07
2.5e+07
3.0e+07
3.5e+07
Position

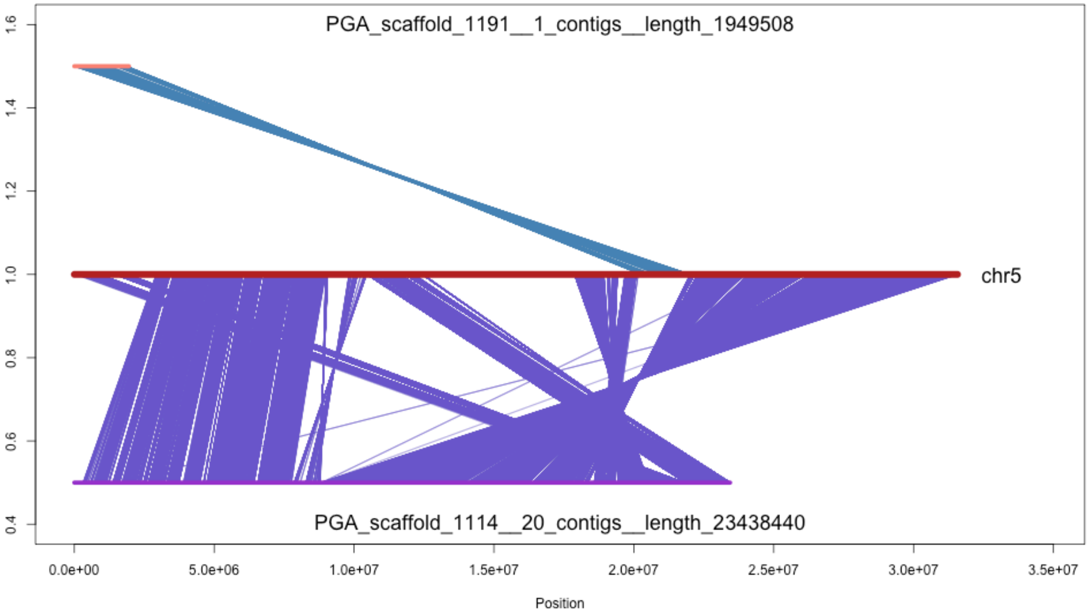

PGA_scaffold_1191__1_contigs__length_1949508
chr5
PGA_scaffold_1114__20_contigs__length_23438440
1.6
1.4
1.2
1.0
0.8
0.6
0.4
0.0e+00
5.0e+06
1.0e+07
1.5e+07
2.0e+07
2.5e+07
3.0e+07
3.5e+07
Position

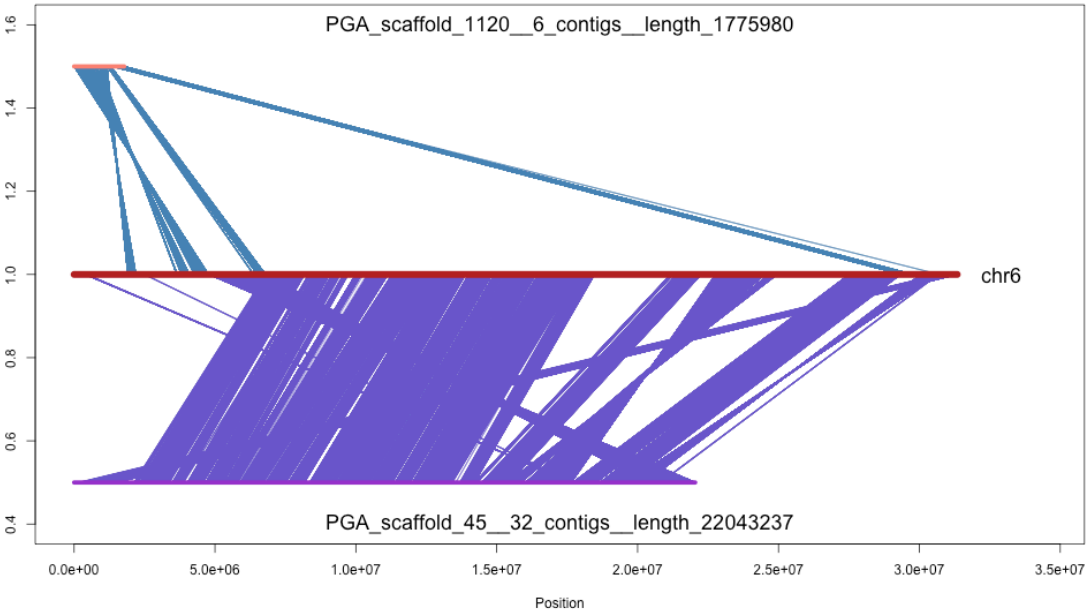
PGA_scaffold_1120__6_contigs__length_1775980
chr6
PGA_scaffold_45__32_contigs__length_22043237
1.6
1.4
1.2
1.0
0.8
0.6
0.4
0.0e+00
5.0e+06
1.0e+07
1.5e+07
2.0e+07
2.5e+07
3.0e+07
3.5e+07
Position

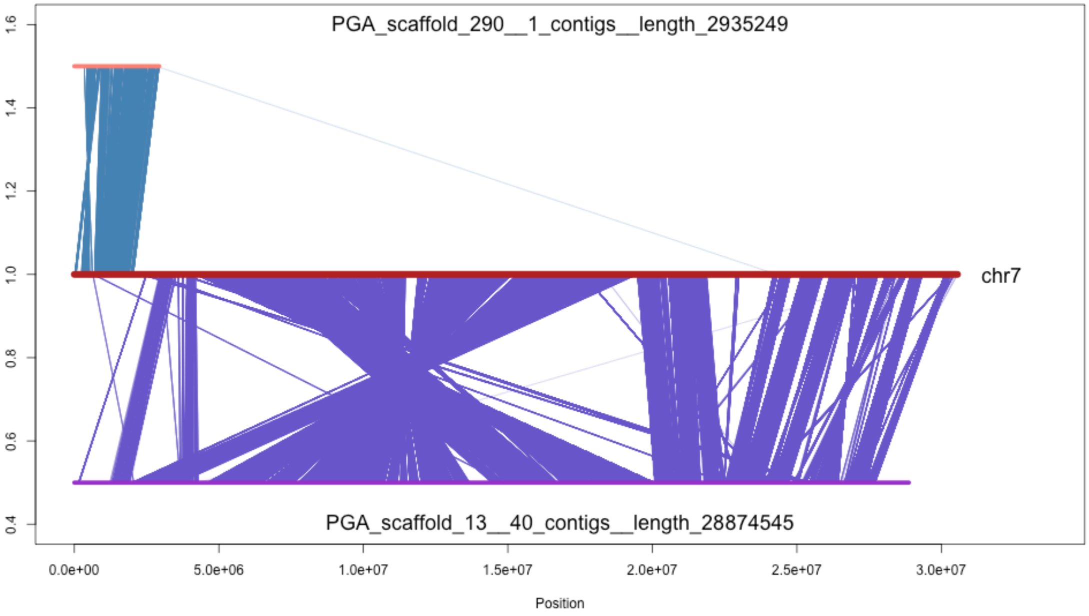
PGA_scaffold_290__1_contigs__length_2935249
chr7
PGA_scaffold_13__40_contigs__length_28874545
1.6
1.4
1.2
1.0
0.8
0.6
0.4
0.0e+00
5.0e+06
1.0e+07
1.5e+07
2.0e+07
2.5e+07
3.0e+07
Position

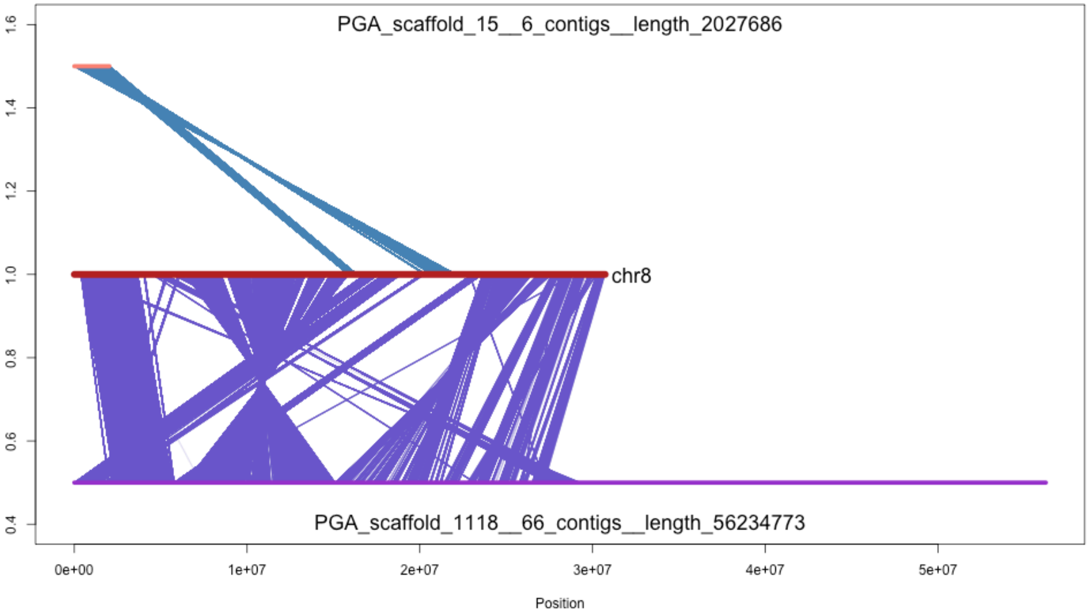
PGA_scaffold_15__6_contigs__length_2027686
chr8
PGA_scaffold_1118__66_contigs__length_56234773
1.6
1.4
1.2
1.0
0.8
0.6
0.4
0e+00
1e+07
2e+07
3e+07
4e+07
5e+07
Position

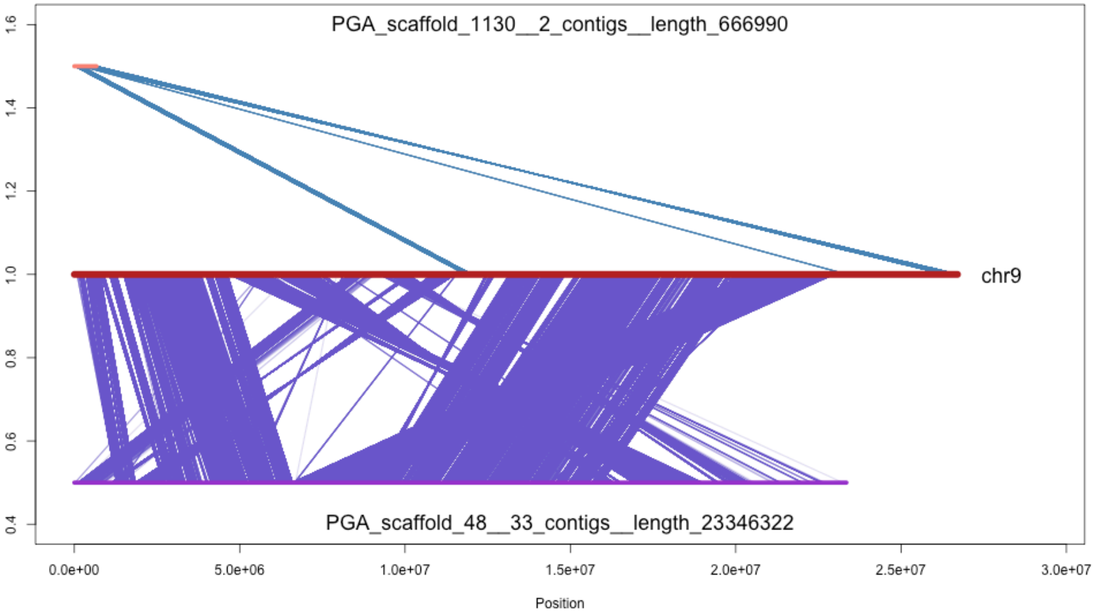

PGA_scaffold_1130__2_contigs__length_666990
chr9
PGA_scaffold_48__33_contigs__length_23346322
1.6
1.4
1.2
1.0
0.8
0.6
0.4
0.0e+00
5.0e+06
1.0e+07
1.5e+07
2.0e+07
2.5e+07
3.0e+07
Position

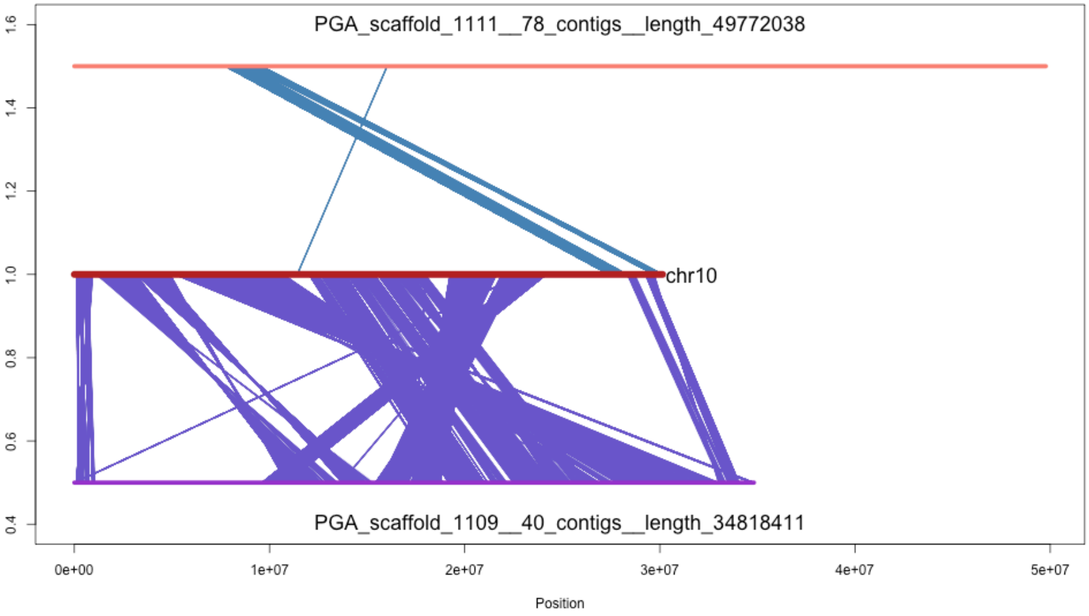

PGA_scaffold_1111__78_contigs__length_49772038
chr10
PGA_scaffold_1109__40_contigs__length_34818411
0.4
0.6
0.8
1.0
1.2
1.4
1.6
0e+00
1e+07
2e+07
3e+07
4e+07
5e+07
Position

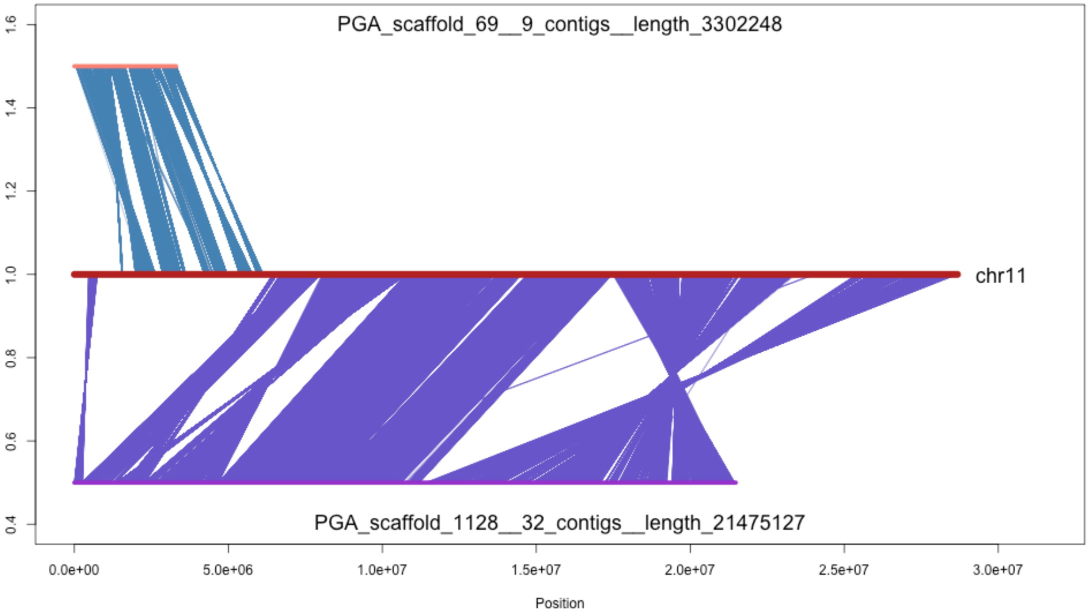

PGA_scaffold_69__9_contigs__length_3302248
chr11
PGA_scaffold_1128__32_contigs__length_21475127
1.6
1.4
1.2
1.0
0.8
0.6
0.4
0.0e+00
5.0e+06
1.0e+07
1.5e+07
2.0e+07
2.5e+07
3.0e+07
Position

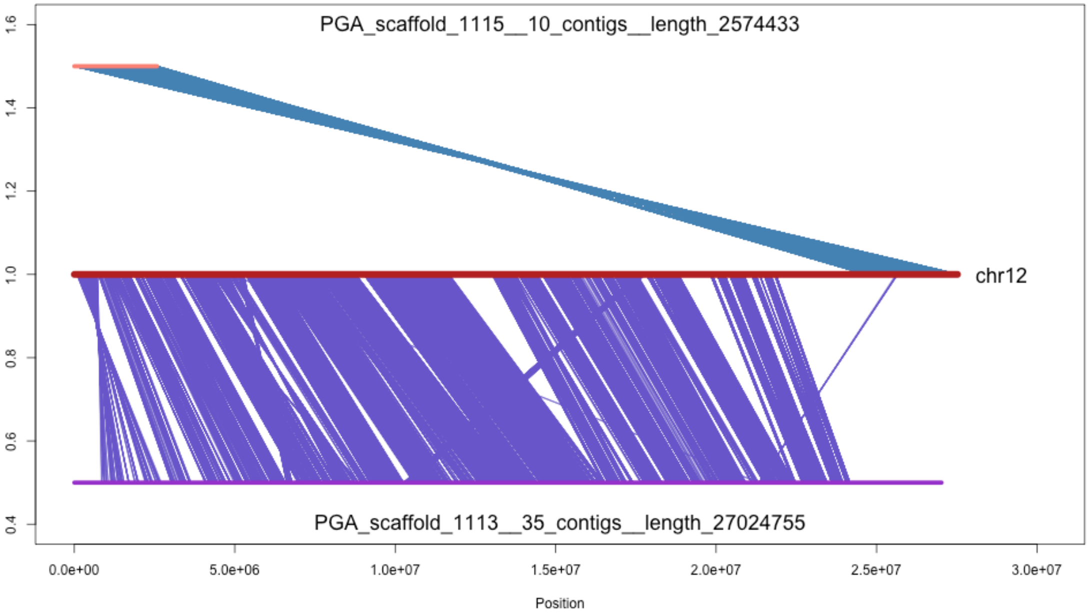

PGA_scaffold_1115__10_contigs__length_2574433
chr12
PGA_scaffold_1113__35_contigs__length_27024755
1.6
1.4
1.2
1.0
0.8
0.6
0.4
0.0e+00
5.0e+06
1.0e+07
1.5e+07
2.0e+07
2.5e+07
3.0e+07
Position

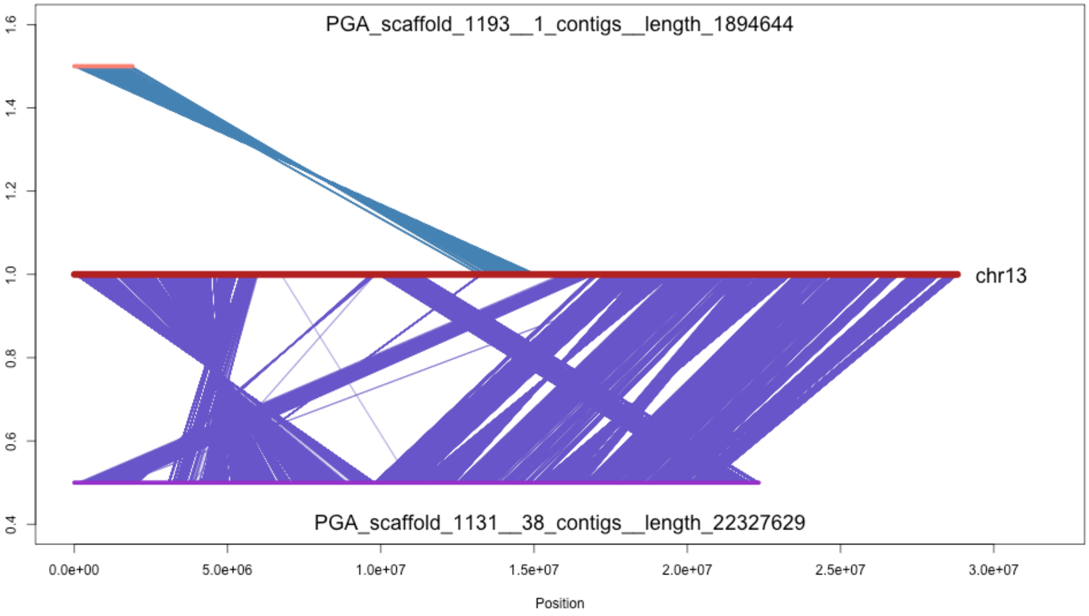
PGA_scaffold_1193__1_contigs__length_1894644
chr13
PGA_scaffold_1131__38_contigs__length_22327629
1.6
1.4
1.2
1.0
0.8
0.6
0.4
0.0e+00
5.0e+06
1.0e+07
1.5e+07
2.0e+07
2.5e+07
3.0e+07
Position

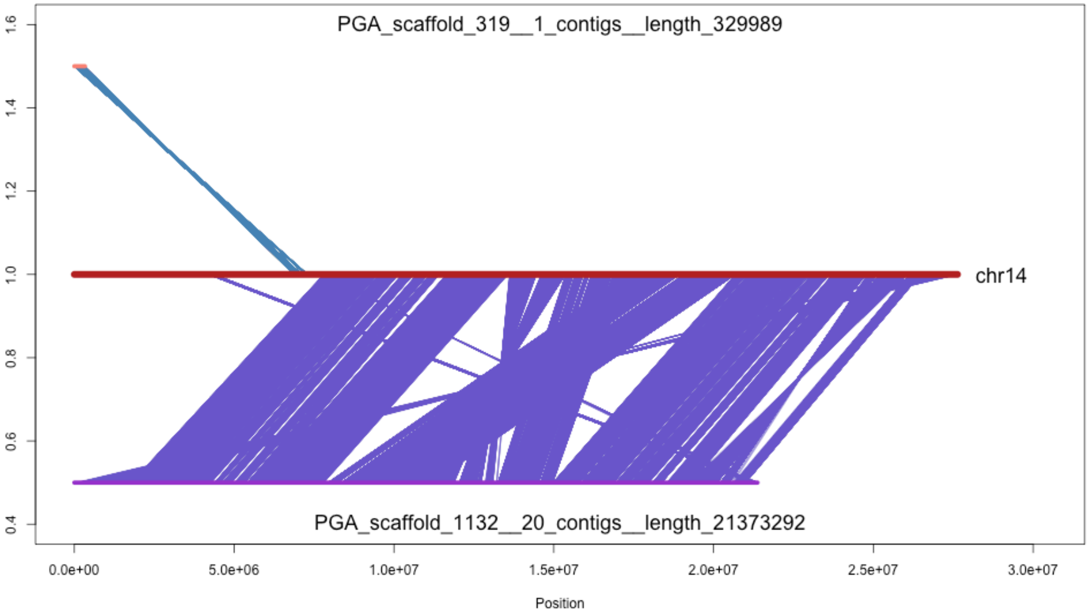
PGA_scaffold_319__1_contigs__length_329989
chr14
PGA_scaffold_1132__20_contigs__length_21373292
1.6
1.4
1.2
1.0
0.8
0.6
0.4
0.0e+00
5.0e+06
1.0e+07
1.5e+07
2.0e+07
2.5e+07
3.0e+07
Position

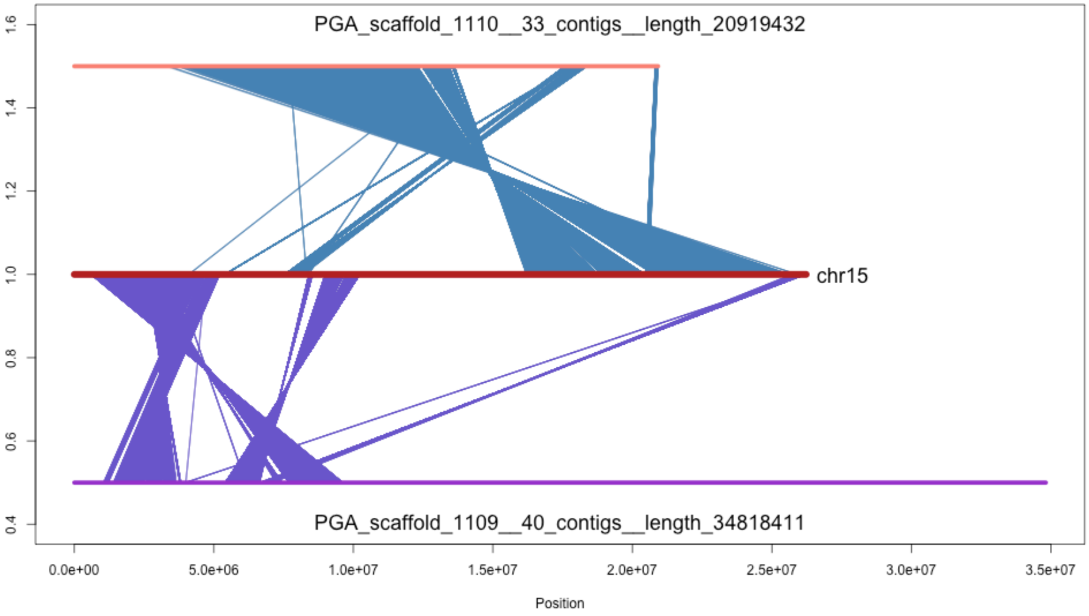
PGA_scaffold_1110__33_contigs__length_20919432
chr15
PGA_scaffold_1109__40_contigs__length_34818411
0.4
0.6
0.8
1.0
1.2
1.4
1.6
0.0e+00
5.0e+06
1.0e+07
1.5e+07
2.0e+07
2.5e+07
3.0e+07
3.5e+07
Position

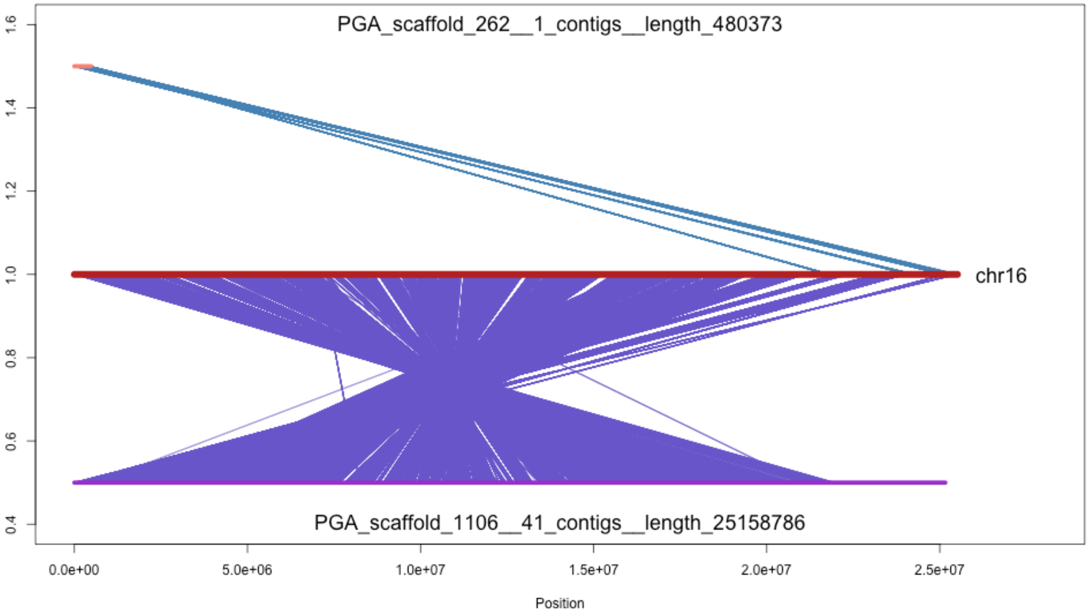

PGA_scaffold_262__1_contigs__length_480373
chr16
PGA_scaffold_1106__41_contigs__length_25158786
1.6
1.4
1.2
1.0
0.8
0.6
0.4
0.0e+00
5.0e+06
1.0e+07
1.5e+07
2.0e+07
2.5e+07
Position

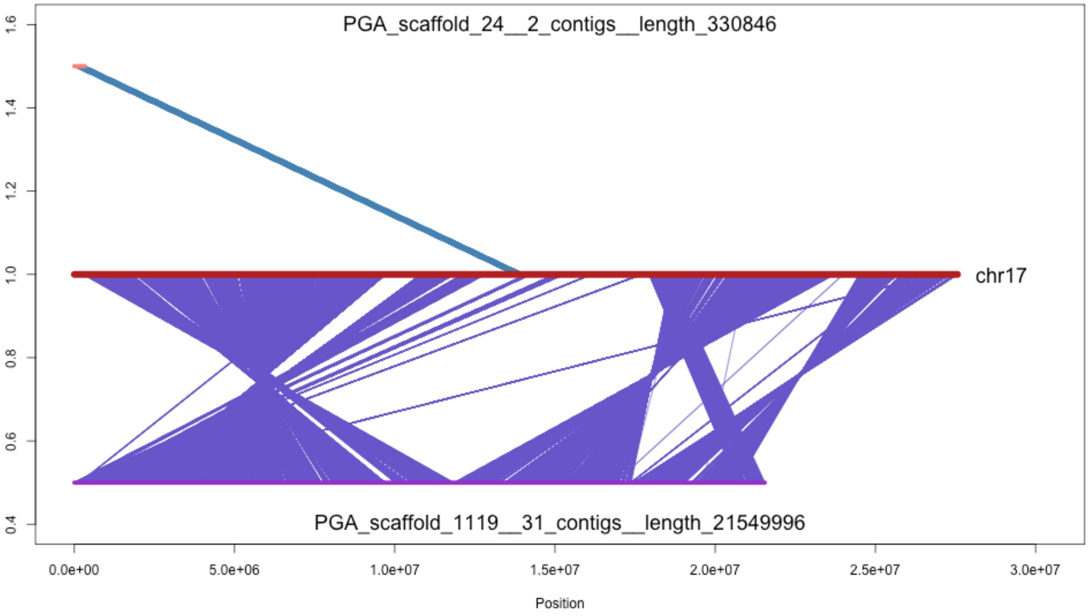
PGA_scaffold_24__2_contigs__length_330846
chr17
PGA_scaffold_1119__31_contigs__length_21549996
1.6
1.4
1.2
1.0
0.8
0.6
0.4
0.0e+00
5.0e+06
1.0e+07
1.5e+07
2.0e+07
2.5e+07
3.0e+07
Position

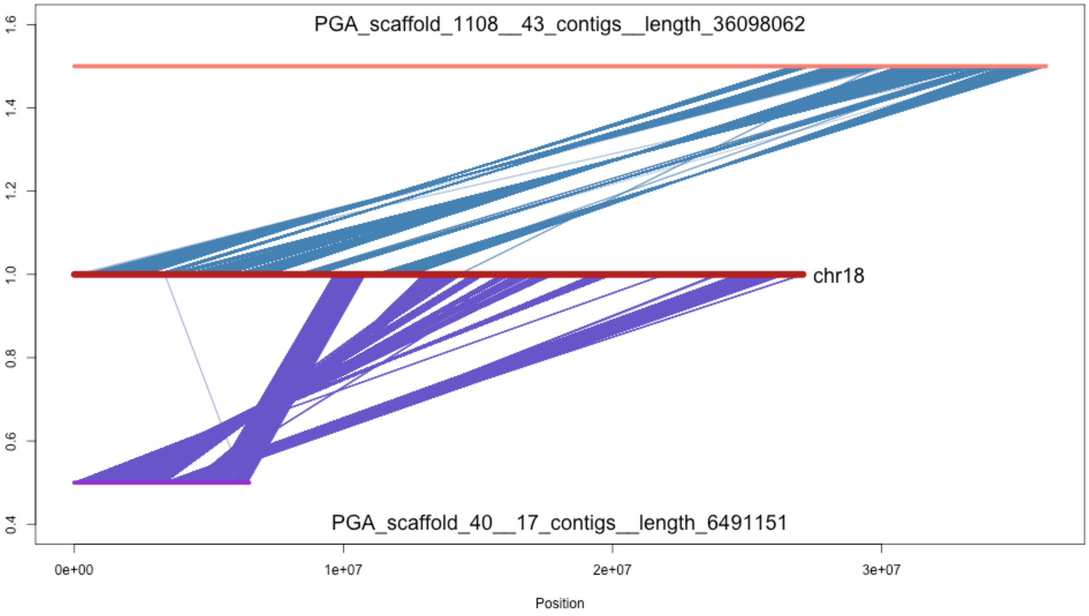
PGA_scaffold_1108__43_contigs__length_36098062
chr18
PGA_scaffold_40__17_contigs__length_6491151
1.6
1.4
1.2
1.0
0.8
0.6
0.4
0e+00
1e+07
2e+07
3e+07
Position

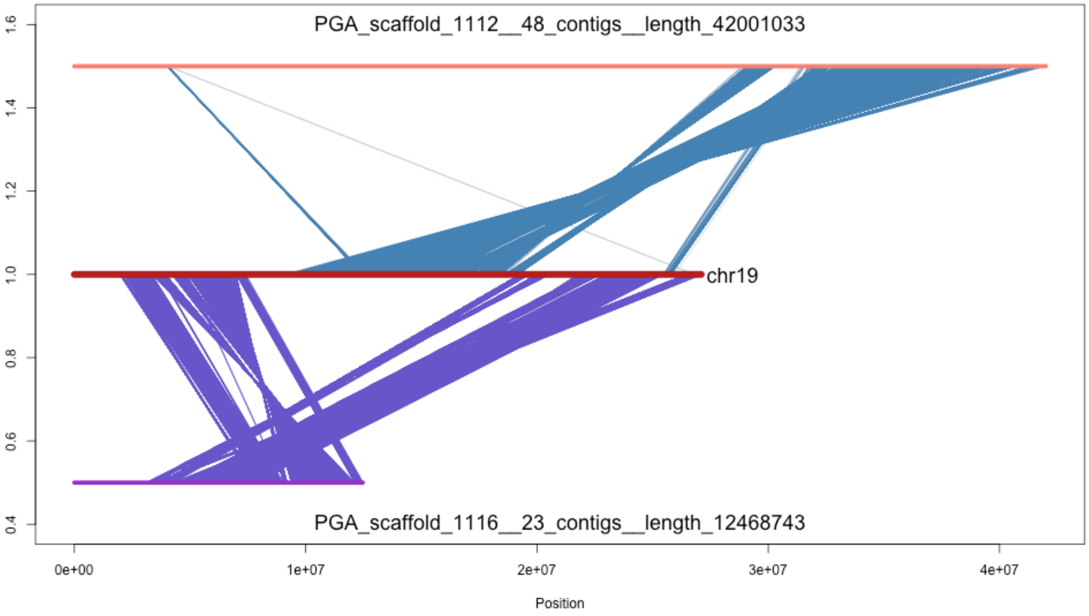

PGA_scaffold_1112__48_contigs__length_42001033
chr19
PGA_scaffold_1116__23_contigs__length_12468743
1.6
1.4
1.2
1.0
0.8
0.6
0.4
0e+00
1e+07
2e+07
3e+07
4e+07
Position

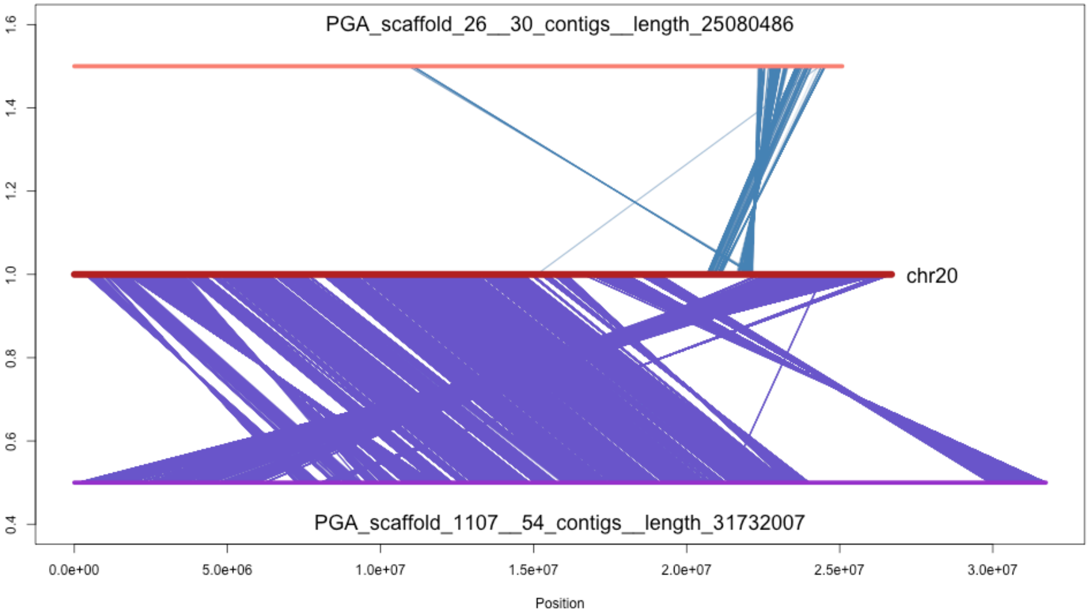
PGA_scaffold_26__30_contigs__length_25080486
chr20
PGA_scaffold_1107__54_contigs__length_31732007
1.6
1.4
1.2
1.0
0.8
0.6
0.4
0.0e+00
5.0e+06
1.0e+07
1.5e+07
2.0e+07
2.5e+07
3.0e+07
Position

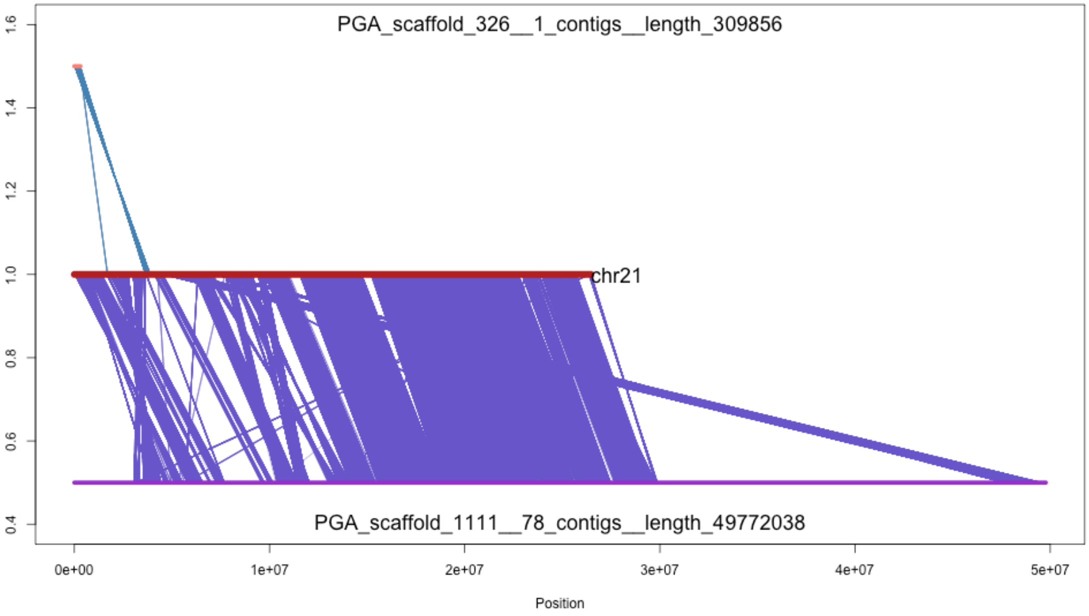
PGA_scaffold_326__1_contigs__length_309856
chr21
PGA_scaffold_1111__78_contigs__length_49772038
Position
0e+00
1e+07
2e+07
3e+07
4e+07
5e+07
0.4
0.6
0.8
1.0
1.2
1.4
1.6

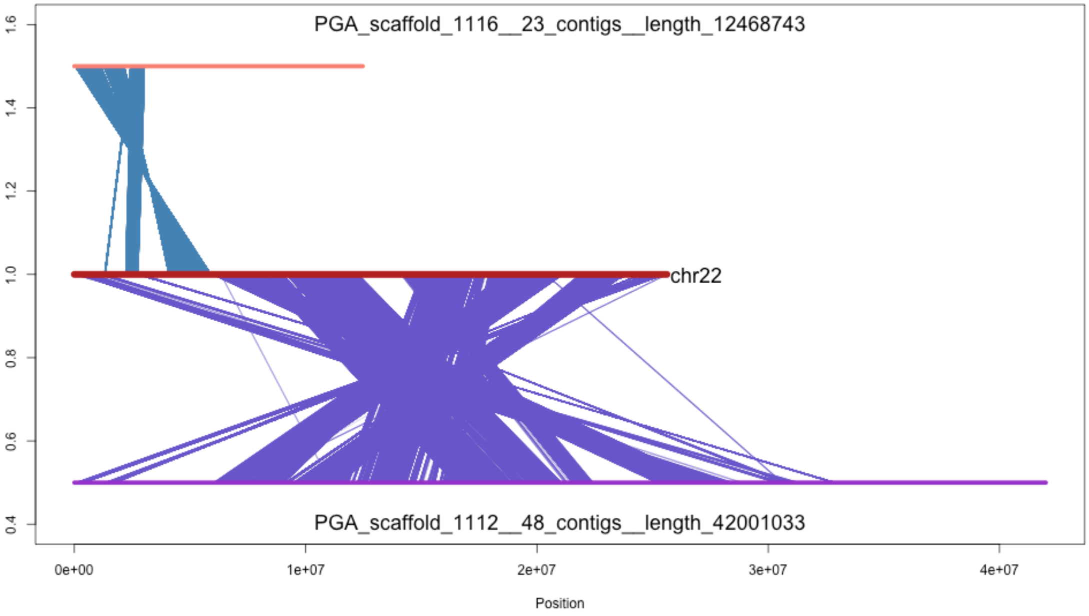
PGA_scaffold_1116__23_contigs__length_12468743
chr22
PGA_scaffold_1112__48_contigs__length_42001033
0.4
0.6
0.8
1.0
1.2
1.4
1.6
0e+00
1e+07
2e+07
3e+07
4e+07
Position

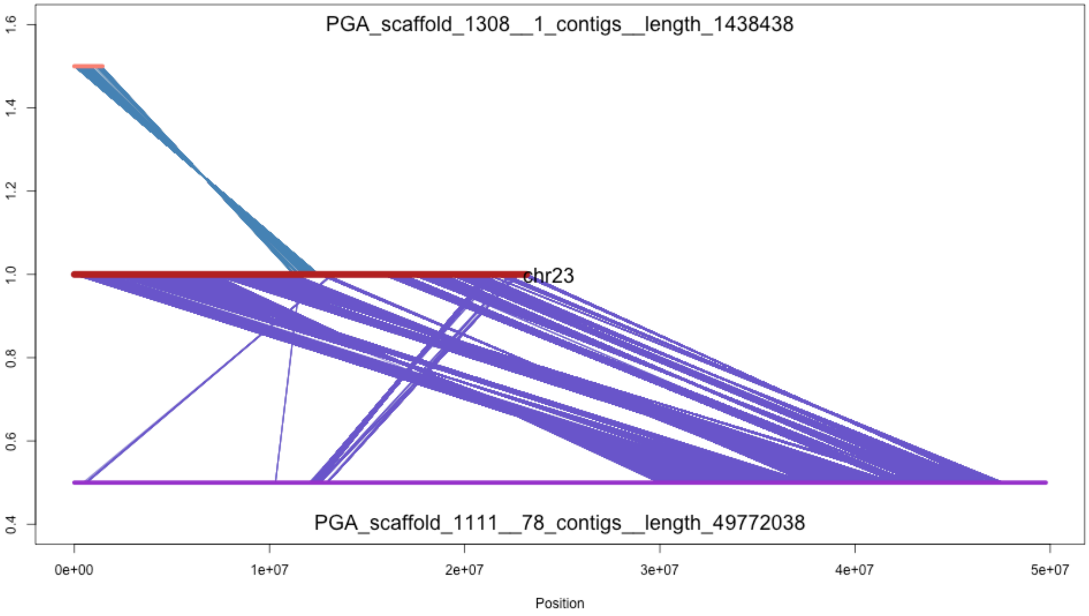
PGA_scaffold_1308__1_contigs__length_1438438
chr23
PGA_scaffold_1111__78_contigs__length_49772038
1.6
1.4
1.2
1.0
0.8
0.6
0.4
0e+00
1e+07
2e+07
3e+07
4e+07
5e+07
Position

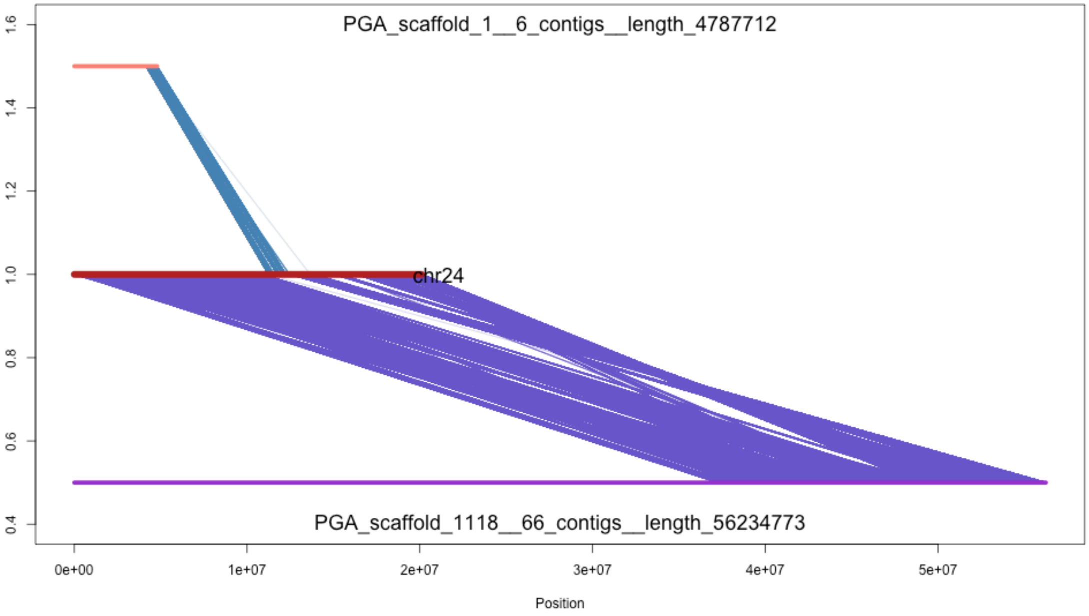

PGA_scaffold_1__6_contigs__length_4787712
chr24
PGA_scaffold_1118__66_contigs__length_56234773
1.6
1.4
1.2
1.0
0.8
0.6
0.4
0e+00
1e+07
2e+07
3e+07
4e+07
5e+07
Position

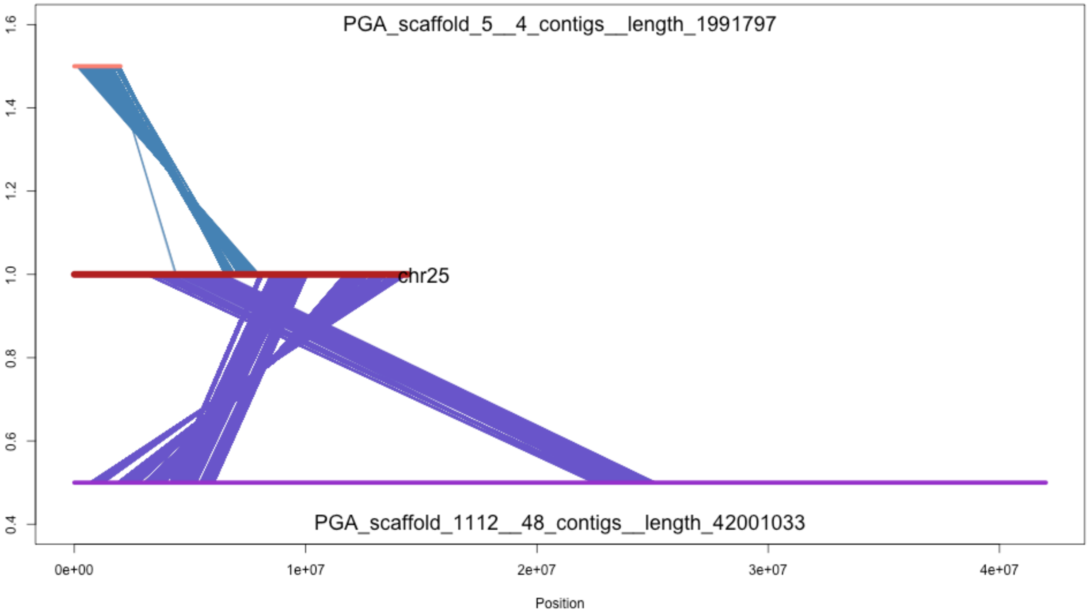
PGA_scaffold_5__4_contigs__length_1991797
chr25
PGA_scaffold_1112__48_contigs__length_42001033
0.4
0.6
0.8
1.0
1.2
1.4
1.6
0e+00
1e+07
2e+07
3e+07
4e+07
Position

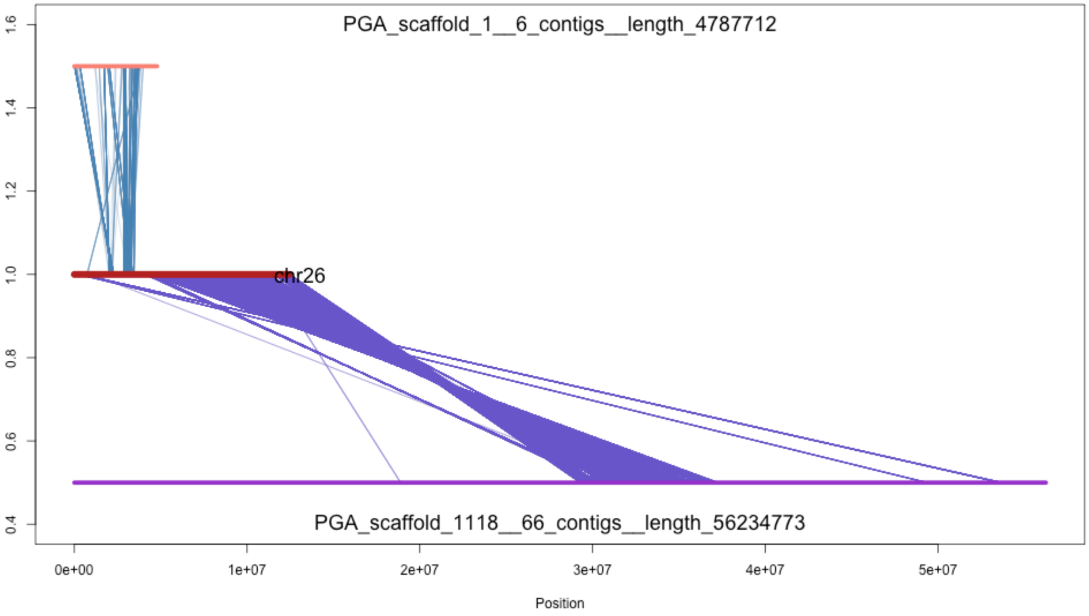
PGA_scaffold_1__6_contigs__length_4787712
chr26
PGA_scaffold_1118__66_contigs__length_56234773
1.6
1.4
1.2
1.0
0.8
0.6
0.4
0e+00
1e+07
2e+07
3e+07
4e+07
5e+07
Position
