## Supplementary figure 2b for "Limited parallelism in genetic adaptation to brackish water bodies in European sprat and Atlantic herring"

Supplementary Fig. 2. Cross-mapping between major sprat scaffolds and C. harengus chromosomes. a) SNP liftover from sprat scaffolds to the two most represented herring chromosomes. b) as in (a), but in opposite direction.

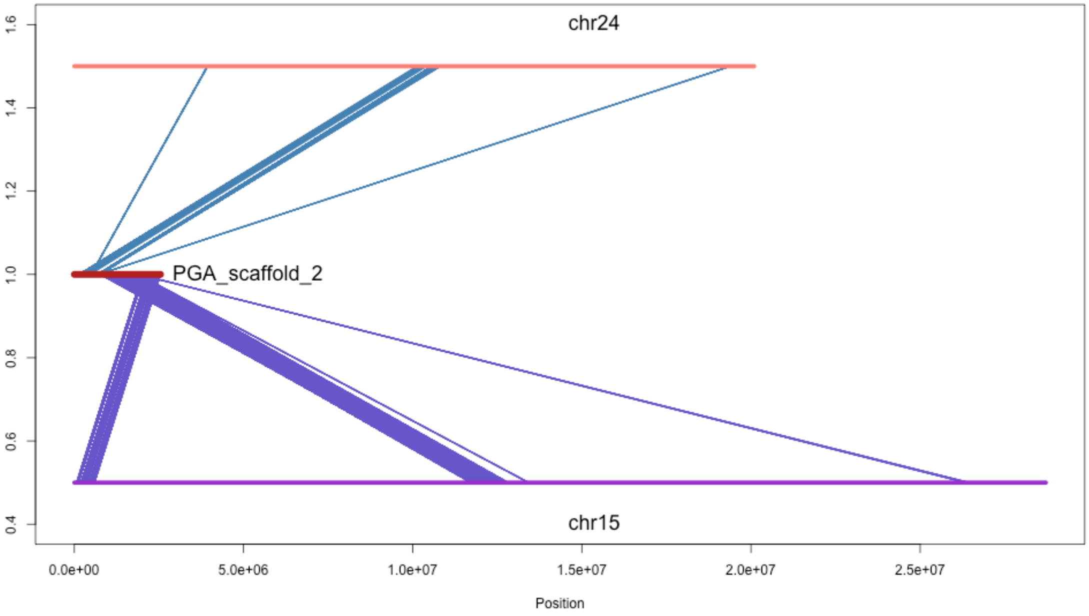

chr24
PGA_scaffold_2
chr15
1.6
1.4
1.2
1.0
0.8
0.6
0.4
0.0e+00
5.0e+06
1.0e+07
1.5e+07
2.0e+07
2.5e+07
Position

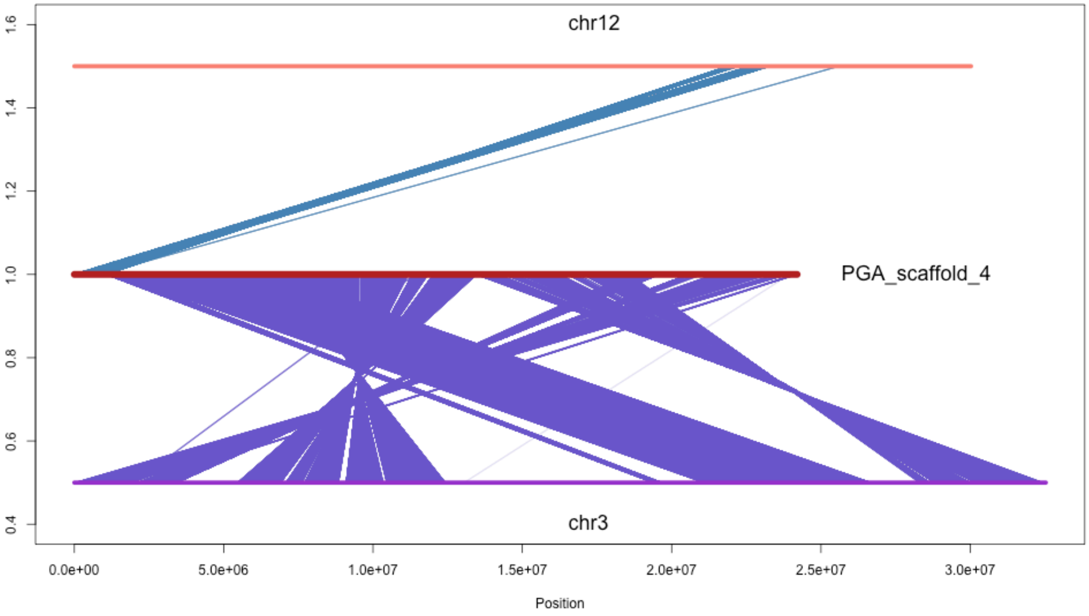
chr12
PGA_scaffold_4
chr3
1.6
1.4
1.2
1.0
0.8
0.6
0.4
0.0e+00
5.0e+06
1.0e+07
1.5e+07
2.0e+07
2.5e+07
3.0e+07
Position

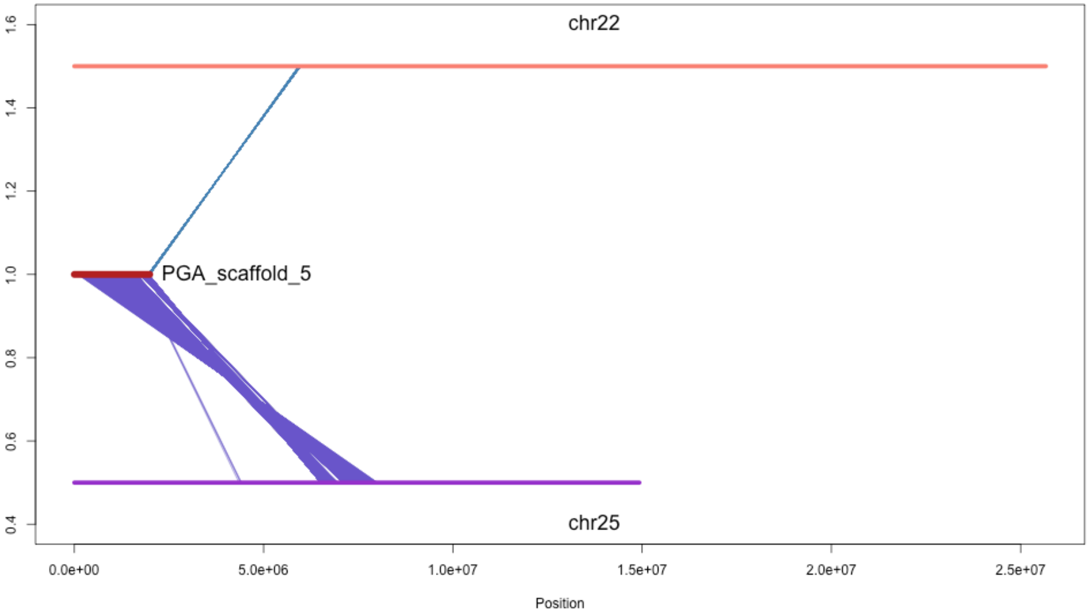
chr22
PGA_scaffold_5
chr25
0.4
0.6
0.8
1.0
1.2
1.4
1.6
0.0e+00
5.0e+06
1.0e+07
1.5e+07
2.0e+07
2.5e+07
Position

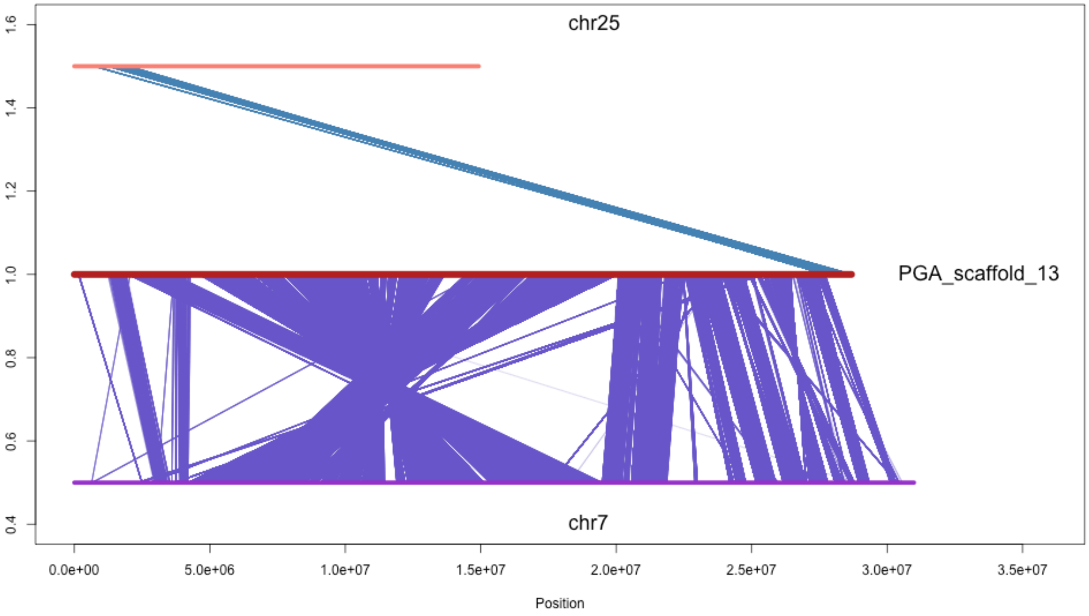
chr25
PGA_scaffold_13
chr7
1.6
1.4
1.2
1.0
0.8
0.6
0.4
0.0e+00
5.0e+06
1.0e+07
1.5e+07
2.0e+07
2.5e+07
3.0e+07
3.5e+07
Position

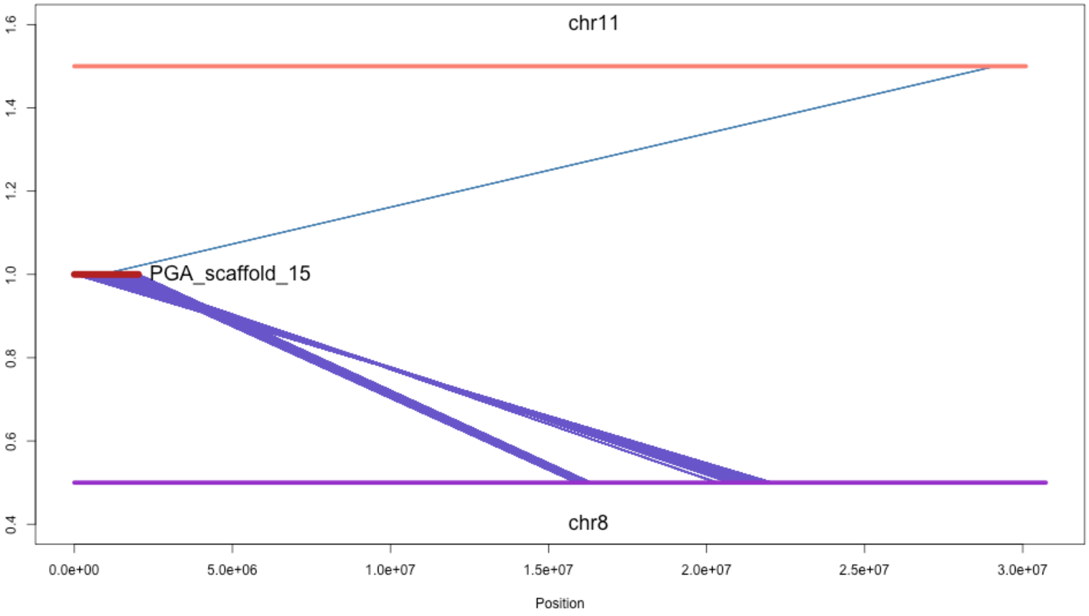
chr11
PGA_scaffold_15
chr8
1.6
1.4
1.2
1.0
0.8
0.6
0.4
0.0e+00
5.0e+06
1.0e+07
1.5e+07
2.0e+07
2.5e+07
3.0e+07
Position

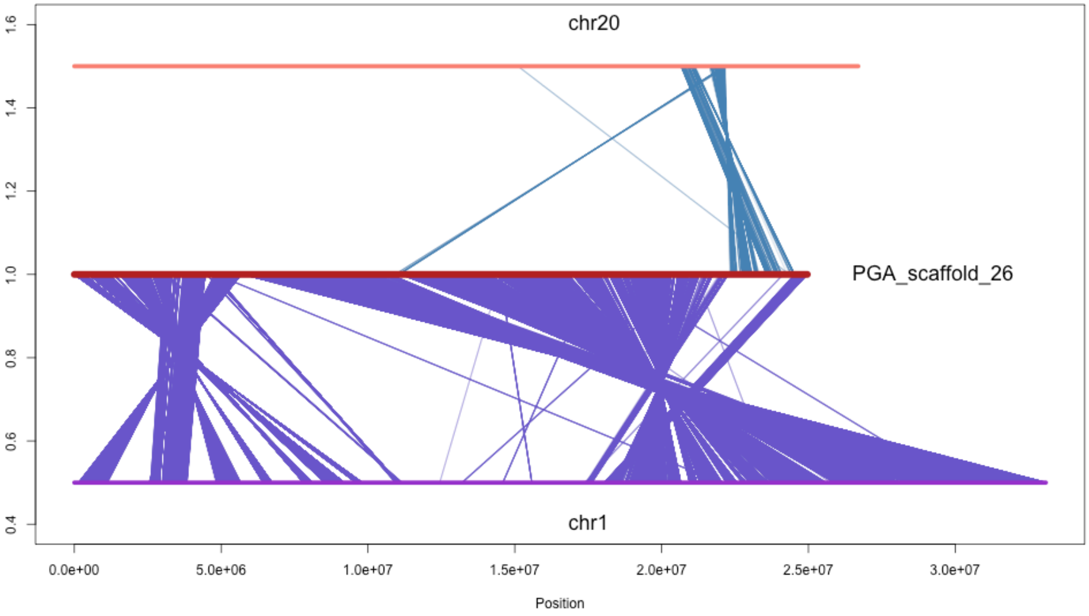
chr20
PGA_scaffold_26
chr1
1.6
1.4
1.2
1.0
0.8
0.6
0.4
0.0e+00
5.0e+06
1.0e+07
1.5e+07
2.0e+07
2.5e+07
3.0e+07
Position

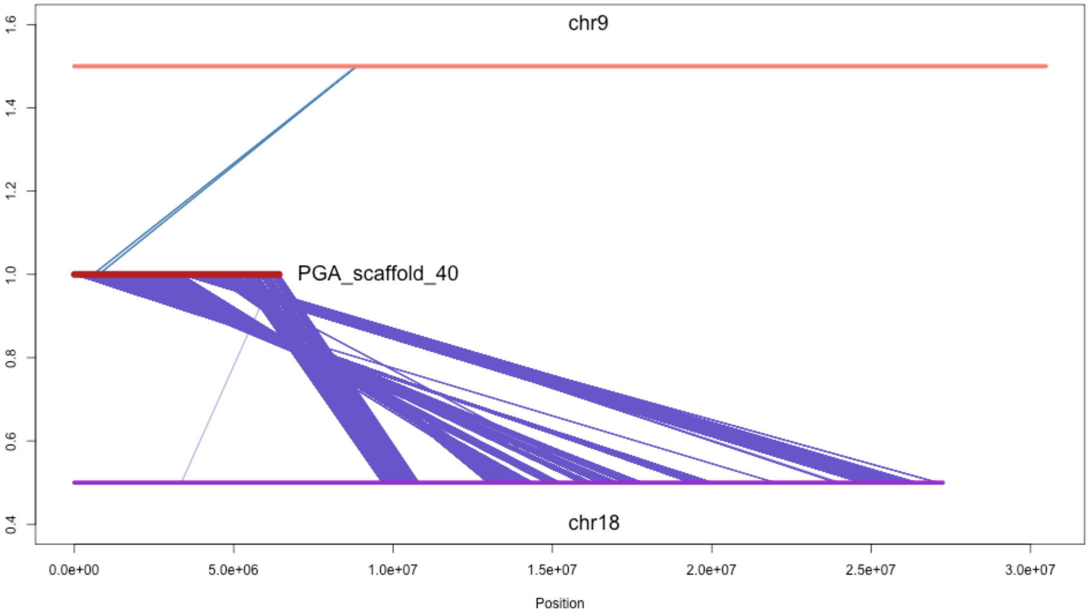

chr9
PGA_scaffold_40
chr18
1.6
1.4
1.2
1.0
0.8
0.6
0.4
0.0e+00
5.0e+06
1.0e+07
1.5e+07
2.0e+07
2.5e+07
3.0e+07
Position

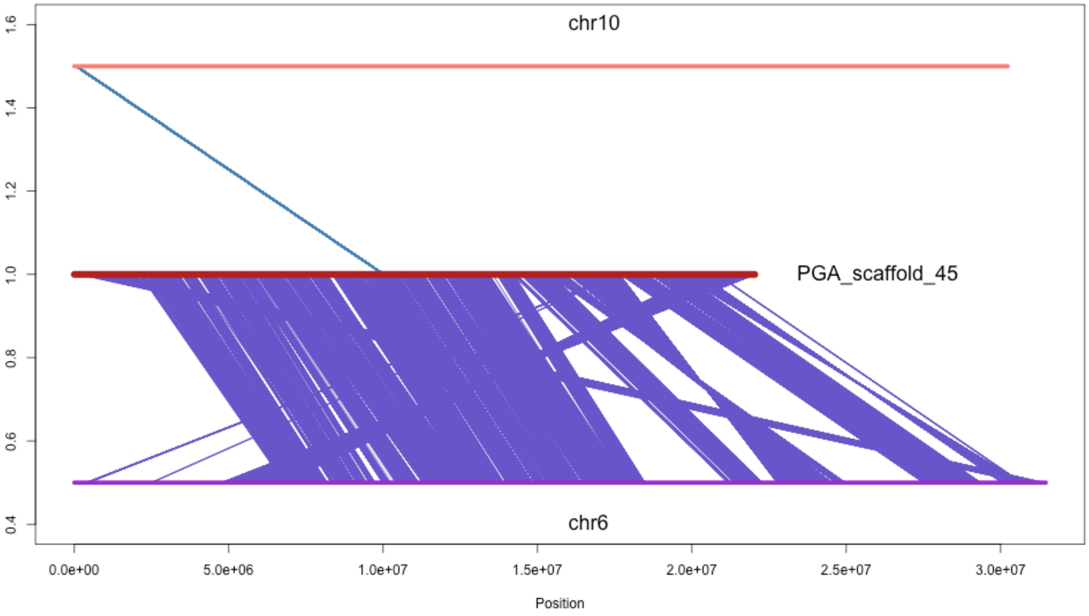
chr10
PGA_scaffold_45
chr6
1.6
1.4
1.2
1.0
0.8
0.6
0.4
0.0e+00
5.0e+06
1.0e+07
1.5e+07
2.0e+07
2.5e+07
3.0e+07
Position

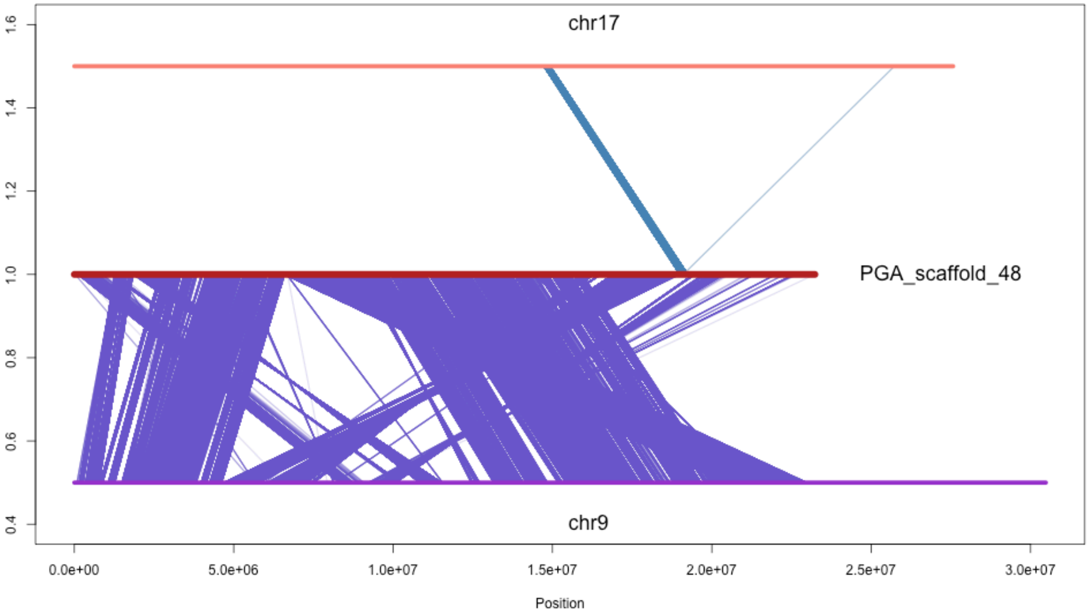
chr17
PGA_scaffold_48
chr9
1.6
1.4
1.2
1.0
0.8
0.6
0.4
0.0e+00
5.0e+06
1.0e+07
1.5e+07
2.0e+07
2.5e+07
3.0e+07
Position

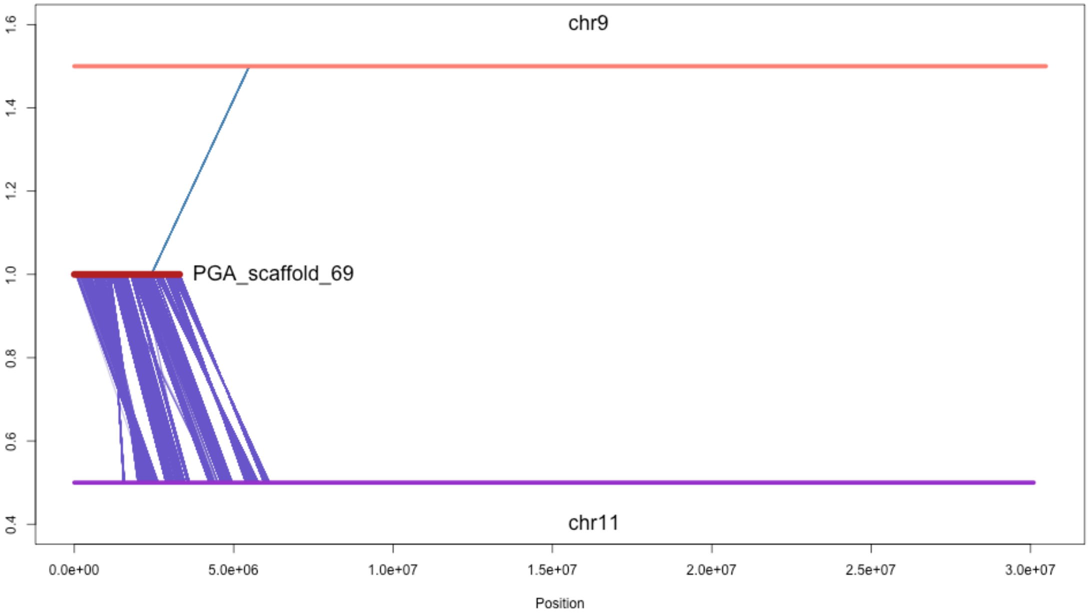

chr9
PGA_scaffold_69
chr11
1.6
1.4
1.2
1.0
0.8
0.6
0.4
0.0e+00
5.0e+06
1.0e+07
1.5e+07
2.0e+07
2.5e+07
3.0e+07
Position

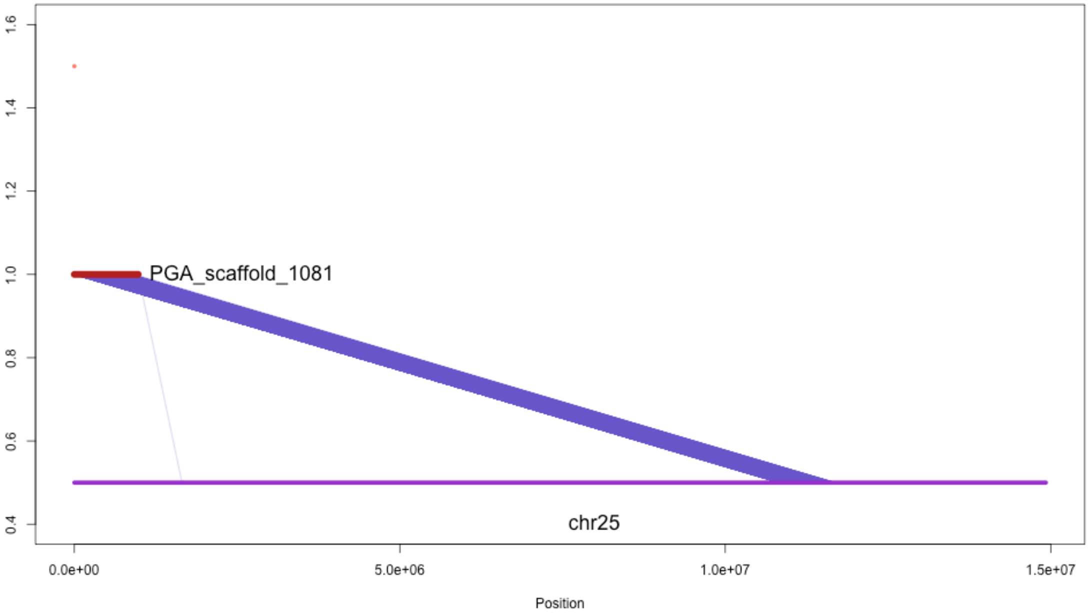

PGA_scaffold_1081
chr25
1.6
1.4
1.2
1.0
0.8
0.6
0.4
0.0e+00
5.0e+06
1.0e+07
1.5e+07
Position

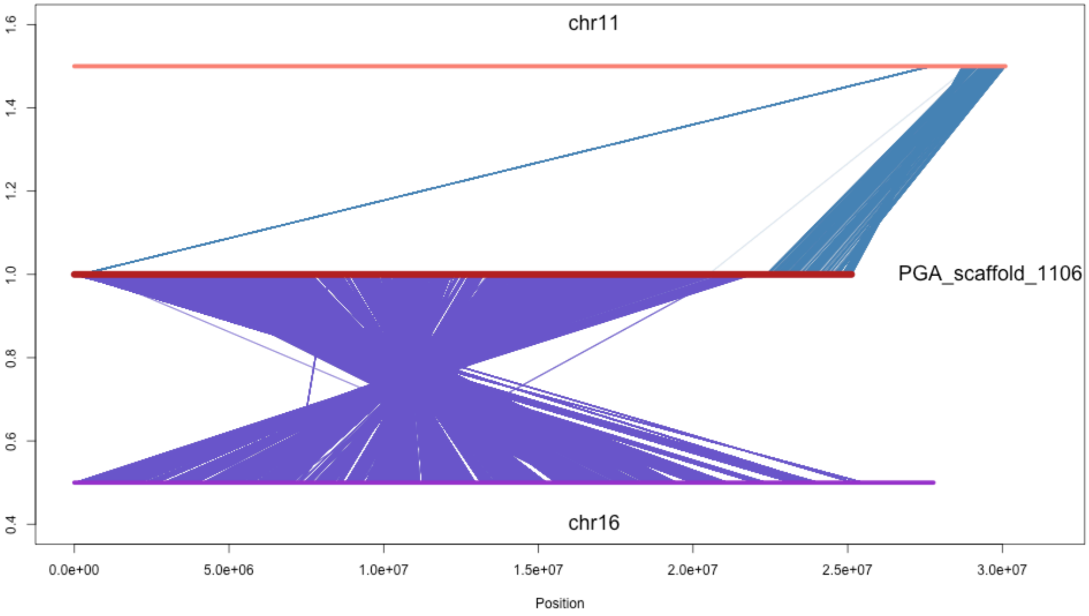

chr11
PGA_scaffold_1106
chr16
1.6
1.4
1.2
1.0
0.8
0.6
0.4
0.0e+00
5.0e+06
1.0e+07
1.5e+07
2.0e+07
2.5e+07
3.0e+07
Position

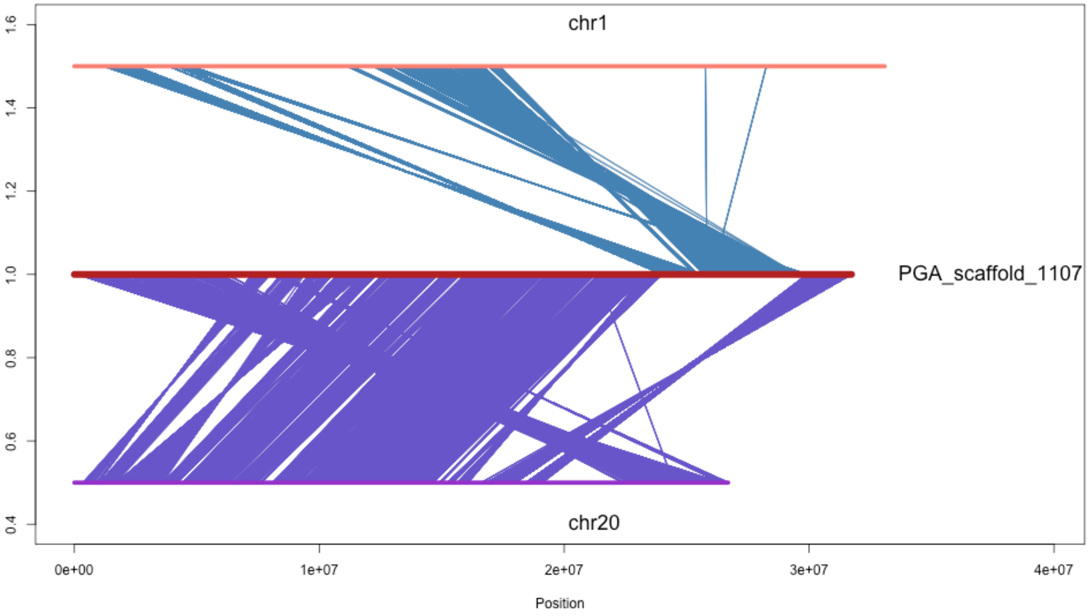
chr1
PGA_scaffold_1107
chr20
1.6
1.4
1.2
1.0
0.8
0.6
0.4
0e+00
1e+07
2e+07
3e+07
4e+07
Position

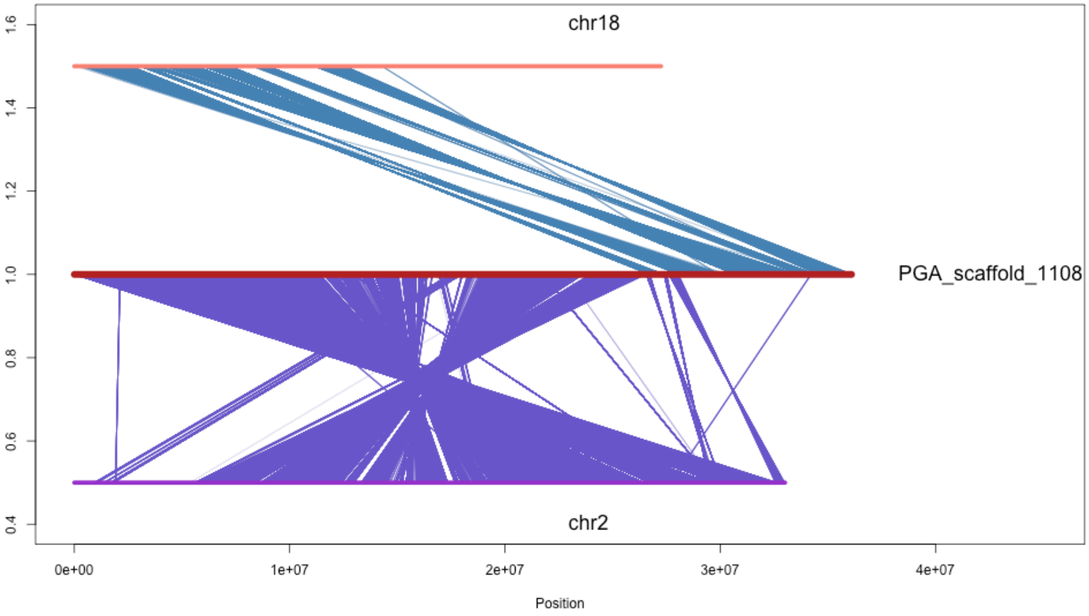
chr18
PGA_scaffold_1108
chr2
1.6
1.4
1.2
1.0
0.8
0.6
0.4
0e+00
1e+07
2e+07
3e+07
4e+07
Position

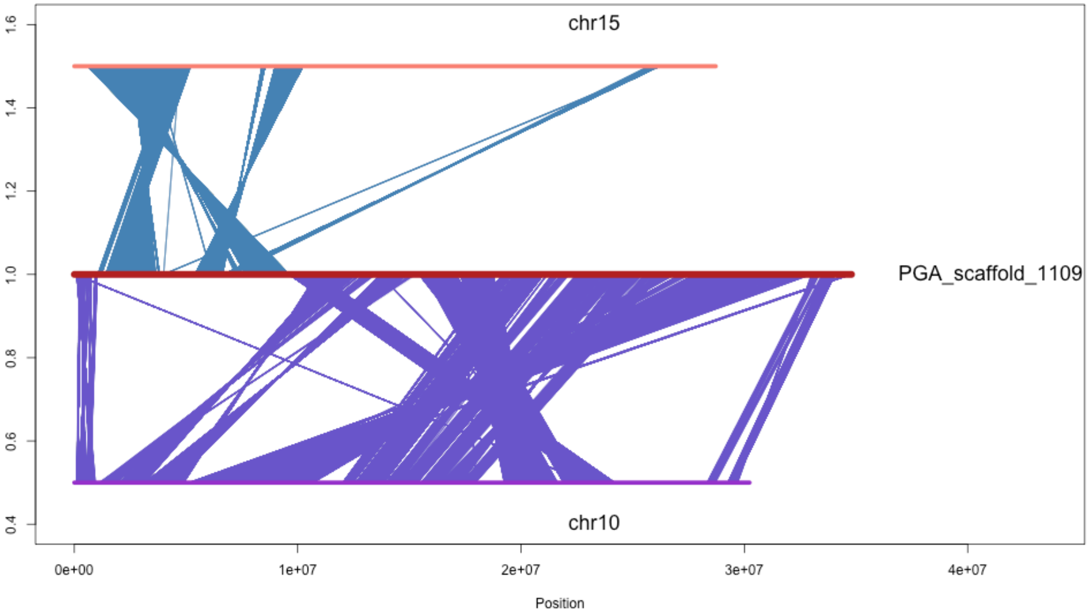
chr15
PGA_scaffold_1109
chr10
Position
0e+00
1e+07
2e+07
3e+07
4e+07
0.4
0.6
0.8
1.0
1.2
1.4
1.6

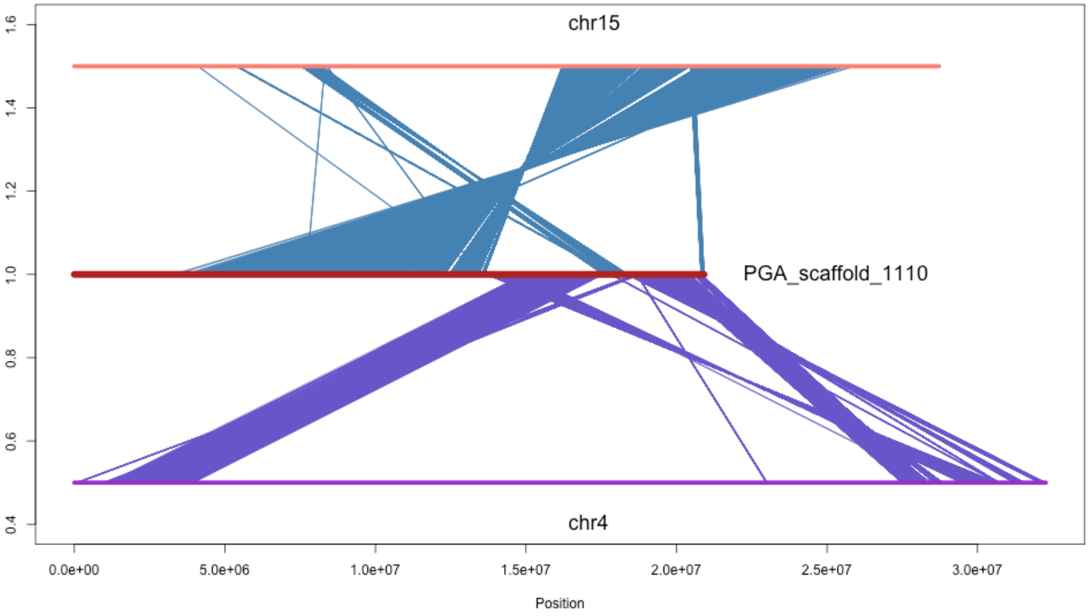

chr15
PGA_scaffold_1110
chr4
1.6
1.4
1.2
1.0
0.8
0.6
0.4
0.0e+00
5.0e+06
1.0e+07
1.5e+07
2.0e+07
2.5e+07
3.0e+07
Position

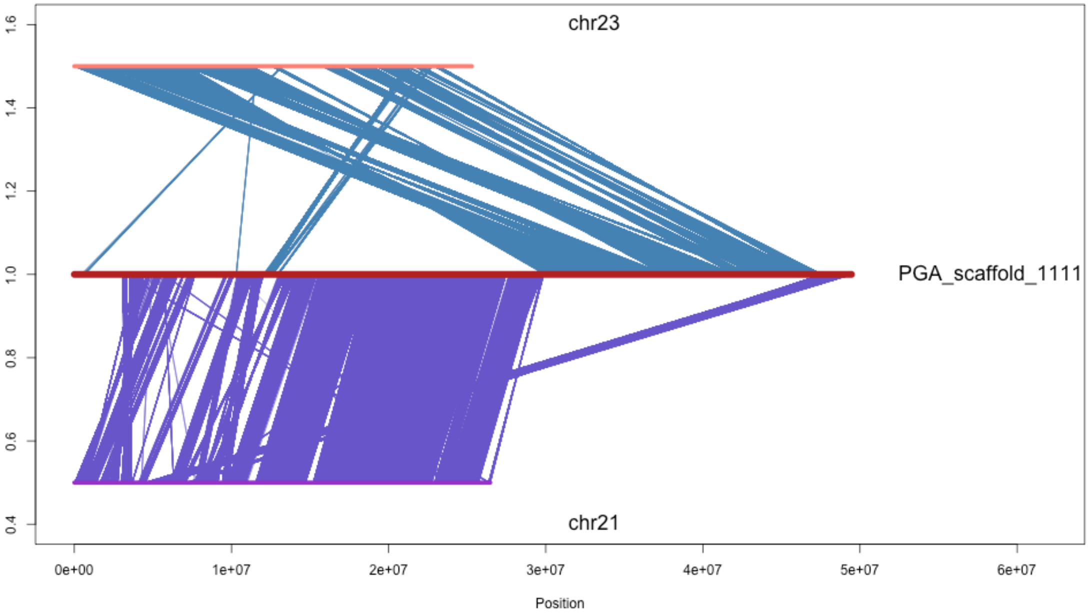
chr23
PGA_scaffold_1111
chr21
1.6
1.4
1.2
1.0
0.8
0.6
0.4
0e+00
1e+07
2e+07
3e+07
4e+07
5e+07
6e+07
Position

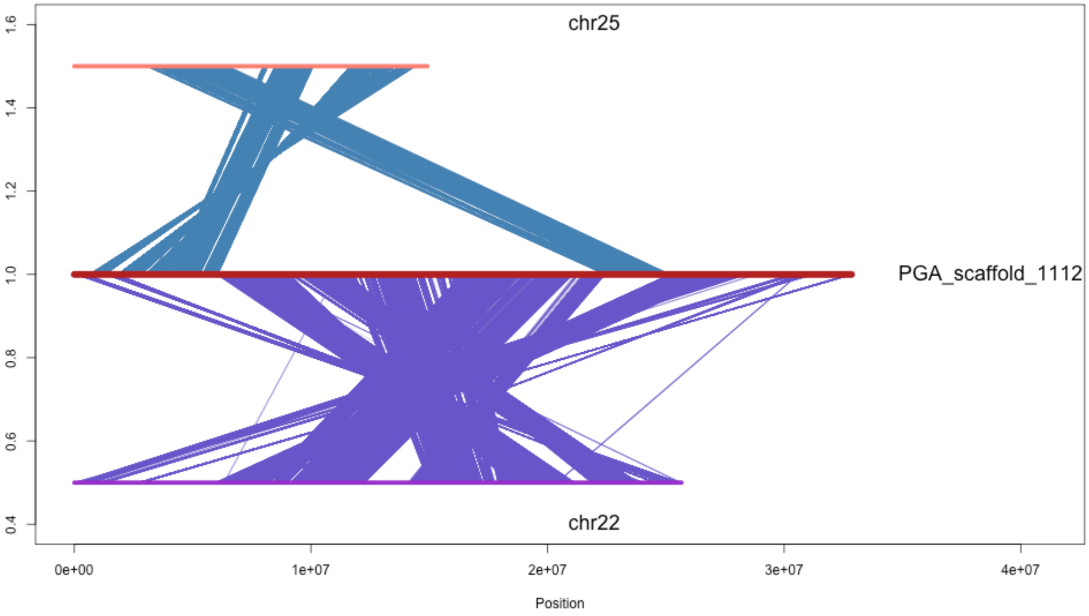
chr25
PGA_scaffold_1112
chr22
1.6
1.4
1.2
1.0
0.8
0.6
0.4
0e+00
1e+07
2e+07
3e+07
4e+07
Position

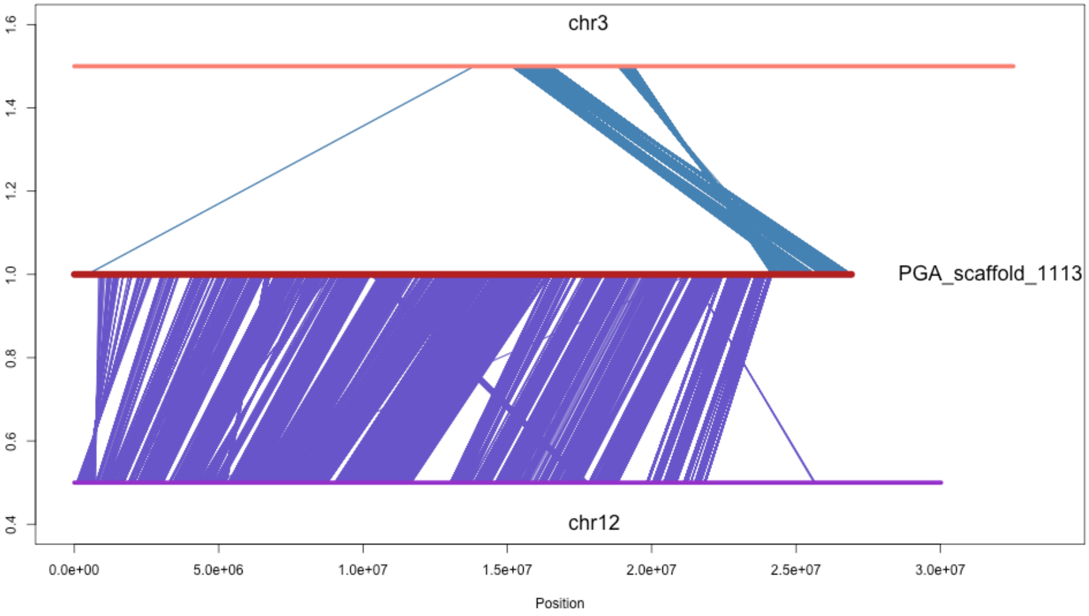

chr3
PGA_scaffold_1113
chr12
1.6
1.4
1.2
1.0
0.8
0.6
0.4
0.0e+00
5.0e+06
1.0e+07
1.5e+07
2.0e+07
2.5e+07
3.0e+07
Position

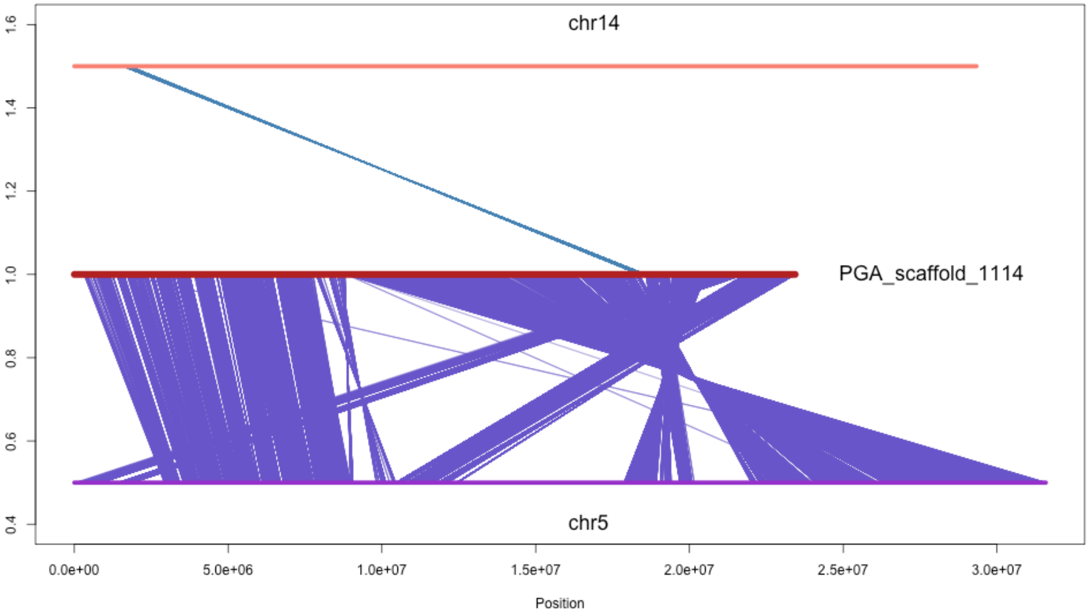
chr14
PGA_scaffold_1114
chr5
1.6
1.4
1.2
1.0
0.8
0.6
0.4
0.0e+00
5.0e+06
1.0e+07
1.5e+07
2.0e+07
2.5e+07
3.0e+07
Position

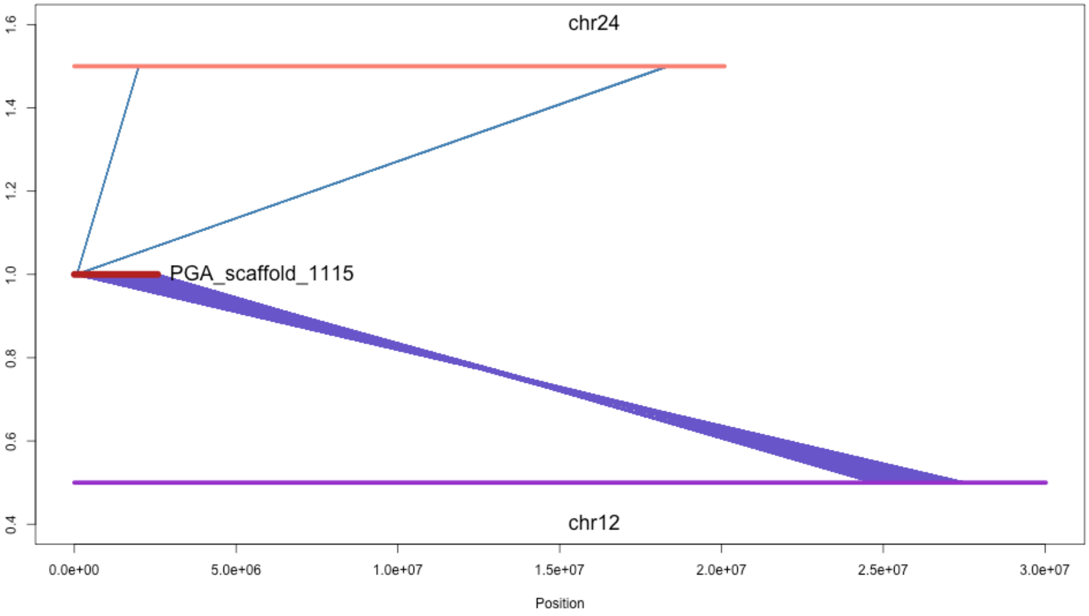
chr24
PGA_scaffold_1115
chr12
1.6
1.4
1.2
1.0
0.8
0.6
0.4
0.0e+00
5.0e+06
1.0e+07
1.5e+07
2.0e+07
2.5e+07
3.0e+07
Position

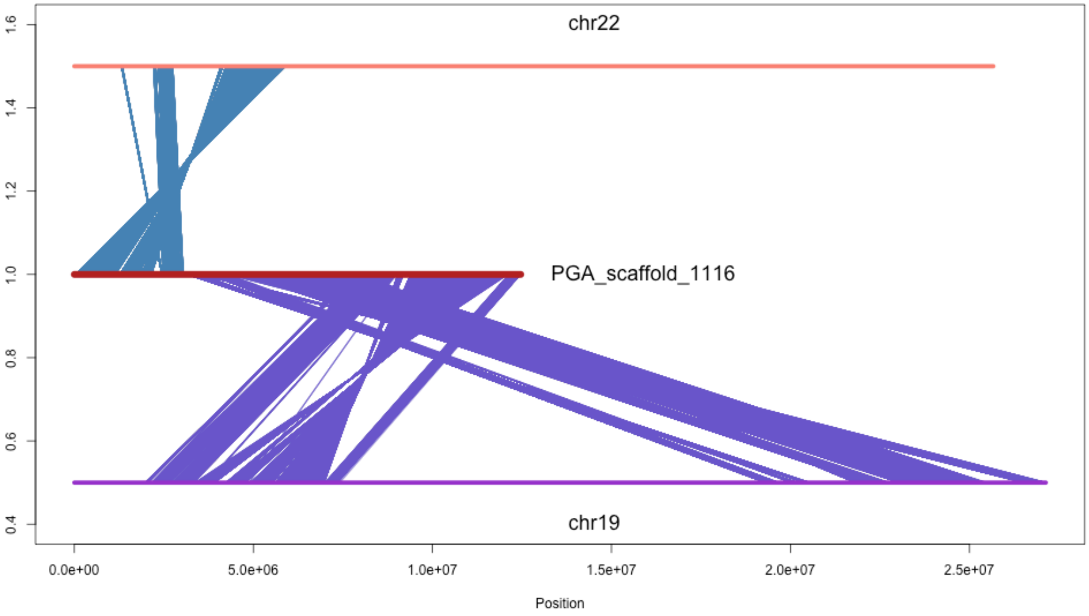
chr22
PGA_scaffold_1116
chr19
1.6
1.4
1.2
1.0
0.8
0.6
0.4
0.0e+00
5.0e+06
1.0e+07
1.5e+07
2.0e+07
2.5e+07
Position

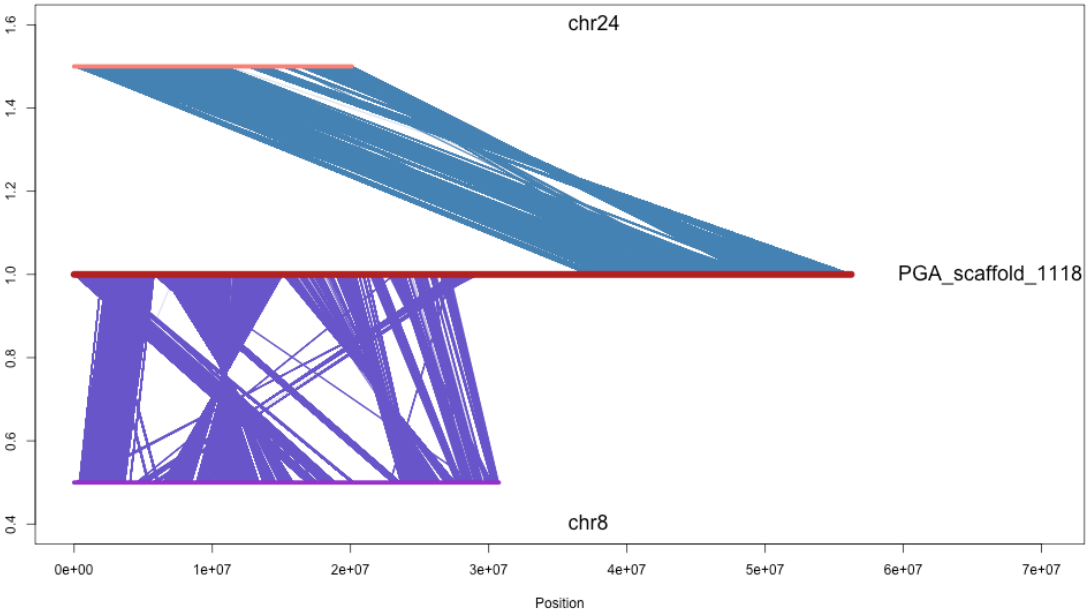
chr24
PGA_scaffold_1118
chr8
1.6
1.4
1.2
1.0
0.8
0.6
0.4
0e+00
1e+07
2e+07
3e+07
4e+07
5e+07
6e+07
7e+07
Position

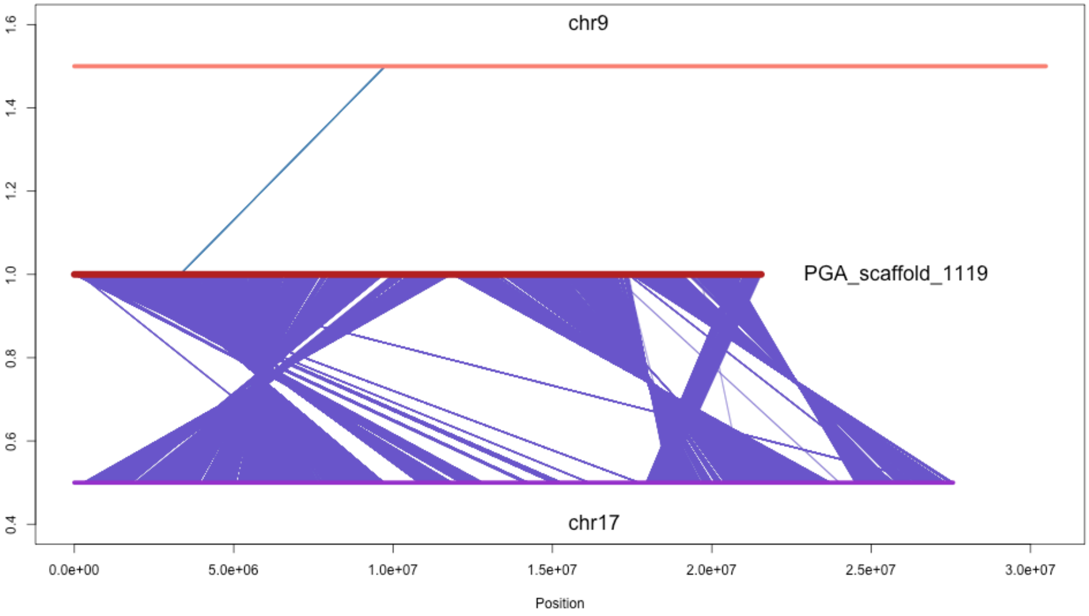

chr9
PGA_scaffold_1119
chr17
1.6
1.4
1.2
1.0
0.8
0.6
0.4
0.0e+00
5.0e+06
1.0e+07
1.5e+07
2.0e+07
2.5e+07
3.0e+07
Position

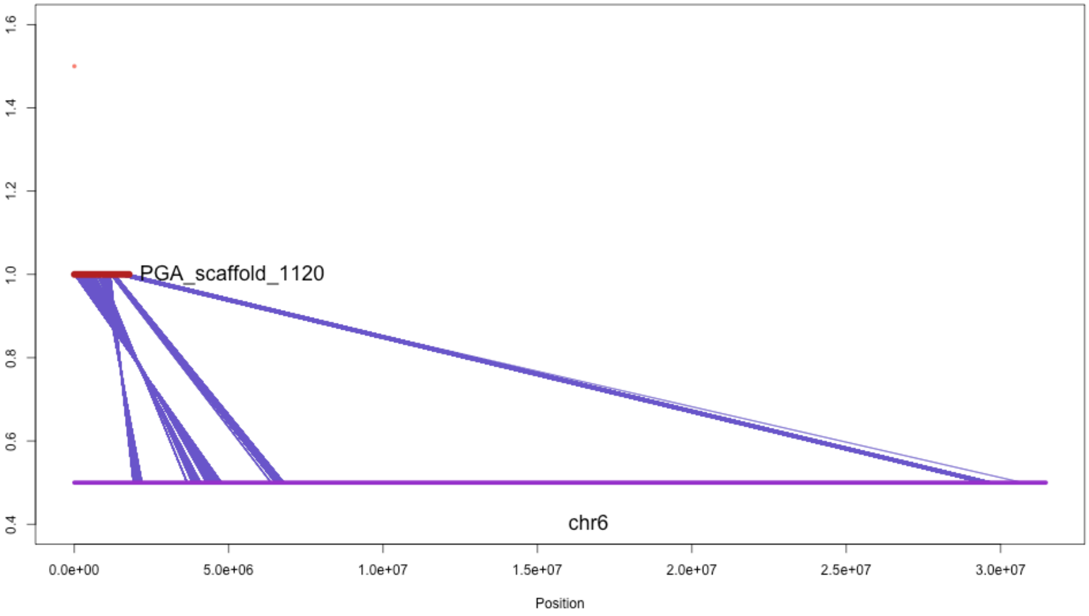

PGA_scaffold_1120
chr6
Position
0.0e+00
5.0e+06
1.0e+07
1.5e+07
2.0e+07
2.5e+07
3.0e+07
0.4
0.6
0.8
1.0
1.2
1.4
1.6

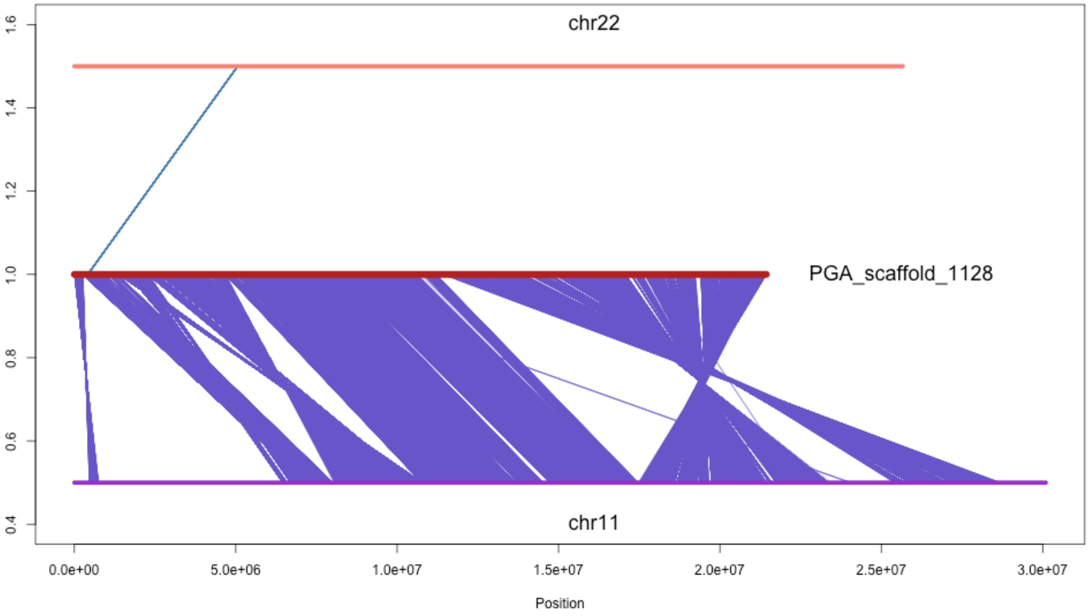

chr22
PGA_scaffold_1128
chr11
1.6
1.4
1.2
1.0
0.8
0.6
0.4
0.0e+00
5.0e+06
1.0e+07
1.5e+07
2.0e+07
2.5e+07
3.0e+07
Position

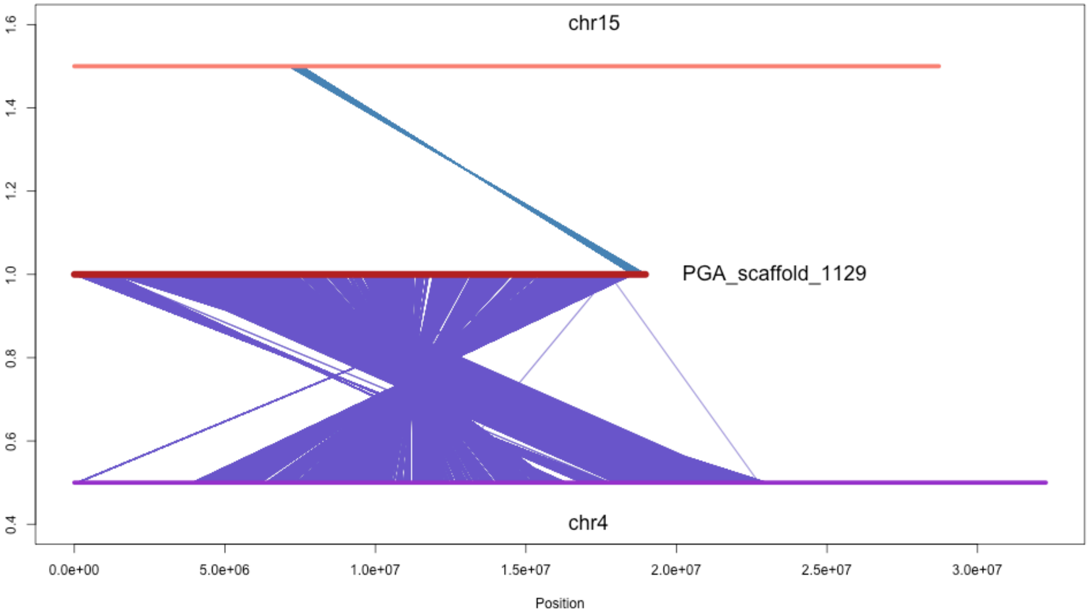
chr15
PGA_scaffold_1129
chr4
1.6
1.4
1.2
1.0
0.8
0.6
0.4
0.0e+00
5.0e+06
1.0e+07
1.5e+07
2.0e+07
2.5e+07
3.0e+07
Position

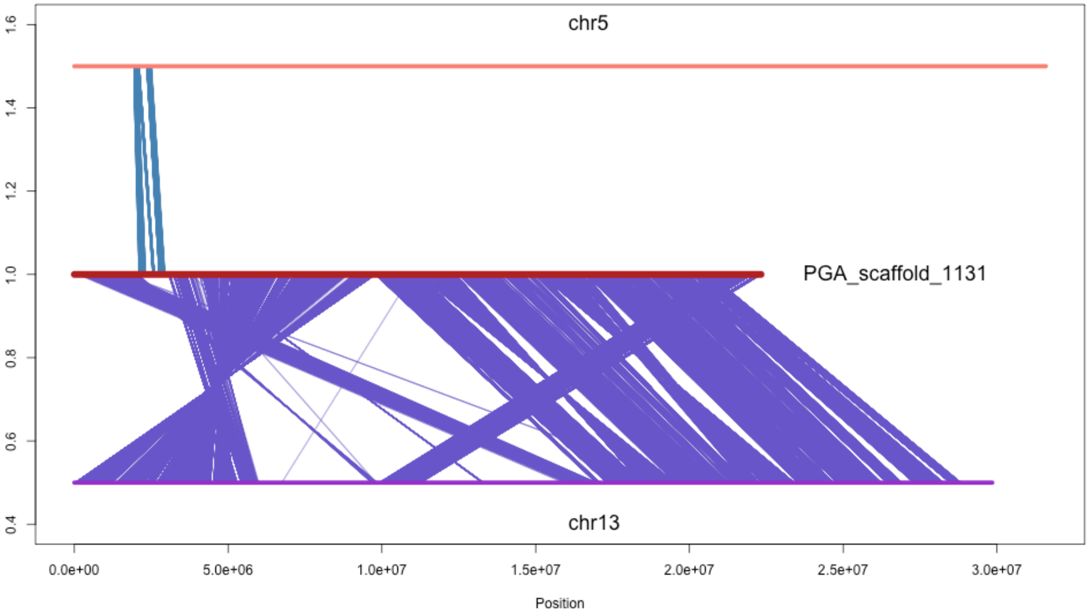

chr5
PGA_scaffold_1131
chr13
1.6
1.4
1.2
1.0
0.8
0.6
0.4
0.0e+00
5.0e+06
1.0e+07
1.5e+07
2.0e+07
2.5e+07
3.0e+07
Position

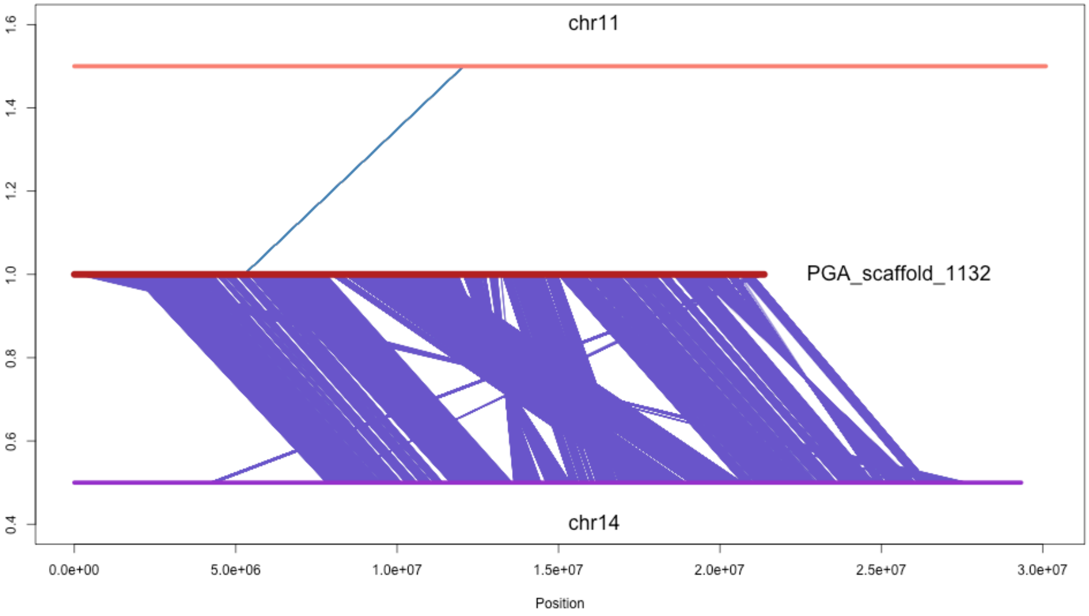
chr11
PGA_scaffold_1132
chr14
1.6
1.4
1.2
1.0
0.8
0.6
0.4
0.0e+00
5.0e+06
1.0e+07
1.5e+07
2.0e+07
2.5e+07
3.0e+07
Position
